# Targeting of TP53-independent cell cycle checkpoints overcomes FOLFOX resistance in Metastatic Colorectal Cancer

**DOI:** 10.1101/2021.02.04.429849

**Authors:** Corina Behrenbruch, Momeneh Foroutan, Phoebe Lind, Jai Smith, Mélodie Grandin, Benjamin Cooper, Carolyn Shembrey, Susanne Ramm, Karla Cowley, Iva Nikolic, Jennii Luu, Joseph Cursons, Rosie Millen, Ann-Marie Patch, Nicholas D. Huntington, Alain Puisieux, Michael Michael, Brett Knowles, Benjamin NJ. Thomson, Robert G. Ramsay, Sean M. Grimmond, Kaylene Simpson, Alexander G. Heriot, Frédéric Hollande

**Affiliations:** The University of Melbourne, Department of Clinical Pathology, Victorian Comprehensive Cancer Centre, Melbourne, VIC3000, Australia; University of Melbourne Centre for Cancer Research, Victorian Comprehensive Cancer Centre, Melbourne VIC3000, Australia; The University of Melbourne, Sir Peter MacCallum Department of Oncology, Victorian Comprehensive Cancer Centre, Melbourne, VIC3000, Australia; Department of General Surgical Specialties, The Royal Melbourne Hospital, Parkville, VIC 3050, Australia; Biomedicine Discovery Institute, Monash University, VIC, 3800, Australia; The University of Melbourne, Department of Pharmacology and Therapeutics, Medical Building, Parkville VIC3010, Australia; Victorian Centre for Functional Genomics, Peter MacCallum Cancer Centre, Melbourne, VIC3000, Australia; Peter MacCallum Cancer Centre, Melbourne, VIC3000, Australia; QIMR, 300 Herston Rd, Herston QLD 4006, Australia; Department of Medical Oncology, Peter MacCallum Cancer Centre, Victorian Comprehensive Cancer Centre, Melbourne, VIC3000, Australia; The University of Melbourne, Royal Melbourne Hospital Department of Surgery, Parkville, VIC3050, Australia; Institut Curie, 26 rue d’Ulm75248 Paris Cedex 05 - France; Department of Cancer Surgery, Peter MacCallum Cancer Centre, Victorian Comprehensive Cancer Centre, Melbourne, VIC3000, Australia; The University of Melbourne, Department of Surgery, St Vincent’s Hospital, Fitzroy, VIC3065, Australia

**Author notes:** These authors contributed equally to the present work (Corina Behrenbruch, Momeneh Foroutan). Current address: Biomedicine Discovery Institute, Monash University, VIC, 3800, Australia.

## Abstract

Patients with colorectal cancer (CRC) frequently develop liver metastases during the course of their disease. A substantial proportion of them receive neoadjuvant FOLFOX (5-Fluorouracil, Oxaliplatin, Leucovorin) prior to surgery in an attempt to enable successful surgical removal of their metastases and to reduce the risk of recurrence. Yet, the majority of patients progress during treatment or recur following surgery, and molecular mechanisms that contribute to FOLFOX resistance remain poorly understood. Here, using a combination of phenotypic, transcriptomic and genomic analyses of both tumor samples derived from patients with metastatic CRC and matching patient-derived tumor organoids (PDTOs), we characterize a novel FOLFOX resistance mechanism and identify inhibitors that target this mechanism to resensitize metastatic organoids to FOLFOX. Resistant PDTOs, identified after *in vitro* exposure to FOLFOX, exhibited elevated expression of E2F pathway, S phase, G_2_/M and spindle assembly checkpoints (SAC) genes. Similar molecular features were detected in CRLM from patients with progressive disease while under neoadjuvant FOLFOX treatment, highlighting the relevance of this finding. FOLFOX resistant PDTOs displayed inactivating mutations of TP53 and exhibited transcriptional features of P53 pathway downregulation. We found that they accumulated in early S-phase and underwent significant DNA damage during FOLFOX exposure, thereafter arresting in G_2_/M while they repaired their DNA after FOLFOX withdrawal. In parallel, results of a large kinase inhibitor screen indicated that drugs targeting regulators of the DNA damage response, G_2_M checkpoint and SAC had cytotoxic effects on PDTOs generated from patients whose disease progressed during treatment with FOLFOX. Corroborating this finding, CHK1 and WEE1 inhibitors were found to synergize with FOLFOX and sensitize previously resistant PDTOs. Additionally, targeting the SAC master regulator MPS1 using empesertib after exposure to FOLFOX, when cells accumulate in G_2_M, was also very effective to kill FOLFOX-resistant PDTOs. Our results indicate that targeted and timely inhibition of specific cell cycle checkpoints shows great potential to improve response rates to FOLFOX in patients with metastatic CRC, for whom therapeutic alternatives remain extremely limited.

## INTRODUCTION

At least 50% of patients with colorectal cancer (CRC) develop colorectal liver metastasis (CRLM) during the course of their disease^1^. In patients with CRLM, surgical resection currently offers the only chance of cure, improving 5-year survival rates from 13% with chemotherapy alone to 33-55% if all sites of disease can be resected^2–4^. However, most patients are not eligible for upfront surgical resection and will instead receive chemotherapeutic intervention as first line therapy. FOLFOX (5-fluorouracil, oxaliplatin, leucovorin) chemotherapy is one of the first line neoadjuvant regimens administered prior to surgical resection of CRLM^1,5^. Trials for both upfront and potentially resectable CRLM have shown that a significant number of patients progress during treatment with FOLFOX, and recurrence is frequent (> 50%) following surgical resection^6–9^. Additionally, oxaliplatin-induced cumulative, dose-dependent grade 3 peripheral neuropathy during treatment contributes to up to 30% of patients failing to complete the prescribed course of treatment, not to mention the effect that it has on their quality of life^10 11^. Despite its widespread use and toxicity, the mechanisms contributing to FOLFOX resistance in patients with metastatic colorectal cancer (mCRC) remain poorly understood. Improving our understanding of FOLFOX resistance could enable a better selection of patients that who might benefit from FOLFOX treatment, limit the exposure to its toxicity in those that will not benefit and/or allow the identification of combination regimens that may circumnavigate resistance.

To address this unmet need we combined phenotypic, transcriptomic and genomic analysis of tumor samples and matching patient derived tumor organoids (PDTOs) collected from patients with mCRC to identify a novel resistance signature that underpins CRLM resistance to FOLFOX chemotherapy. Resistant PDTOs were characterized by elevated expression of genes contributing to S-phase, G2/M and spindle assembly cell cycle checkpoints, and by reduced expression of TP53 pathway genes. They also displayed a two-step suspension of cell-cycle progression, first during exposure to FOLFOX, then during post-treatment recovery. Comparative genomic analysis of PDTO and matching tumor samples did not highlight any unique mutational profile that might drive this resistant phenotype. Additionally, a large drug screening assay using PDTOs generated from patients with short progression free survival following neoadjuvant

FOLFOX chemotherapy showed efficacy of inhibitors regulating DNA damage response, as well as the TP53 independent S phase, G_2_/M transition and mitotic checkpoints. Validation experiments demonstrated that FOLFOX-resistant PDTOs were very effectively killed by concomitant treatment with inhibitors of Checkpoint Kinase 1 (CHK1) or Wee1, or by sequential treatment with an inhibitor of the spindle assembly checkpoint master regulator MPS1. Collectively our results identify a novel mechanism of FOLFOX resistance in metastatic colorectal cancer and demonstrate that blocking the S-phase checkpoint during FOLFOX exposure or the G_2_/M block during the post-treatment DNA damage repair phase severely affects the survival ability of resistant PDTOs, thus highlighting several therapeutic opportunities for patients with CRLM.

## RESULTS

### Organoid Derivation, Molecular Characterization and Correlation with Patient Tumor Data

A total of 62 samples were collected from 38 patients with stage IV CRC, including 49 CRLM. Thirteen matched primary tumor samples were also collected from 9 patients (Details in **Suppl. Figure 1)**. Of the 38 patients from which samples were obtained, 20 (53%) had received neoadjuvant chemotherapy (neoCT) at the time of surgical resection, while 16/38 (47%) were chemotherapy naïve. Details of patient demographics, clinico-pathological details and specific chemotherapy regimens are provided in **Table 1 and Suppl. Table 1**. To allow investigation of resistance mechanisms *ex-vivo*, we generated PDTOs from chemo-naïve and post-neoadjuvant CRLM, as well as from primary tumors that matched a subset of the chemo-naïve CRLM. PDTOs are useful tools to predict drug sensitivity and/or identify potential therapeutic candidates^12,13^. A total of 24 PDTOs demonstrated sufficient growth for amplification and processing towards whole genome sequencing (WGS)/Whole Exome Sequencing (WES) and RNA sequencing (RNAseq) analyses **(Suppl. Figure 1**).

**Table 1.**
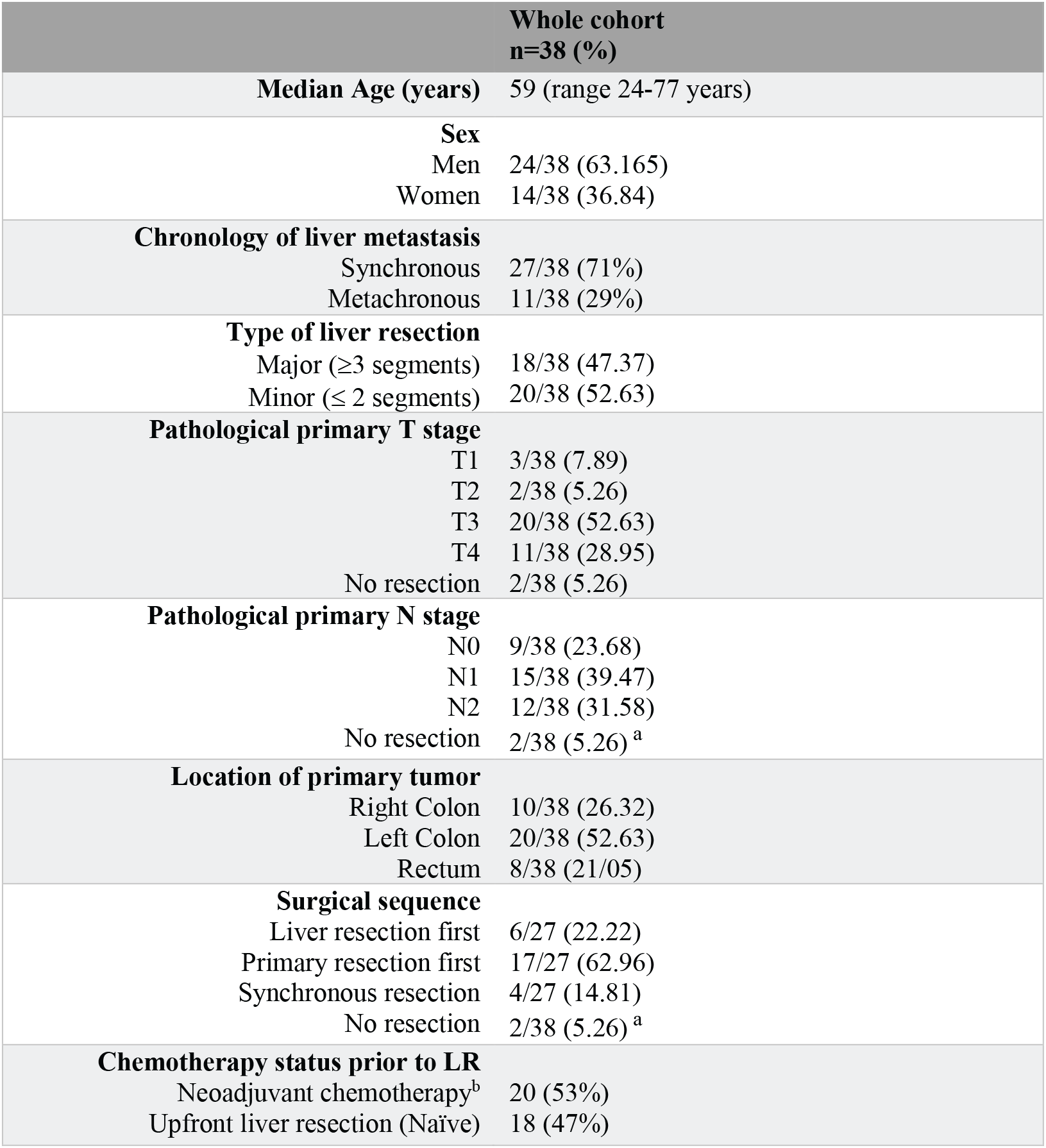
Clinicopathological Details of Patient Cohort. L=Liver resection, NeoCT=Neoadjuvant chemotherapy, ^a^Two Patients no resection of primary, one due to progression of disease and the other due to complete response in primary tumor, ^b^3/7 patients had neoadjuvant FOLFOX interspersed with radiotherapy to primary tumor.

In parallel, tumor samples and matching PDTOs were embedded and processed for morphological and histological analyses (**Figure 1a**). Hematoxylin and eosin (H&E) staining indicated the morphology and tissue architecture of PDTOs was consistent with an adenocarcinoma origin, including solid epithelial structures, glandular-like budding and cysts, with morphological features varying across patients. Positive staining for CK20 and negative staining for CK7 confirmed all PDTOs to be consistent with adenocarcinoma of colorectal origin. Total RNA was then extracted from PDTOs and their parent tumors (*N_Pairs_*= 18) and analyzed using RNAseq. Within matched samples, the RNA abundance log-ratios were mostly centered around zero and as expected abundance data were highly correlated (*rSpearman*: 0.68 to 0.86), demonstrating good consistency between PDTOs and their parent tumors (**Figure 1b, *left and right panels respectively*)**. Genes that were highly expressed in tumor samples (logRPKM > 5) and lowly expressed in PDTOs (logRPKM < 0.05), were mostly genes associated with ECM/stroma (e.g. Collagens, VIM, ZEB1/2, FN1), and infiltrating immune cells (PTPRC, CD8A, CD3D/E/G, TIGIT, and LAG3).

**Figure 1.**
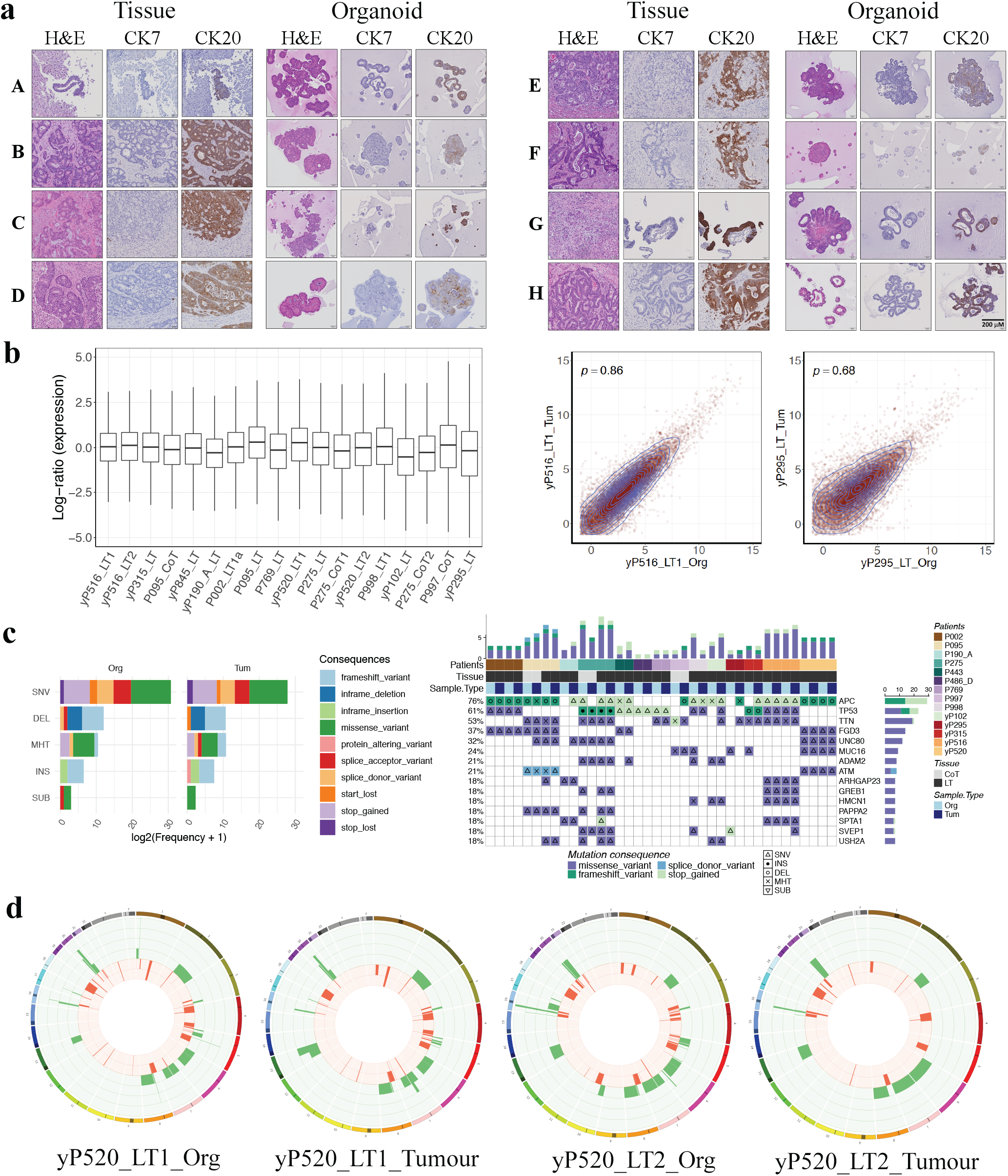
Histological and Molecular Concordance of PDTOs compared with parent tumor. **a)** Histochemical staining of CRLM tumors and matching PDTOs: Morphological concordance with hematoxylin and eosin (H&E) staining; immunostaining for CK7 (negative control) and for CK20, confirming the colorectal cancer origin of PDTOs. **b)** Comparison of total RNA from PDTOs and parent tumors using RNAseq (*NPairs*= 18): Distribution of overall gene expression shown through analysis of log-ratio of gene expression values (*left panel*), and 2 examples illustrating the highest and lowest correlation observed between overall expression in PDTOs and matching tumor sample (Spearman’s coefficient ranging from 0.86 to 0.68, *right panel*). **c)** Comparison of mutational profiles between PDTOs and matching tumors: Histogram showing concordance in number, type and consequence of mutations (*left panel*) and oncoprint showing concordance of highly mutated genes (HMG (*right panel*). **d)** Circos plots illustrating the pattern of copy number variations (gains and losses) in two examples of PDTO and matching tumors, showing the good concordance between PDTO and parent tumors, but highlighting the inter-metastatic heterogeneity between two CRLM from the same patient (yP520_LT1 and LT2, *right and left pairs respectively*).

Similar to previous studies, a high level of genomic consistency was demonstrated between PDTOs and their parent tumor using WGS/WES ^14–16^, in regards to the number, type and consequence of mutations (**Figure 1c,*left panel*)**, the mutation patterns across highly mutated genes (HMG) **(Figure 1c,*right panel)*** and the copy number variation (CNV) patterns **(Figure 1d; Suppl. Figure 2a)**. The most frequent mutation type in both PDTO and tumor samples were single nucleotide variants (SNVs; ~93% of variants in both tumor and PDTOs), while substitutions were the least common DNA alteration in both groups (~0.2% of variants in both PDTOs and tumor samples). Mutations were most frequently responsible for missense variants (~84% of mutation consequences in both tumors and PDTOs), followed by stop-gain (~6% of mutation consequences in tumors and PDTOs) (**Figure 1c**). Between most tumor-PDTO pairs there was high retention of highly mutated genes across the sample pairs, with only one sample (P998) showing lower concordance (**Figure 1c,*right panel***). The top 5 most highly mutated genes within tumor samples included *APC, TP53, TTN, KRAS* and *UNC80* (> 20% of samples), also seen across the matched PDTOs **(Figure 1c).** Finally, while patterns of copy number gains and losses were quite consistent between PDTOs and their matching tumor samples, heterogeneity was detected between different metastases collected from the same patients (**Figure 1d**; **Suppl. Figure 2a)**

Tumor samples collected in this study were all found to be microsatellite stable but displayed features of chromosomal instability (CIN). Of the 73 tumor samples and PDTOs that underwent Whole Exome Sequencing (WES) or Whole Genome Sequencing (WGS) in our study (40 CRLM samples, 9 primary tumor samples, 19 CRLM PDTOs and 5 primary tumor-derived PDTOs), 65 (89%) displayed clear aneuploidy **(Suppl. Table 2)**. The median tumor mutation burden (TMB) across tumor samples was 3.6 mut/Mb (0.021 to 10.91) **(Suppl. Table 2)**. Concordant CNVs detected in matched tumor and organoid pairs reflected a high levels of copy number gain, as well as copy number neutral loss of heterozygosity (cnLOH) **(Suppl. Figure 2b)**. CNV for all samples are available in **Suppl. Table 3**. Finally, in samples that underwent WGS (n = 36), concordant structural rearrangements were detected in tumors and matched PDTOs **(Suppl. Figure 3)**.

Across our tumor and PDTO sample cohort, mutations were identified in genes previously found to be altered in non-hypermutated CRC (non-MSI, CIN positive) (TCGA colorectal cancer, primary tumor analysis n = 224) and microsatellite stable CRC (Yaeger et al, n = 979 metastatic, n = 120 early CRC) ^17,18^ (**Suppl. Figure 4a)**. Specifically, within our CRLM tumors (n = 40), the frequencies of mutations in these genes were: *APC* (68%), *TP53* (60%), *TTN* (35%), *KRAS* (20%), *TCF7L2* (12%), *PIK3CA* (8%), *NRAS* (8%), *FBXW7* (5%), and *SOX9 (3%)* (**Suppl. Figure 4b**), consistent with a study by Pitroda et al that focused exclusively on CRLM (n = 59 tumors)^19^. Mutations involving *MAP2K7* and *PTEN* were rare in our cohort, and mutations in *CTNNB1, SMAD2, FAM123B* or *ARID1A* were absent. This could be due to the low frequency of these mutations in CRC coupled with the moderate size of our sample cohort, or alternatively, they may be less frequently represented in CRLM as compared with primary tumors. Finally, while some of the highly mutated genes in our cohort were common across primary and metastatic samples (*TTN, APC, TP53*, *UNC80…*), some genes were more frequently mutated in primary compared with metastatic samples. For example, mutations in *KRAS, FGD3, MUC12* and *MUC16* were more frequent in CRLM samples, whereas primary samples showed more mutations within *AHNAK2* and *ABCA12*. Collectively, these results consolidate findings from previous studies that PDTOs exhibit similar genomics characteristics to those of matching tumors^14–16^.

### Organoids show heterogeneous responses to in vitro FOLFOX treatment

PDTOs that demonstrated robust growth and stable morphology over consecutive passages were assessed for their sensitivity to FOLFOX. We chose to use this drug combination rather than individual agents to parallel the treatment of patient tumors. Tumor cells were seeded at high density **(Suppl. Figure 5a)** and PDTOs were cultured until macroscopic **(Suppl. Figure 5b)** to better recapitulate the phenotypic heterogeneity and architectural complexity of metastatic tumors, then re-plated as *whole* organoids in fresh Matrigel^®^ into 96-well plates 24 hours prior to treatment with FOLFOX. Determination of IC50 following 72-hour treatment with increasing FOLFOX concentrations highlighted the varying sensitivity amongst the PDTOs (**Suppl. Figure 5c).** PDTOs demonstrating the most different responses were classified as either FOLFOX sensitive (IC50 < 5 μM) or FOLFOX-resistant (IC50 > 50 μM), with a further group of PDTOs representing an intermediate sensitivity group thus classified as ‘semi-sensitive’.

To identify the molecular profile contributing to FOLFOX resistance, 14 PDTOs were subsequently treated for 72 hours with 25 μM FOLFOX (treatment group) or vehicle (control group). Consistent with previous dose-response experiments, resistant PDTOs (n = 3) did not show a reduction in viability under these conditions (as quantified by Cell-Titer Glo 3D), adopting a cystic phenotype with little or no evidence of death (as determined via bright field (BF) and fluorescence microscopy with incorporation of 7-Aminoactinomycin (7-AAD)) (**Figure 2a,***all* **PDTOs images Suppl. Figure 6a-b)**. By contrast, sensitive PDTOs (n = 4) had a reduction in viability of 70-90% compared to vehicle-treated matching controls, supported by the strong incorporation of 7-AAD and loss of organoid architecture. PDTOs classified as semi-sensitive (n = 7) exhibited significantly reduced CTG viability (mean decrease of 20-50% compared to matching controls), with evidence of cell death on imaging, but preservation of overall organoid structure (**Figure 2a; Suppl. Figure 6a-b).** Samples from all 14 PDTOs treated under these conditions were processed for RNA extraction **(Suppl. Table 4)**.

**Figure 2.**
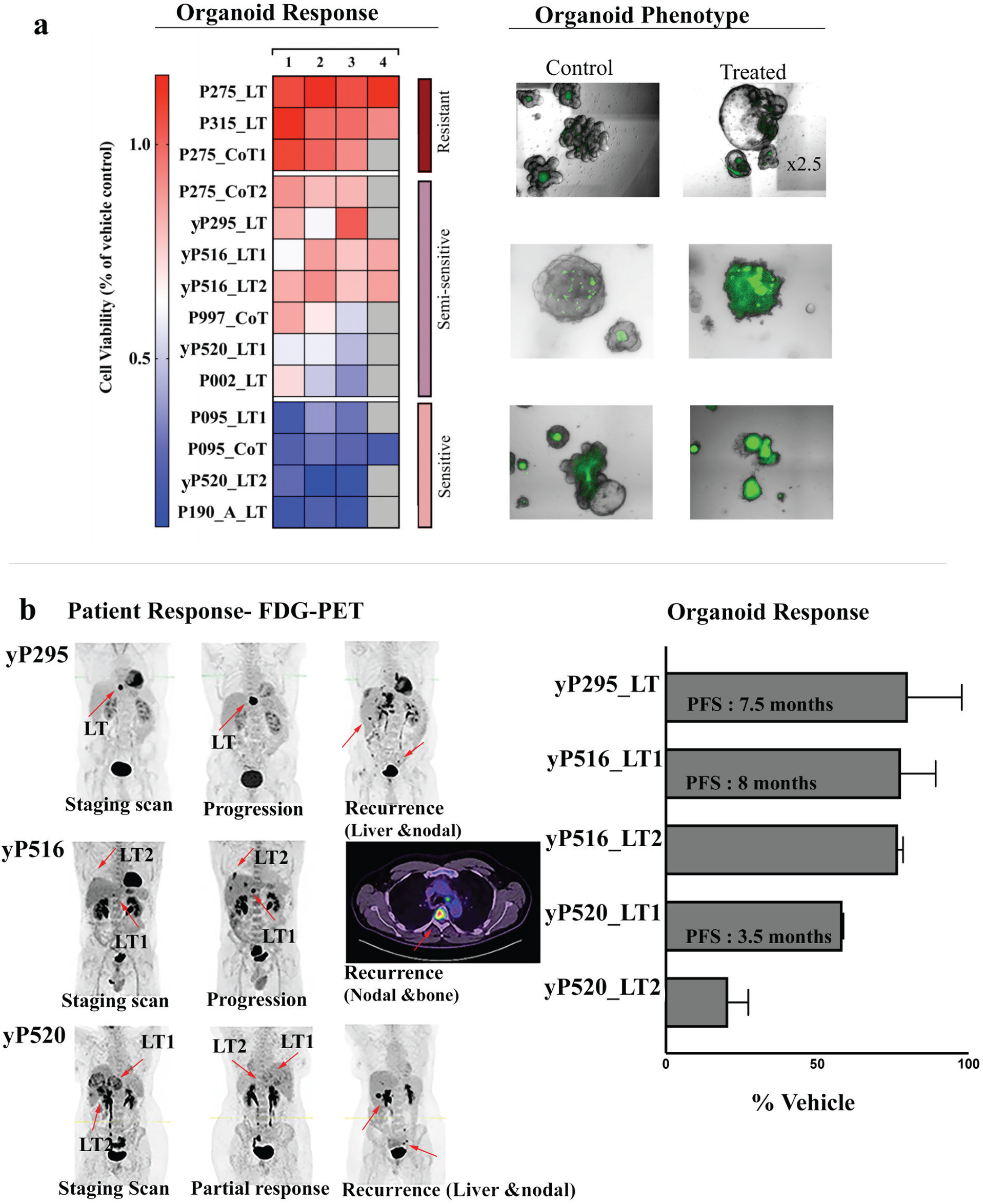
PDTO Response Categories and Concordance with Patient Response. **a)** Heatmap summarizing the PDTOs Response to FOLFOX: 14 PDTOs treated with 25uM FOLFOX for 72h were categorized into resistant, semi-semi sensitive, and sensitive PDTOs based on their viability compared to control assessed using CTG (*left panel*) and imaging (*right panel*). Columns 1-4 represent independent experimental repeats, each one including 3 replicate wells per condition. Grey cells indicate when no 4^th^ experimental repeat was run. **b)** Consistency between organoid and patient response. *Left panel*: representative FDG-PET scanning images taken at staging, after treatment and at time of recurrence from three patients treated with FOLFOX-based neoadjuvant chemotherapy prior to colorectal liver resection. Patient yP295: progression of disease, single liver metastasis. yP516: Mixed partial and progression, LT1 and LT2 collected showed progressive disease. yP520: partial response; LT1 showed progressive disease on repeat imaging, LT2 continued to respond to treatment. *Right panel*: Histogram summarizing the response to in vitro FOLFOX treatment of PDTOs derived from tumors described above (collection from metastases indicated by red arrows in left panel), demonstrating correlation with patient response and ability to detect heterogenous responses between different liver metastases in the same patient (yP520). All three patients had very short progression free survival, as detailed on the histogram.

To ensure that the PDTOs response to FOLFOX reflected the clinical response of matching tumors, we compared the PDTO responses with the treatment responses on FDG-PET (18F-fluorodeoxyglucose-Positron Emission Topography) in patients who had received FOLFOX-based neoadjuvant chemotherapy prior to resection of their CRLM (n = 5 tumors from 3 patients) (**Figure 2b,*left panel***). Four of the five tumors demonstrated progression in the patient on imaging, and matching PDTOs all had a viability of more than 60% in response to FOLFOX *ex-vivo*, with maintenance of organoid architecture. In contrast, one of the tumors showed a partial metabolic response in the patient and the corresponding PDTO was sensitive with loss of architecture (**Figure 2b, *right panel***). Interestingly, one patient had a differential response between their two metastases (progressive vs partial response), and this difference was reflected in the PDTO responses *in vitro* (semi-sensitive vs sensitive). Our results thus demonstrate that PDTOs recapitulate the metabolic responses on FDG-PET to FOLFOX, including the ability to detect heterogeneous inter-metastatic responses, suggesting PDTOs represent a robust model to analyze molecular mechanisms that contribute to FOLFOX resistance in poor responders.

### FOLFOX-resistant PDTOs are enriched for E2F and late cell cycle regulation pathways

RNAseq was then used to interrogate the transcriptomic profiles of PDTOs within the treatment and control groups. Multiple analyses were performed to compare FOLFOX-treated and control (vehicle-treated) groups in all three response categories (FOLFOX resistant, -sensitive and -semi-sensitive, n = 14) (**Suppl. Figure 7a, and Suppl. Table 5**). The transcriptomic profile of semi-sensitive PDTOs exhibited a larger overlap with that of the FOLFOX-sensitive PDTOs than with the profile of resistant organoids. Consequently, fewer differentially expressed genes (DEGs) were identified between the semi-sensitive and sensitive organoid subgroups (**Suppl. Figure 7a**, Ctrl_Sen_SemiSen, N = 97 DEGs between sensitive and semi-sensitive PDTOs, compared with Ctrl_Res_SemiSen, N = 496 DEGs between resistant and semi-sensitive PDTOs and Ctrl_Res_sen, N = 623 DEGs between resistant and sensitive PDTOs). We therefore focused our analysis on comparing the DEGs between FOLFOX-treated and control groups in the resistant (n = 3) and sensitive (n = 4) PDTOs (*N_DEGs_* = 2121 with absolute log fold change > 1 and adjusted p-value < 0.05, now referred to as ***Treatment_Res_Sens*** signature) (**Figure 3a, Suppl. Table 5a)**. KEGG pathway analysis using the ***Treatment_Res_Sens*** signature identified significant pathways associated with DNA replication and repair, cell cycle and TP53 signaling. These results were consistent with the top significant biological processes (BPs) identified using the gene ontology (GO) enrichment analysis (**Figure 3b, Suppl. Table 6; Sheets 1 & 2).**

**Figure 3.**
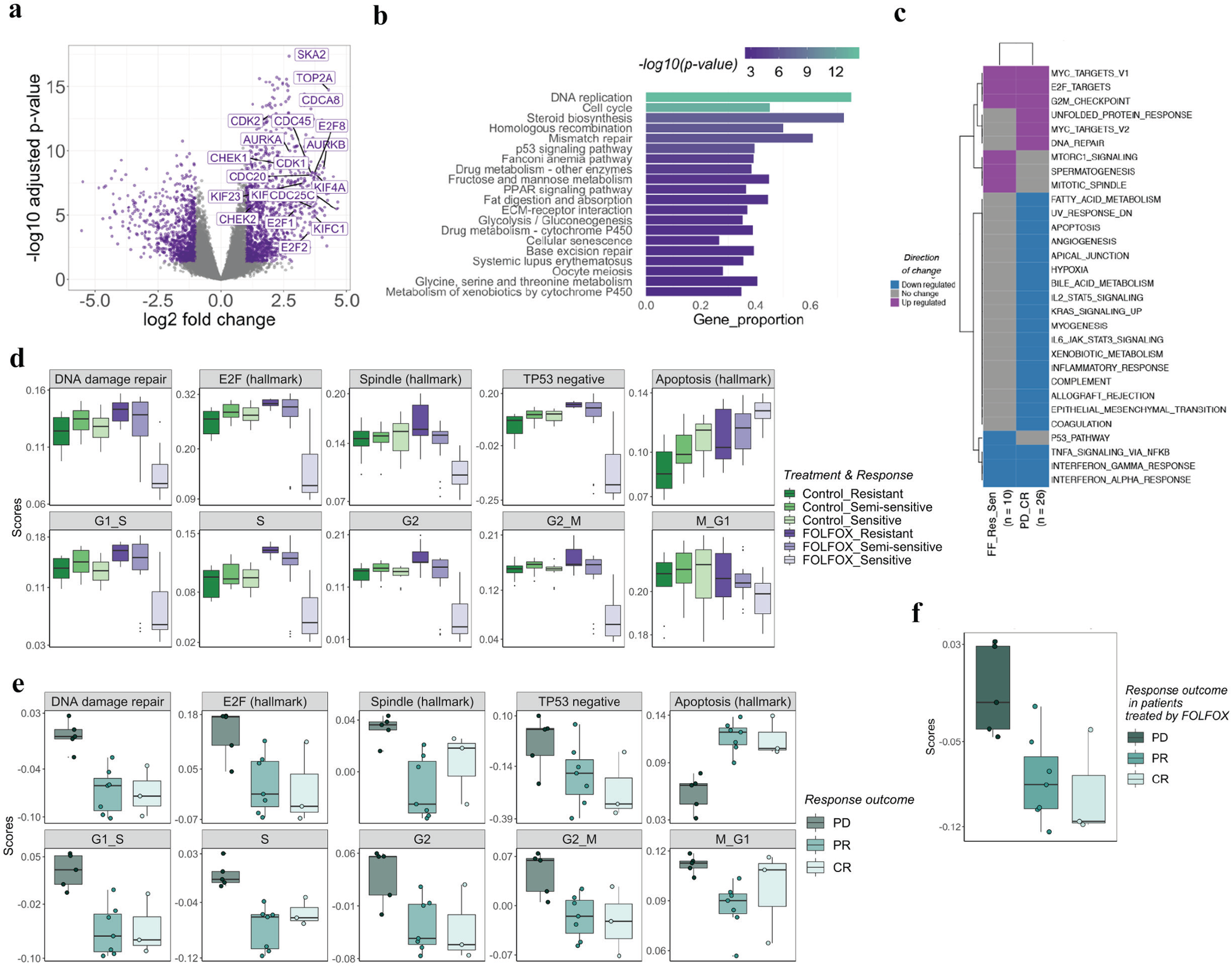
RNAseq Analysis of FOLFOX-Treated PDTOs. **a)** *Treatment_Res_Sens* signature: DEGs between FOLFOX-induced genes in resistant (n = 3) vs sensitive (n = 4) PDTOs (3 biological repeats, each with 3 technical replicates, for all vehicle and FOLFOX-treated PDTO samples) (*NDEGs* = 2121 with absolute log fold change > 1 and adjusted p-value < 0.5). **b)** Top 20 significant enriched pathways based on KEGG pathway analysis: Using the DEGs identified in (a), we found significant pathways associated with DNA replication and repair, cell cycle and TP53 signaling. **c)** Significant Hallmark gene sets found to be up-or down-regulated in FOLFOX resistant PDTOs (FF_Res_Sen = resistant vs sensitive samples in FOLFOX treated PDTOs, *left column*) and in tumor samples from patients whose disease progressed on FOLFOX (PD_CR = samples from patients with progressive disease vs complete response to FOLFOX, *right column*). **d)** Singscore plots for PDTOs in vehicle-and FOLFOX-treated groups stratified by treatment outcome, analyzed against several Cell cycle, DNA repair, E2F, spindle, TP53 and apoptosis MSigDB hallmark signatures. **e)** Scores obtained using the same hallmark signatures as in (**d**) for tumor samples from patients treated with neoadjuvant FOLFOX and stratified by treatment outcome. **f)** FF_Res_Sens signature scores in tumor samples: DEGs between resistant and sensitive samples within the *FOLFOX-treated* groups were identified ( n = 1354 with absolute log fold change > 1 and adjusted p-value < 0.05) and then used to score to FOLFOX-treated tumor samples, highlighting the higher score in tumors that progressed under FOLFOX treatment.

To validate these results, we also performed competitive gene-set testing of MSigDB hallmark signatures^**20**^ against the whole transcriptome of FOLFOX-resistant vs FOLFOX-sensitive PDTOs, as well as against all other comparisons tested above between FOLFOX-resistant, -semi-sensitive and -sensitive groups **(Suppl. Figure 7b)**. FOLFOX-resistant PDTOs demonstrated an up regulation of E2F targets, G2M checkpoints and mitotic spindle signatures, and a down-regulation of the TP53 pathway genes compared to sensitive PDTOs (**Figure 3c, left column; Suppl. Figure 7c)**.

To further support these findings, we used *singscore^21^*, an unsupervised, single-sample gene-set scoring method, to score individual PDTOs against several Hallmark molecular signatures related to the cell cycle, E2F pathway, DNA damage response and apoptotic pathways **(Figure 3d)**. Scores for most of these signatures were largely similar across all categories in vehicle-treated PDTOs. However, after exposure to FOLFOX, the transcriptome of resistant PDTOs came strongly enriched in scores for E2F targets, DNA damage repair, TP53 pathway downregulation, as well as late cell cycle-signatures, when compared to FOLFOX-sensitive PDTOs. In particular, resistant PDTOs exhibited high scores for G_1_/S, S-phase, G_2_, G_2_/M transition and mitotic spindle hallmarks, whereas the hallmark signature for mitosis to G_1_ transition was not significantly different between FOLFOX response categories. In contrast, scores for the apoptosis hallmark signature were increased in FOLFOX-sensitive PDTOs compared to resistant samples **(Figure 3d)**. Overall, scores for the semi-sensitive samples were spread between those of resistant and sensitive PDTOSs. Taken together these data demonstrate that resistant PDTOs exhibit enriched transcriptomic features for E2F signaling and for late cell-cycle genes when exposed to FOLFOX, with a concomitant downregulation of p53 signaling.

### Liver metastases that progress under neoadjuvant FOLFOX display similar pathway enrichment to FOLFOX-resistant PDTOs

To assess whether the transcriptional profiles of FOLFOX-resistant PDTOs identified above reflect the resistance process within patients, tumor samples derived from patients treated with neoadjuvant FOLFOX were independently analyzed. Differential gene expression analyses were performed to compare tumor subgroups selected according to their clinical imaging response within patient while under chemotherapy (**Suppl. Table 5**). The largest number of DEGs (*N_DEGs_* = 2578) was detected when comparing tumors that exhibited progressive disease (PD) and those demonstrating a complete response (CR) under FOLFOX treatment (Called PD_CR; **Suppl. Figure 8a; Suppl. Table 5)**. Gene-set testing using MSigDB Hallmark signatures indicated that tumors that progressed under FOLFOX treatment (PD) exhibited a similar up-regulation of E2F target, *MYC* target and G_2_M checkpoint signatures as observed for PDTOs. Consistent with the results of the KEGG and GO BPs analyses using the PD_CR DEGs (**Suppl. Table 6; Sheets 3 & 4**), several Hallmark signatures associated with inflammatory response, leukocyte migration and cell adhesions were strongly downregulated in samples with PD. Most of these signatures were not detected in PDTOs that lack stromal and immune components, however a significant downregulation of alpha/gamma interferon (IFNα;/IFNγ;) response and tumor necrosis factor alpha (TNF-α signaling (via NF-κB)) was identified in both the PDTO and tumor data (**Figure 3c right column and Suppl. Figure 8b**).

In support of both of these findings, DEGs that discriminated PD from CR tumors included a large number of genes from the E2F Hallmark signatures (42/200, all upregulated in tumor samples exhibiting PD) as well as half of the genes represented in immune and stromal hallmark signatures (136/282, all downregulated) **(Suppl. Figure 8c)**. These observations were further reinforced by results of the KEGG and GO enrichment analyses with DEGs **(Suppl. Table 6)**. *Singscore* signature analysis uncovered that tumors that progressed under neoadjuvant FOLFOX treatment (PD) had a significantly lower scores against stromal and immune related signatures than those with partial or complete metabolic response, including signatures reflecting interferon, natural killer (NK) and T cell activation (**Suppl. Figure 8d**). Together these results imply that there is a lower level of immune activation in tumors that progressed under treatment compared with those that partially or completely responded.

Signature analysis was again used to compare the transcriptome of tumors demonstrating PD, CR and partial response (PR) against the Hallmark molecular signatures enriched in FOLFOX-resistant PDTOs (DNA damage repair, E2F target, cell cycle phases, spindle assembly, TP53 negative, and apoptosis signatures). Results indicated that, similar to resistant PDTOs, tumor samples collected from patients who progressed under neoadjuvant FOLFOX (PD) scored significantly higher for most of these molecular signatures when compared with tumors exhibiting a PR or CR, apart from the Mitosis to G1 transition (not significantly different) and the apoptosis hallmark signatures (higher in PR and CR) (**Figure 3e)**.

In the clinical setting, if patients are potentially curable, sampling of CRLM prior to treatment risks the tumor seeding and therefore molecular characterization is generally only possible following neoadjuvant chemotherapy and surgical resection. Therefore, DEGs between resistant and sensitive organoids were only identified within the *FOLFOX-treated* group (signature hereafter called ***FF_Res_Sens***, **Suppl. Fig 7a** and **Suppl. Table 5**). This signature, comprising 1354 DEGs, shared 610 genes with the ***Treatment_Res_Sen*** signature used above (which includes genes where the transcriptional change in response to FOLFOX is in the opposite direction in resistant vs sensitive PDTOs, when compared to their respective DMSO-treated controls). Similarly, enrichment in pathways and genes associated with cell cycle, mitotic spindles and TP53 signaling was evident in this analysis. We then scored individual tumors from the PD, PR and CR response groups using the ***FF_Res_Sens*** signature (from organoids) and established that tumors derived from patient with PD scored significantly higher than those with partial or complete responses (**Figure 3f**).

Collectively these results indicate that resistant PDTOs which have been exposed to FOLFOX *in vitro,* and metastatic tumors that progressed under neoadjuvant FOLFOX treatment in patients, share transcriptomic characteristics that reflect the enrichment of E2F pathway, DNA damage repair, S to G_2_/M phases and spindle assembly checkpoint genes, whilst displaying downregulated TP53 pathway, apoptosis, IFN response and TNF signaling signatures. Thus, our data suggests that FOLFOX-resistant tumor samples and PDTOs are cell cycle-arrested in S phase or during the G_2_/M transition.

### Differential genomic alterations are not sufficient to explain FOLFOX resistance in metastatic colorectal cancer PDTOs and tumors

To determine whether genomic alterations could underlie the transcriptomic differences between FOLFOX-resistant and -sensitive PDTOs and between FOLFOX-treated tumors with different clinical outcomes, we first analyzed the mutational profile of our PDTO cohort (**Suppl. Table 7)**. A small number of genes showed differential mutation profiles between FOLFOX-resistant and -sensitive PDTOs. All FOLFOX-resistant and 50% (3/6) semi-sensitive PDTOs harbored somatic *TP53* variants, whereas 80% (4/5) sensitive PDTOs were wild-type for *TP53* **(Figure 4a).** Interestingly, the consequence of these variants in the resistant PDTO was frameshift mutations, caused by INDELs (insertion c.361dupT in exon 4 or deletion c.635_636delTT in exon 6), which are predicted to be highly deleterious driver mutations, causing loss of function (TIER 1 prediction according to Cancer Genome Interpreter, https://www.cancergenomeinterpreter.org). None of our PDTOs exhibited *RB1* gene mutations, which encode the retinoblastoma tumor suppressor, a known regulator of the E2F family members. Rather, our RNAseq data highlighted that PDTOs carrying these frameshift *TP53* mutations exhibited a high mRNA expression for multiple genes that are usually down-regulated by *TP53* via the *p53-p21-DREAM-CDE/CHR* complex^22–24^, including *PLK1, CDC20, CDC25A, CDC25C, MCM5, BIRC5, CCNA2, CCNB1, CCNB2, CKS1B* and kinesins (*KIF2C, KIF23, and KIF24*). Of note, *TP53* variants in the semi-sensitive PDTO group resulted in missense mutations in exon 6, while the only *TP53*-mutated sensitive sample carried a loss of function mutation (c.641A>G, according to the IARC TP53 Database https://p53.iarc.fr/).

**Figure 4.**
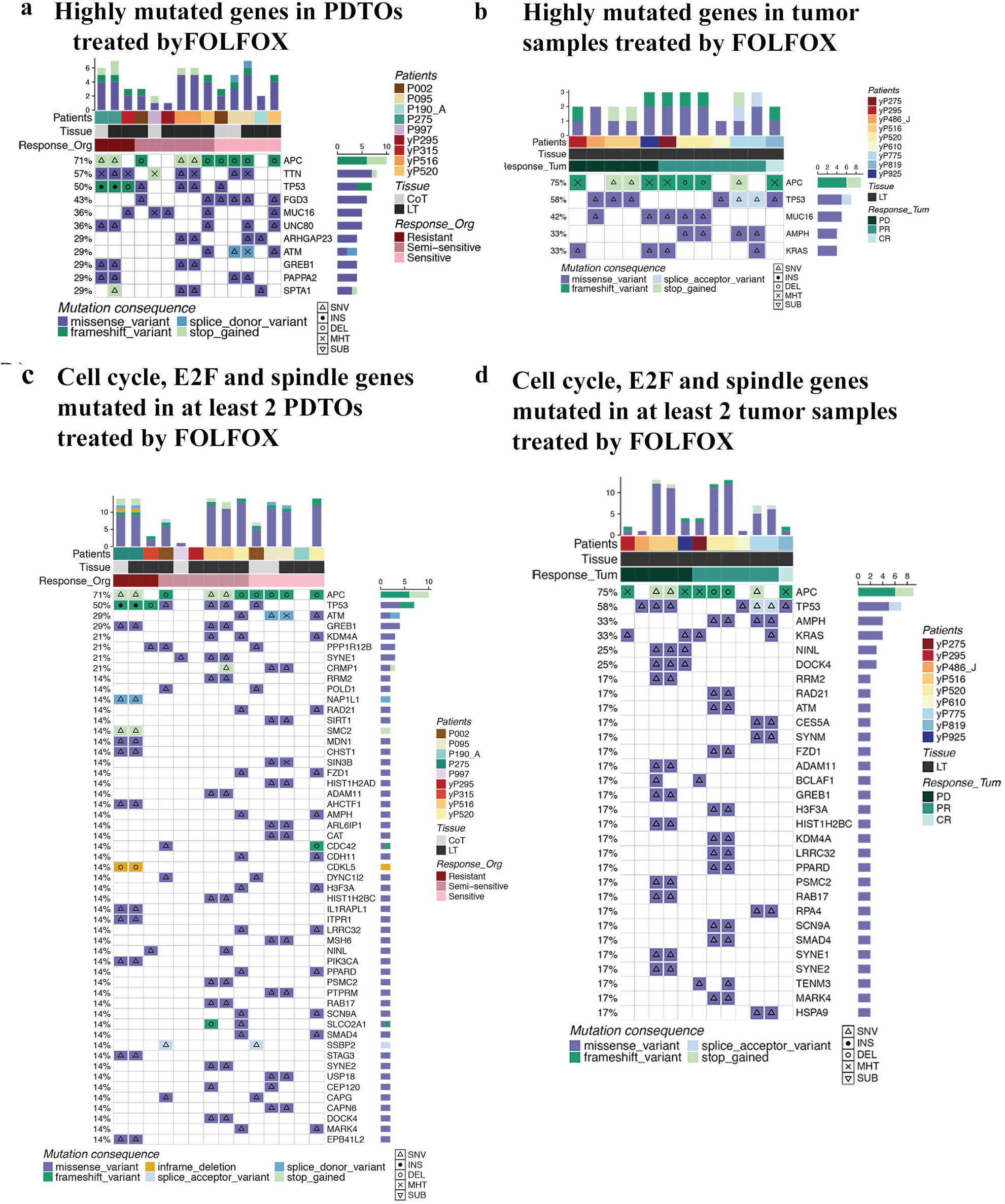
Mutational Profile of FOLFOX Treated PDTOs and Tumors. **a)** Oncoprint showing highly mutated genes in PDTOs grouped according to their response to *in vitro* FOLFOX, indicating the mutation type (Single nucleotide variant – SNV, Insertion – INS, Deletion – DEL, Multi-Hit – MHT, or Substitution – SUB), mutation consequence. **b)** Oncoprint showing highly mutated genes in FOLFOX-treated tumor samples stratified according to patient tumor response (partial response (PR), progressive disease (PD) or complete response (CR)), indicating that TP53 variants are not enriched in tumors that progressed under FOLFOX treatment. **c)** Oncoprint depicting mutations in cell cycle, E2F and mitotic spindle-related genes found to be mutated in at least 2 PDTO samples. **d)** Oncoprint depicting cell cycle, E2F and mitotic spindle-related genes mutated in at least 2 tumor samples.

In contrast, 60% (3/5) sensitive PDTO displayed loss of function variants within the DNA damage repair gene *ATM* **(Figure 4a)**. *ATM* loss of function mutations have been reported as common in CRC^25^. *ATM* appears to be required for tumor cells to repair oxaliplatin-induced double-stranded DNA damage^26^, and IHC-based detection of *ATM* loss is associated with better overall survival following oxaliplatin-based treatment in CRC patients^27^. Overall, TP53 and ATM mutations were mutually exclusive within our PDTO cohort, consistent with the TCGA data^**17**^. Interestingly, 80% (4/5) FOLFOX-sensitive PDTOs were found to carry a missense variant in the *FGD3* gene, which encodes an inhibitor of cell motility. High *FGD3* expression has been found in oxaliplatin and CapeOX (capecitabine/oxaliplatin) resistant gastric cancer cells ^28^ and was recently shown to correlate with favorable prognosis in breast and other cancers^29^. In the present cohort *FGD3* was also mutated in two of the semi-sensitive PDTOs, and *FGD3* and *TP53* mutations were concomitantly detected in 2 PDTOs (one FOLFOX-sensitive, one semi-sensitive).

We then compared the resistant and sensitive PDTOs with the tumor samples from PD vs CR patients to find common genomic alterations that may underlie FOLFOX resistance **(Figure 4b)**. We did not detect any specific enrichment in frameshift *TP53* variants in tumors that progressed under FOLFOX compared to tumors exhibiting partial or complete response, suggesting that the mutational profile of *TP53* identified in FOLFOX-resistant PDTOs is insufficient to explain the resistance to this drug regimen.

Next, we determined the mutational profile of all the genes associated with cell cycle, E2F, and spindle assembly from MSigDB signatures (N = 2626 genes). From these genes, 131 were mutated in at least one of the 14 PDTOs treated with FOLFOX *in vitro*, and only 53 genes were mutated in at least 2 PDTOs **(Figure 4c)**. In tumor samples collected after neoadjuvant FOLFOX treatment, there were 95 genes from the pathways mutated in at least one sample, and only 30 genes mutated in at least 2 samples (**Figure 4d**).

Importantly, while a number of these genes overlapped between PDTOs and tumor samples, we did not identify mutations that were common between resistant PDTOs and tumor samples exhibiting PD under FOLFOX and which might have explained E2F and late cell cycle pathway activations. We further examined genes associated with DNA damage repair. Among the 91 genes listed in this pathway^30^, only 5 were mutated in our PDTOs and 5 in tumor samples treated with neoadjuvant FOLFOX *ATM* was the only common gene mutated in both types of samples, including 60% (3/5) FOLFOX-sensitive organoids **(Suppl. Figure 9a)**, and 33% (2/6) tumor samples (**Suppl. Figure 9b)**that demonstrated a partial response to FOLFOX in patients. Thus, mutation in DNA damage repair genes were not sufficient to discriminate good from poor FOLFOX responders in PDTOs and tumors.

Collectively these results identify differential mutations patterns between FOLFOX-resistant (*TP53* frameshift) and sensitive (*ATM/FGD3*) PDTOs, but also establishes that these mutations alone do not discriminate tumor samples from patients with PD or with CR after neo-adjuvant FOLFOX treatment. This suggests that genomic alterations alone are unlikely to play a major driving role in the resistance of colorectal cancer liver metastases to FOLFOX, and that they cannot explain the differential activation of the E2F and p53-independent cell cycle checkpoint pathways detected in FOLFOX-resistant PDTOs and tumor samples.

### FOLFOX-resistant PDTOs undergo treatment-induced cell cycle arrest in S phase

To gain additional insight into the mechanism that drives FOLFOX resistance in metastatic CRC PDTOs, we examined their apoptotic cell death, DNA damage and proliferative behaviors during and after exposure to this cytotoxic combination. As shown earlier, resistant PDTOs developed large cystic phenotypes when exposed to FOLFOX **(Figure 5a –*upper panel***). Dead tumor cells were observed only within the lumen of cystic organoids, with viable epithelial tumor cells forming the luminal wall. This phenotype remained stable for several days after FOLFOX withdrawal. In contrast, sensitive PDTOs showed cell death with loss of organoid architecture **(Figure 5a- *lower panel*)**. The loss of cell viability was confirmed using cleaved caspase 3 (cCas3) staining (**Figure 5b**), confirming the strong activation of apoptotic pathways detected in the transcriptomic profile of these sensitive PDTOs. While the majority of cells within resistant PDTOs remained alive, apoptotic cells were also detected in their lumen after FOLFOX exposure (**Figure 5b**).

**Figure 5.**
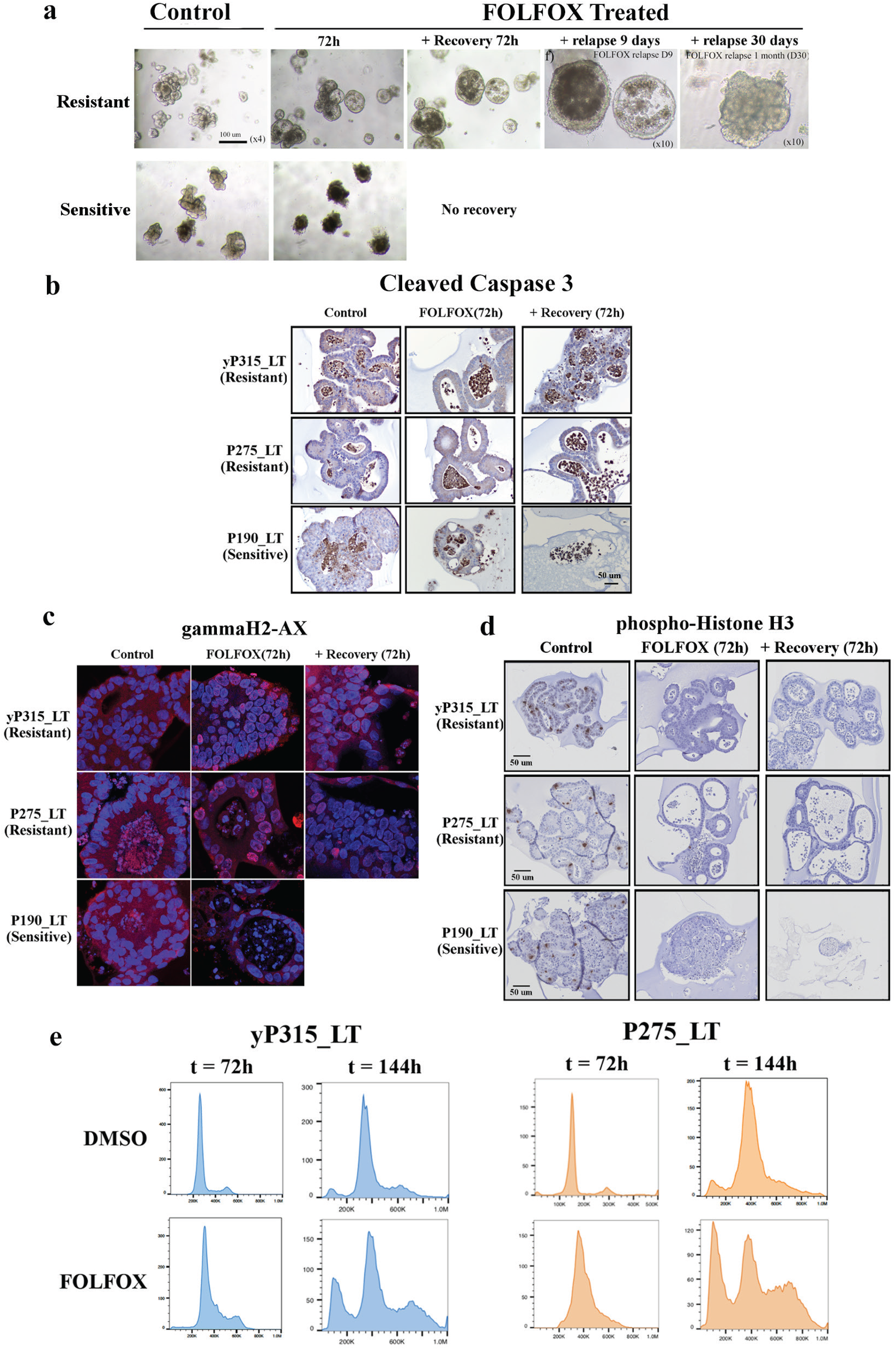
FOLFOX-Resistant PDTOs demonstrate a reduction in proliferation and cell cycle arrest. **a)** Representative bright-field images of resistant and sensitive PDTOs prior to treatment, after 72 hours of treatment and at various time points after FOLFOX withdrawal, illustrating the characteristic cystic phenotype observed in all resistant PDTOs during and shortly after exposure to FOLFOX (*upper panel, first 3 images)*. 9 days after treatment withdrawal PDTOs still display a large lumen but have increased in size *(upper panel, 4^th^ image)*. After 1 month of relapse organoid have completely reverted to a complex branching phenotype *(upper panel, final image)*. In contrast, sensitive PDTOs show darkening of the organoid and loss of architecture after 72 hours of treatment and are most often not sustained in culture beyond that point (*lower panel*). **b)** Immunohistochemical staining for cleaved caspase 3 (cCas3), highlighting the large proportion of apoptotic cells in sensitive PDTOs after exposure to FOLFOX (P190_LT, *bottom panel*); apoptotic cells are also detected in the lumen of resistant organoids (P315 LT, P275_LT, *top and middle panels*), but their main epithelial layers don’t show signs of apoptosis. **c)** Representative γH2Ax immunostaining and DAPI nuclear staining in three independent PDTO lines, illustrating the large amount of DNA damage detected after a 72-hour FOLFOX treatment in resistant (*top and middle panels*) and sensitive organoids (*bottom panel*); In resistant PDTOs, most cells display minimal to no residual γH2Ax foci 72 hours after FOLFOX withdrawal, while a small minority of cells have continued accumulating damage (*far right panel*). In contrast, treatment sensitive PDTO are unable to repair their DNA and are mostly dead after a 72-hour recovery. **d)** Representative phospho-Histone H3 (pHisH3) immunostaining images of FOLFOX-resistant and sensitive PDTOs, illustrating the proliferation arrest in response to FOLFOX treatment. Note that proliferation has not resumed by 72 hours after treatment cessation. **e)** Flow cytometry analysis of cell-cycle in resistant PDTOs following treatment with vehicle or FOLFOX for 72h (t = 72h) or treatment for 72h followed by a 72-hour recovery period in the absence of drug (t = 144h). Magnification bars (b, d) = 50um; N = 3 independent repeats.

Since our transcriptomic analysis suggested that the DNA damage repair pathway was activated in FOLFOX-resistant PDTOs, we quantified double-stranded DNA breaks in these organoids during and after exposure to FOLFOX, in comparison with FOLFOX-sensitive PDTOs. Using γH2Ax staining, the presence of DNA damage foci was robustly detected in resistant as well as sensitive PDTOs after a 72-hour FOLFOX treatment **(Figure 5c)**. 72 hours after FOLFOX withdrawal, resistant PDTOs exhibited lower levels of γH2Ax foci, while cells within sensitive PDTO were no longer viable **(Figure 5c)**. These results indicate that FOLFOX-resistant PDTOs appear to undergo a similar extent of FOLFOX-induced DNA damage to that detected in sensitive organoids, but that they are capable of repairing this damage rather than activating apoptotic pathways.

Furthermore, despite the enrichment of E2F pathway transcriptomic signatures detected above in FOLFOX-resistant PDTOs, no evidence of growth was detected in resistant PDTOs during treatment and up to 72 hours after drug withdrawal, as suggested by bright field imaging **(Figure 5a)**. In accordance with this observation, phospho-Histone H3 (pHisH3) staining confirmed that resistant PDTOs were not proliferating by the end of the 72-hour treatment period, as well as 72 hours after FOLFOX withdrawal, suggesting they may be undergoing cell cycle arrest when exposed to FOLFOX (**Figure 5d**). However, several days after treatment withdrawal, these samples recovered well and grew rapidly **(Figure 5a, final image)**. Since these PDTOs exhibit inactivating *TP53* mutations as well as clear signs of transcriptional activation of G_1_/S, G_2_/M and SAC genes, we hypothesized that treatment with FOLFOX may induce a *TP53* independent cell cycle arrest either during the G_1_/S or G_2_/M transition or in early mitosis, prior to interphase.

To determine the impact of FOLFOX treatment on cell cycle progression, a flow cytometry analysis using propidium iodide was performed on FOLFOX-resistant PDTOs following a 72-hour treatment with FOLFOX, with or without a further 72-hour recovery period. We found that, after a 72-hour incubation with FOLFOX, resistant organoids exhibited an increased percentage of cells in early S-phase compared to DMSO-treated controls (**Figure 5e**). In contrast, 72 hours after withdrawal of FOLFOX, surviving cells were predominantly accumulating in G_2_/M (**Figure 5e**), indicative of cell cycle arrest since they did not show any evidence of active cell division at that time **(Figure 5d).** Apoptotic cells were also detected at this time point, confirming our activated caspase 3 staining results. Together these results demonstrate that cells from resistant PDTOs cease dividing and remain in early S-phase during exposure to FOLFOX, that they progress through pre-mitotic (S and G_2_) phases and accumulate in G_2_/M as they start repairing their DNA immediately after FOLFOX withdrawal, and that they later resume proliferation and recover their original complex branching phenotype.

### TP53-independent DNA-damage response kinases identified as potential novel treatments in mCRC

To identify potential novel compounds that could be used to target FOLFOX-resistant PDTOs, either as a monotherapy or in combination with FOLFOX, we performed an automated drug screen assay using a bank of 429 targeted kinase inhibitors on 3 independent PDTO lines generated from CRLM that had progressed during neoadjuvant FOLFOX-based treatment **(Suppl. Table 8-*Sheet 1*)**. PDTOs were seeded and grown for 24 hours before being exposed to 2 doses (0.5um and 5uM) of each compound. 72 hours after treatment onset, viability was first assessed using the Cell-Titer Glo 3D (CTG), then confirmed using bright field (BF) imaging at 0, 24, 48 and 72 hours after treatment as well as imaging with the viability marker 7-AAD at 72 hours.

For identification of hits, strict cut-offs were applied to the CTG data: Only compounds that had a percentage fold change (PercFC) less than or equal to 50% *and* a significant Z-score less than or equal to −3 were considered as hits (see methods section). Further to this, we incorporated phenotypic analysis as a secondary measure for hit selection by calculating the multi-dimensional perturbation value (mp-value) and the Mahalanobis distance^31^. These calculations allowed identification of hits by measuring the magnitude of morphological differences between a test compound and DMSO (see Methods). In addition, all BF and 7-AAD images were individually checked to detect potential seeding issues and confirm viability/death in comparison with CTG values. Outcomes detected by imaging were stratified into four categories based on manual review of the imaging: alive with growth (AliveG), whereby the size of individual organoids has increased between 0 and 72 hours; alive but cytostatic (AliveC), with no size increase since seeding; death with growth (DeathG), where organoid size has increased before their death, and death with reduced size (DeathC) **(Suppl. Figure 10)**.

Details of PDTO responses against all compounds detected using CTG and imaging results are provided in **Suppl. Table 8.** Of the compounds tested at 0.5 μM, 24/429 (5.6%) were considered as hits in at least one PDTO **(Suppl. Table 8 - *Sheet 2***), with only 3 being effective in all three PDTOs (**Figure 6a, Suppl. Figure 11a**). At 5 μM, 192/429 (45%) compounds were considered as hits in at least one PDTO, 133 (31%) in at least 2 PDTOs and 71 (16%) in all three (**Figure 6a, Suppl. Table 8 –*Sheets 2 and 3*)**. Further analysis highlighted that of the compounds that were efficient on all 3 PDTOs, 38% (27/71) directly targeted kinases involved in cell cycle progression and DNA damage response pathways, such as CDKs, PLK, Aurora kinases, ATM/ATR **(Suppl. Table 8 –*Sheet 3-5*)** and CHK (**Suppl. Figure 12a-c).** Upon closer examination of compound families, most of the drugs known to preferentially target Aurora Kinases A/B, PLK, CDK1-2 and CHK1-2 were among the hits. In addition, other compounds which effectively targeted all three PDTOs (when used at 5μM) included some inhibitors of the PI3-kinase/mTOR/AKT growth pathway, the SRC/FAK pathway, or the MAPK/ERK pathway.

**Figure 6.**
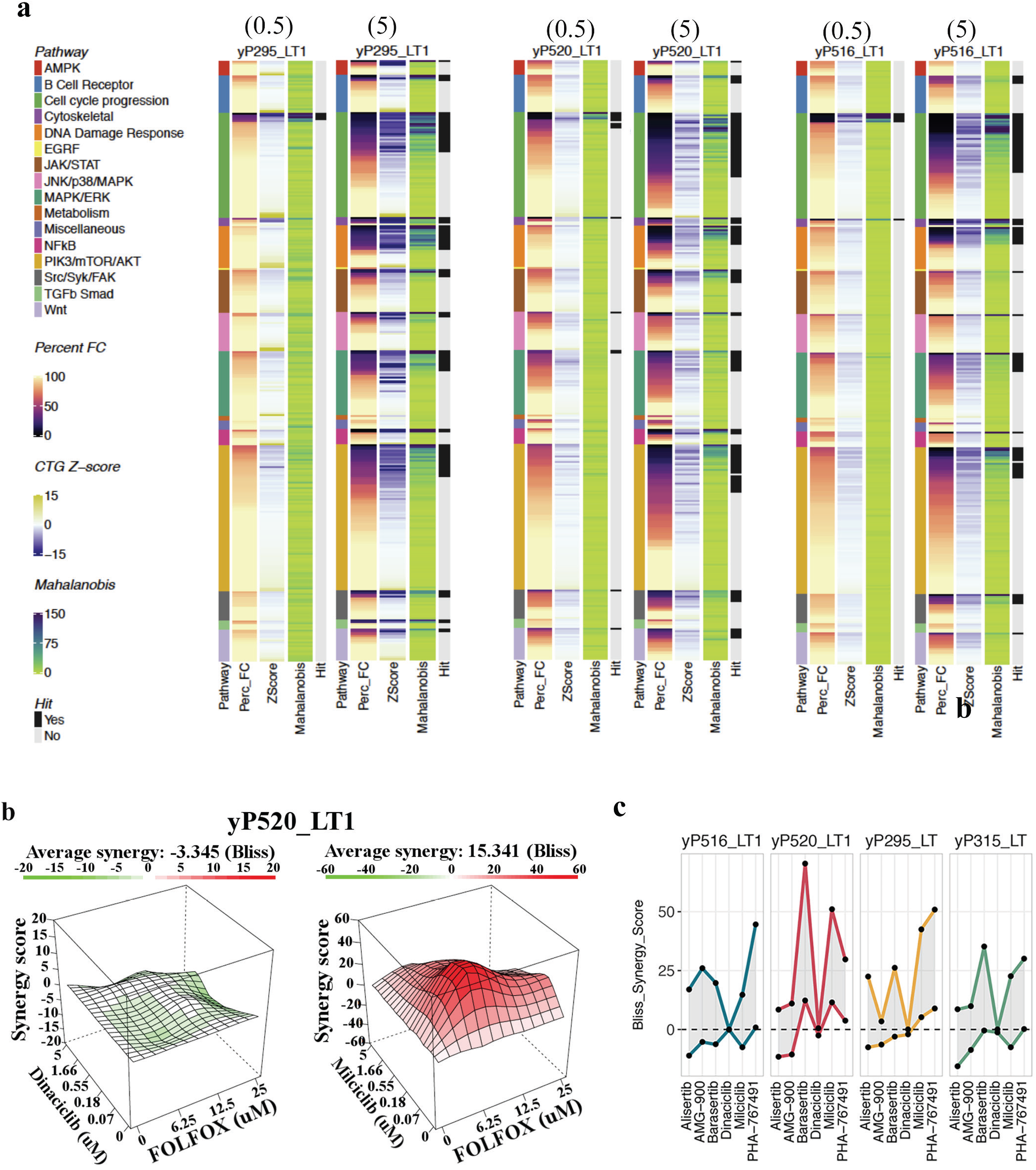
Summary of Primary and Secondary Organoid Screen. **a)** Heatmap summarizing results from the primary kinase inhibitor screening assay performed on PDTOs generated from 3 independent patient tumors that progressed under neoadjuvant FOLFOX treatment (yP295_LT1, yP520_LT1, yP516_LT1). Heatmap columns represent percentage survival (Percent FC, quantified using CTG), the CTG Z-score values, and the Mahalanobis distance values (calculated from the imaging data). Additional bars indicate which pathway compounds used are stratified into and which compounds have been classified as positive hits from the PDTOs Screen. For each PDTO, results are provided for treatment at both 0.5uM and 5um. For better visualization, compounds are stratified based on the main pathway they are ranked into and on their *Percent FC* value for each individual patient; therefore, compounds across patients are not always directly comparable (See Suppl Figure 11 and Suppl tables for direct compound comparison across patients). **b)** Example 3D plots illustrating the possible synergy between FOLFOX and selected compounds in the yP520_LT1 PDTO, calculated using the BLISS model from the CTG data for one ineffective (Dinaciclib) and one effective (Milciclib) compound; Scores above 1 are indicative of the occurrence of synergy, while scores around 0 reflect probable additivity and negative scores suggest antagonism. **c)** Graph summarizing synergy scores calculated using the Bliss Reference Model across 4 PDTO lines. Starting from the left of each graph, the lower line represents the average synergy score across the dose-response grid while the upper line indicates the maximum score that can be reached for any given dose-combination across the grid, highlighting the presence of concentration windows for which strong synergy is detected between FOLFOX and inhibitors of cell cycle progression.

To validate these results and determine if there was potential synergy between FOLFOX and hit compounds identified from the primary screen, a secondary PDTO screen was performed. Given that abrogation of cell cycle progression was implicated in PDTOs resistance to FOLFOX, combined with the observation that a significant proportion of compounds identified as hits were compounds targeting cell cycle progression, 6 compounds from the primary screen targeting this process were selected for further analysis (alisertib, barasertib, AMG-900, milciclib, PHA-767491, dinaciclib). In addition to the 3 PDTOs used for the primary screen, one of the most FOLFOX-resistant PDTOs, yP315_LT, was also subjected to testing.

A total of five concentrations of each drug were tested alone or in combination with 6.25, 12.5 or 25 μM FOLFOX. To assess whether any of these drugs acted synergistically with FOLFOX in inducing cell death, we applied the Bliss reference model, where negative synergy scores indicate antagonism, scores around zero additivity and positive scores indicate synergy^32^ **(Figure 6b)**. While some heterogeneity was detected in the drug response of these 4 PDTOs, 4 of the 6 tested drugs exhibited clear synergy with FOLFOX across all patients, respectively targeting Aurora A (alisertib), Aurora B (barasertib), CDK2 (milciclib) and cdc7/CDK9 (PHA-767491) kinases **(Figure 6c)**. To ensure robustness of these results, the Loewe synergy reference model was also applied to the data, confirming the synergy of several compounds affecting the cell cycle with FOLFOX in 4 different PDTOs **(Suppl. Figure 11b)**.

Collectively, results from this large kinase inhibitor screen performed on PDTOs generated from FOLFOX-resistant CRLM identified multiple candidate compounds for targeted monotherapies. Additionally, we have identified the potential for combination therapy with compounds targeting cell cycle progression and FOLFOX, that may both improve response rates, but also allow for lower doses of FOLFOX to be administered to patients, thereby limiting toxicity and improving chemotherapy completion rates. Across all bioactive compounds, there was enrichment of those targeting progression through the DNA-damage response pathway, with those targeting the G_1_/S and G_2_/M cell cycle transitions among the most promising candidates, highlighting the ability of our approach to identify tool compounds with related biological effects.

### G_1_/S and G_2_/M cell cycle checkpoint inhibitors efficiently target FOLFOX-resistant liver metastasis PDTOs

Results from our PDTO drug screen suggested that compounds targeting kinases involved in p53-independent DNA damage response - particularly inhibitors of cell cycle checkpoint regulators such as CHK1/2, Aurora kinases A/B and CDK2, may be efficient to target metastatic tumors that progressed through FOLFOX treatment. In addition, in the most resistant PDTOs we identified a reduced progression through S phase and G_2_/M transitions, along with an activation of the SAC as key mechanisms of resistance to FOLFOX in metastatic PDTOs. These pathways collectively form the “*Mitotic DNA Damage Checkpoint*” shown to facilitate mitotic arrest in response to DNA damage so as to allow DNA repair^33^. To determine if targeting this checkpoint improved the sensitivity of resistant PDTOs to FOLFOX, we exposed the two most FOLFOX-resistant PDTOs, yP315_LT and P275_LT, to a 72-hour treatment with FOLFOX alone or combined with increasing doses of clinically relevant inhibitors of the Mitotic DNA Damage Checkpoint. Firstly, to target the G1/S component of this checkpoint, we selected the CHK1 inhibitor SRA737 (CCT-245737) and the Wee1 inhibitor adavosertib (AZD1775). Secondly, to target the G_2_/M arrest observed in resistant organoids upon withdrawal of FOLFOX, we used the MPS1/TTK inhibitor empesertib (BAY1161909). In TP53-independent responses to DNA damage, CHK1 plays a key role as regulator of fork stability, DNA repair and transcription through G_1_/S and G_2_/M transitions^***34***^. Wee1 acts downstream from CHK1 to regulate progression through G_1_/S and G_2_/M checkpoints, while MPS1/TTK is a master regulator of the SAC^*35*^. All three inhibitors have been used in the clinical setting, and expression of RNAs encoding all three of these kinases was found to be elevated in resistant PDTOs after FOLFOX exposure, as well as in tumor samples collected from patients with PD under neoadjuvant FOLFOX (**Figure 7a**).

**Figure 7.**
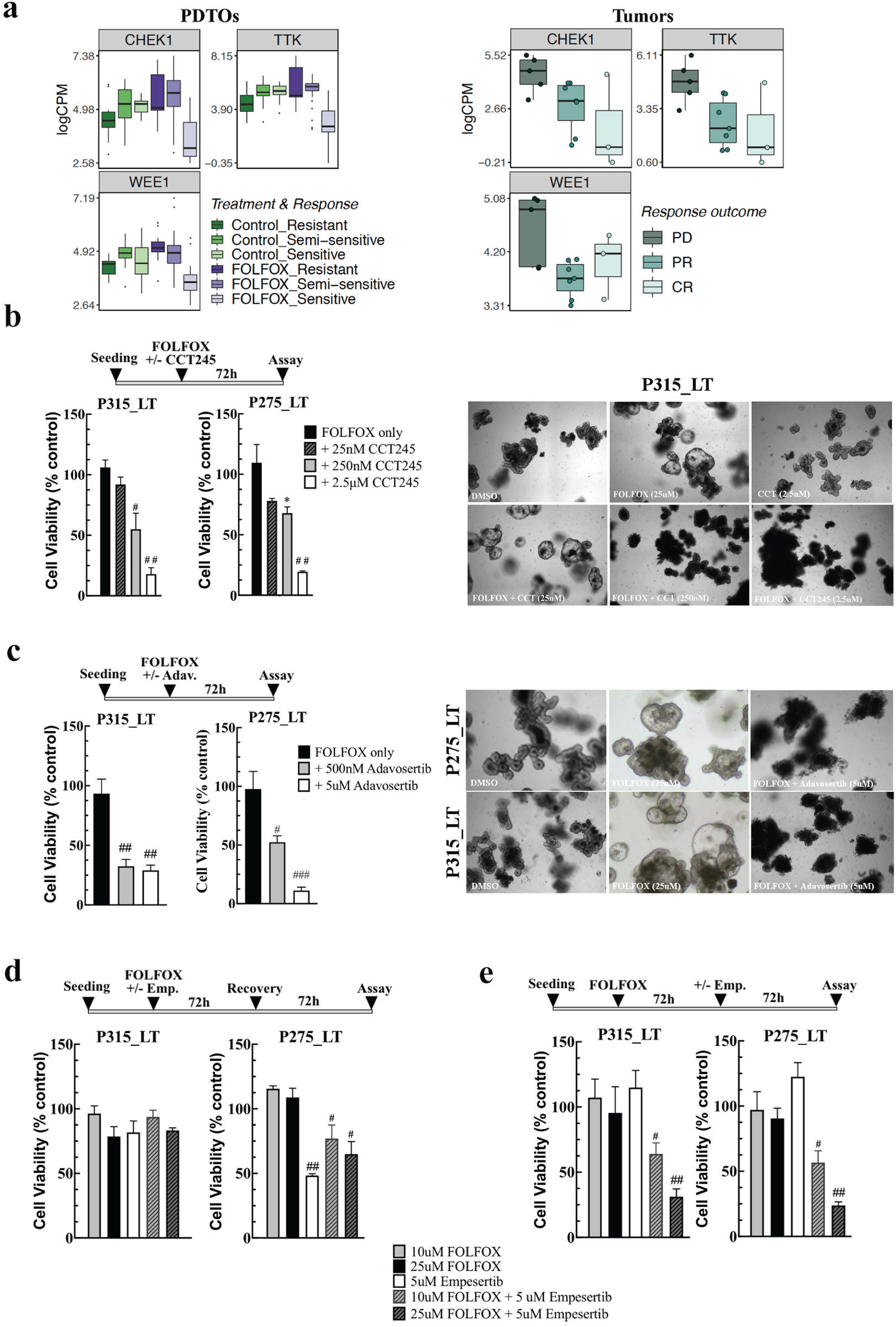
Inhibiting Master Regulators of the Mitotic DNA Damage Checkpoint Sensitizes Resistant PDTOs to FOLFOX. **a)** Boxplots comparing the expression level of the CHEK1, MPS1/TTK and WEE1 genes in vehicle or FOLFO-treated PDTOs stratified according to their response to FOLFOX *in vitro* (*left panels*) and in tumor samples collected from patients treated with neoadjuvant FOLFOX, stratified according to their treatment response (*right panel*) **b)** Bar graph (*left panel*) and bright field images (*right panel*) illustrating the strongly reduced cell viability of FOLFOX-resistant PDTOs following a 72-hour treatment with a combination of 25uM FOLFOX and of the indicated dose of the CHK1 inhibitor CCT-245747. **c)** Bar graph (*left panel*) and bright field images (*right panel*) illustrating the strongly reduced cell viability of FOLFOX-resistant PDTOs following a 72-hour treatment with a combination of FOLFOX and of the indicated dose of the WEE1 inhibitor adavosertib. **d)** Bar Graph illustrating the weak sensitivity of FOLFOX-resistant PDTOs to co-treatment with the MPS1/TTK inhibitor empesertib. **e)** Bar graph illustrating the strongly reduced cell viability of FOLFOX-resistant PDTOs following a sequential treatment with FOLFOX (72h) followed by the indicated dose of the MPS1/TTK inhibitor empersertib for a further 72h. (*, p < 0.05; #, p < 0.005; ##, p < 0.001, ###, p < 0.0005 vs FOLFOX alone, 2-way ANOVA with Dunnett post-hoc test, n = 3 repeats with 3 replicate wells per condition).

The combination of both adavosertib and CCT-245737 with FOLFOX resulted in sensitization of the resistant-PDTOs. Using the Bliss model, CCT-245737 and adavosertib showed indisputable synergy when combined with FOLFOX **(Figure 7 and Suppl. Figure 13a-d)**^32,36^. In contrast the MPS1/TTK inhibitor empesertib did not efficiently inhibit the survival of resistant PDTOs when used concomitantly with FOLFOX (**Figure 7d**), consistent with our prior observation that FOLFOX-resistant cells are predominantly arrested during the G_1_/S transition while under treatment. Given that resistant cells were shown to accumulate in G_2_/M shortly after FOLFOX withdrawal, we reasoned that administering empesertib during that post-treatment 72-hour window may increase its efficacy. Indeed, we found that a 72-hour treatment with 5 μM empesertib immediately following 72 hours of FOLFOX exposure was very effective in reducing the survival of FOLFOX-resistant PDTOs, by up to 90% (**Figure 7e and Suppl. Figure 13e**). Together these results i) indicate that clinically relevant targeted inhibitors of kinases that regulate the G_1_/S and G_2_/M checkpoints are highly efficient in killing FOLFOX-resistant CRLM PDTOs and ii) confirm the respective importance of sequential G_1_/S and G_2_/M blocks during and after FOLFOX exposure in the ability of FOLFOX-resistant PDTOs to bypass treatment **(Suppl. Figure 14)**.

## DISCUSSION

Despite improvements in treatment over the last few decades, the survival of patients with mCRC remains unacceptably poor. Understanding the mechanisms of resistance contributing to failure of standard chemotherapy administered in patients with mCRC is vital, both to improve patient survival and avoid the associated debilitating toxicity in patients who are unlikely to derive benefit from treatment. This study aimed to identify mechanisms of resistance to FOLFOX, the most frequently prescribed regimen in patients with stage II and IV CRC^13^. While various mechanisms have been identified for oxaliplatin or 5-FU monotherapy (including proteins involved in drug uptake, export and metabolism, modulation of thymidylate synthase expression, or alterations of apoptosis/survival effectors, as reviewed), there are few preclinical studies investigating resistance to 5-FU/Oxaliplatin combinations^37^. Among those, previous studies have suggested that *ERCC1* over-expression, leading to increased DNA damage repair through the nucleotide excision repair pathway, correlates with FOLFOX resistance^38,39^.

PDTO models represent an ideal model to identify and characterize chemotherapy resistance mechanisms, as their genomic and phenotypic features recapitulate those of tumors they originate from, and because their response to treatment is frequently similar to responses in patients^**13–15,40–45**^. To date, although oxaliplatin is not clinically given as monotherapy^15,46^, previous studies have largely assessed the impact of oxaliplatin or 5-FU on PDTO survival as part of larger drug panels^47,48^, and the concordance between PDTO responses and patient response to FOLFOX remains unclear. In this study we have identified differential responses to FOLFOX in a panel of mCRC PDTOs, that correspond to FDG-PET responses of matching CRLM in patients. Additionally, we describe for the first time a FOLFOX-resistance gene expression signature in mCRC PDTOs and demonstrate that this signature is enriched in CRLM that progress under neo-adjuvant FOLFOX treatment in patients.

Resistant PDTOs exposed to FOLFOX ceased to divide and accumulated in early S-phase, as demonstrated using a combination of phospho-Histone H3 immunostaining and flow cytometry cell cycle analyses. Following FOLFOX withdrawal, surviving cells moved to G2M but did not resume proliferation, allowing for DNA damage to be repaired. Importantly this process occurs in the absence of functional *TP53,* which was mutated in these samples. RNAseq analysis indicated that FOLFOX-resistant PDTOs, as well as CRLM that progressed under neoadjuvant FOLFOX treatment, were characterized by high expression of genes involved in the E2F pathway (primarily E2F1 and E2F4) and in *TP53* independent cell cycle arrest including S-phase, G_2_M and spindle assembly checkpoints, compared with FOLFOX sensitive PDTOs.

In this study, the combination of RNAseq analyses, large scale PDTO drug screen and secondary screen indicate that abrogation of cell-cycle progression is implicated in FOLFOX resistance, and that agents targeting specific cell cycle checkpoints synergize with FOLFOX to overcome resistance. Inhibition of these checkpoints may be particularly effective in colorectal tumors exhibiting features of the chromosomal instability (CIN) pathway, recently shown to represent 85% of sporadic CRC in consensus molecular subtype analyses^49,50^. Overexpression of the spindle assembly complex as well as frequent mutations of the DNA damage response (DDR) pathway genes are associated with CIN^51,52^, and there is increasing evidence of a more formal pathway linking the DNA damage response and SAC in controlling mitotic progression, via a “*Mitotic DNA Damage Checkpoint*”^33^. The “*Mitotic DNA Damage Checkpoint*” is thought to be independent of a functional kinetochore and is an overlap of the DNA damage (or G_1_/S) checkpoint and of the SAC, converging through their inhibition of PDS1^53^.

Our study demonstrates that specific targeting of several key “*Mitotic DNA Damage Checkpoint*” regulators including CHK1 (CCT-245737), Wee-1 (adavosertib) and MPS1/TTK (empesertib) results in sensitization of previously resistant PDTOs to FOLFOX treatment. Importantly, the timing for optimal efficacy of these inhibitors corroborates our observation of a dual process of reversible cell cycle arrest in FOLFOX-resistant cells. Indeed, inhibitors able to release the G1-S checkpoint (CCT-245737 and adavosertib) are effective when combined with FOLFOX, whereas empesertib is mostly effective as a sequential treatment to induce elimination of tumor cells that have escaped death and are arrested in G2/M whilst repairing their DNA. Collectively these results indicate that compounds targeting these checkpoints offer a promising avenue for the treatment of patients with colorectal cancer unresectable liver metastases, for whom therapeutic options are extremely limited and survival outlook is very poor. In addition, the compounds validated here are showing clear synergy with FOLFOX, suggesting that combination treatment with such inhibitors may enable clinicians to reduce FOLFOX dosage. As neurotoxicity from FOLFOX is correlated with cumulative doses of oxaliplatin, this would significantly improve treatment tolerability and thereby therapeutic completion rates. Finally, given the majority of colorectal cancers have CIN underlying their genomic instability, therapies targeting the Mitotic DNA Damage Checkpoint have the potential for widespread application.

In summary we have identified a novel FOLFOX resistance signature in mCRC, derived from PDTOs and validated in tumor samples from patients that progressed on neoadjuvant FOLFOX based chemotherapy. Additionally, we have shown synergy with agents targeting cell cycle progression with FOLFOX. Specifically, compounds inhibiting TP53 independent cell cycle arrest and the SAC have been shown to be highly efficient in inducing cell death in FOLFOX resistant PDTOs. Our results thus suggest that CHK1, WEE1 and MPS1 inhibitors represent promising candidates that warrant further preclinical characterization to define optimal conditions for their use and toxicity profile in combination with FOLFOX for metastatic colorectal cancer.

## Acknowledgements

The authors are very grateful to staff from the University of Melbourne imaging (Paul McMillan and Ellie Cho) and Flow cytometry (Vanta Jameson) platforms, the Peter MacCallum Cancer CAHM (Centre for Advanced Histology and Microscopy) and the Molecular Genomics Core for their technical help during completion of this work. We also wish to acknowledge funding from the CSSANZ Foundation Grant (AGH and CB), the PMCC Foundation Grant (AGH and CB), Covidien Grant (CB), the Tour de Cure Foundation (Senior Research Grant, FH) and the NHMRC of Australia (Project grants #GNT1164081, FH). We also thank Mr. Jamie Keck, Mr. Michael Johnston, Mr. Adrian Fox, Mr. Simon Banting, Mr. Jacob McCormick, Mr. Satish Warrier, Mr David Chan, Mr. Glen Guerra, and Mr. Joseph Kong for facilitating sample and reagent access and for helpful discussions. The Victorian Centre for Functional Genomics (K.J.S.) is funded by the Australian Cancer Research Foundation (ACRF), Phenomics Australia (PA) through funding from the Australian Government’s National Collaborative Research Infrastructure Strategy (NCRIS) program, the Peter MacCallum Cancer Centre Foundation and the University of Melbourne Research Collaborative Infrastructure Program (MCRIP). We acknowledge Compounds Australia (Griffith University) for their provision of specialized compound management and logistics research services to the project.

## METHODS

### PREPARATION OF ORGANOIDS

#### Sample collection

Colorectal liver metastasis and where possible matched primary tumors were collected from patients undergoing treatment at Peter MacCallum Cancer Centre, The Royal Melbourne Hospital, St Vincent’s Hospital, St Vincent’s Private Hospital and Morwell hospital. The study was approved by the ethics committee at Peter MacCallum Cancer Centre (15/169) and written informed consent was obtained from all patients. For establishment of organoids, tumors approximately 0.25-1cm^3^ were collected either from the operating theatre or directly from the pathologist.

#### Organoid culture

Tumors were washed prior to processing with DMEM/F12 and a combination of gentamycin, penicillin/streptomycin, nystatin and amphotericin to ensure broad spectrum cover of enteric bacteria and fungi. Under sterile conditions the normal tissue was separated from the tumor. The tumor was then cut into smaller fragments using a sterile scalpel. Tumor fragments were then enzymatically digested using the Human Tumor Dissociation Kit (Miltenyi Biotec^R^), followed by homogenization with the Gentle Macs Rotator^R^ (Miltenyi Biotec^R^). The digested and homogenized tumor was incubated for a total of 20 minutes at 37C. Digested samples were then passed through a 70um filter. Smaller filter sizes were not used so as to maintain the tumor in clusters of cells, rather than single cells, to improve growth. Tumor clusters were then resuspended in 100% Matrigel (Corning), 40ul per well (24 well). The Matrigel^R^ was then allowed to set for a minimum of 10 minutes in the incubator and then 500 ul of warm organoid media as added to the Matrigel dome (**Suppl. Table 9).**Organoid cultures were initially grown in hypoxic conditions (2% oxygen), then transferred to normoxic conditions (20% oxygen) after 5-7 days. Growth media was changed twice per week.

#### Passaging of organoids

Organoids were passaged once they were macroscopic. Cold PBS was added to each of the wells and then to preserved heterogeneity wells were combined and collected in a 15ml tube and centrifuged at 1400 rpm for 5 mins. Tryple^R^ (Gibco) enzyme solution was then added in the volume of 10x the amount of Matrigel^R^ and organoids broken down manually with 1ml pipette. The 15ml tube was then placed in 6-minute intervals into the incubator and checked to ensure dissociation into fragments, with the average time being 6-12 minutes. Cold PBS was then used to dilute the Tryple^R^ and centrifuged again at 1400 RPM for 5 mins. Matrigel^R^ was then added to resuspend the organoid fragments. Fast growing organoids were diluted 1:3 upon passaging, while slower growing ones were diluted 1:2.

#### Immunohistochemistry

To prepare organoids for fixation, media was removed from the well and a solution of 4% Paraformaldehyde and 0.5% Glutaraldehyde was instilled into the well (500 μl for 24 well plate) to fix the whole Matrigel pellet with the organoids embedded, for 20 minutes at room temperature. The fixation solution was then aspirated and washed with PBS. 70% Ethanol solution was then instilled into the well and the firm Matrigel^R^ pellet transferred into a solution of 70% ethanol. This pellet was then embedded in paraffin. Paraffin embedded tissue and organoid sections were dewaxed in a Jung XL auto-stainer and antigen retrieval performed in 10 mM citrate buffer at 125C for 15 minutes in a pressure cooker. Endogenous peroxide blocking was performed in 1% water for 15 minutes before blocking in 2.5% normal horse serum (Vector Laboratories -MP-7401) for 1 hour. Incubation with the primary antibody was performed overnight at 4C in PBS containing 3% BSA/0.5% Tween 20. Primary antibodies were mouse anti-Cytokeratin 7 (DAKO - M7018, 1:400), Rabbit anti-Cytokeratin 20 (Abcam - ab76126, 1:400), Rabbit anti-phospho-Histone H3 (Ser10) (Cell Signaling – 9701, 1:400), Rabbit anti- Caspase-3 (cleaved) (Abcam – ab13847, 1:4000).

Incubation with the secondary antibody (ImmPRESS HRP Anti-Rabbit IgG (Vector Laboratories - MP-7401) and Anti-Mouse IgG (Vector Laboratories - MP-7402) (Peroxidase) Polymer Detection Kits) was performed for 30 minutes at room temperature. The chromogen 3,3’-Diaminobenzidine (ABACUS DX - SK-4100) was used to visualize the staining before the slides were counterstained with hematoxylin.

#### γH2AX Immunofluorescent staining

Organoid sections were dewaxed, and antigen retrieval was performed as per the IHC samples. Following antigen retrieval, the slides were blocked with 3% BSA in PBS + 0.5% Tween 20 for 1 hour. The sections were then incubated in the primary antibody (Mouse anti-gamma-H2AX (phospho S139) (Abcam - ab22551, 1: 1000) overnight at 4C. The slides were then exposed to the secondary antibody (Goat anti-Mouse IgG (Alexa Fluor 568) (Molecular Probes - A-11031) and to a DAPI counterstain for 30 minutes at room temperature before being cover slipped and sealed. Imaging was performed using the 60x objective of a Nikon C3 confocal microscope. Final images are presented as Z-projections prepared from a minimum of 6 1um z-stack images.

#### LEGO Viral Transduction and assessment of Organoid Heterogeneity

Organoids dissociated into single cells/small clusters using Tryple^R^. Three wells of single cells combined together in one ultralow adherent 24 well plate in 500 ul of organoid media. Fluorescent lentiviral gene ontology (LEGO) vectors added in volumes dependent on pre-calculated viral titre. These included eBFP, Sapphire, eGFP, Venus, mOrange and dKatushka. After 24 hours virus was washed away from the tumor cells and they were then resuspended in Matrigel. Once macroscopic organoids were passaged as described above and seeded in increasing cellular density of 10,000, 20,000 and 200,000 cells. Organoids were again cultured until macroscopic and then imaged with Olympus FV3000 to determine the level of heterogeneity following single cell vs whole organoid seeding approaches **(Suppl. Figure 5a)**.

### CYTOTOXIC STUDIES

#### FOLFOX Treatment and Targeted Compounds

Organoids were cultured until macroscopic in 24 well plates with 40 μl of Matrigel^R^ **(Suppl. Figure 5b)** Organoids were seeded the day prior to cytotoxic treatment (minimum 16 hours). Cold Organoid Harvesting Solution^R^ (Cultrex) was added to each well and the Matrigel^R^ dome very gently dissociated with a 1ml pipette. The whole 24-well plate was then placed in the cold room for 1 hour (4-8C). The contents of the wells were then collected in a 15ml tube, cold PBS added and then centrifuged at 1400 rpm for 5 minutes. The organoids were resuspended in new Matrigel^R^ to the desired density, usually 1:4. 10 μl of Matrigel^R^ per well was instilled using a cold 20 μl pipette. Once the Matrigel^R^ dome had set, 100 μl of warmed Organoid Media was then added to each well. Chemotherapy was then added in a volume of 20 μl volume to make a final concentration of 25 μM of oxaliplatin and 25 μM of 5-Fluorouracil. Organoids were treated for 72 hours at and imaged at D0 and D3. Cell Titre Glo (CTG) 3D^R^ was (Promega) used to determine viability. The CTG was left to equilibrate to room temperature overnight. Plates were then removed from the incubator and left to equilibrate at room temperature for 30 mins. CTG was then added at a ratio of 1:3 (e.g. 50 μl for 100 μl media), the plate vigorously shaken on a plate shaker at 180 pm for 20 minutes and the result read on the plate reader (Perkin Elmer EnSpire). There were 5 technical replicates of each treatment and control well in each experiment, and experiments were independently repeated at least 3 times for each organoid. All organoids undergoing cytotoxic treatment tested negative for mycoplasma.

#### Cell-cycle FACS Experiments

Matrigel was dislodged with cold PBS and transferred into 1.5ml tubes, spun down (5mins, 1400rpm) and PBS removed. 500 μl of Tryple^R^ was added and tubes were incubated on the orbital shaker at 37°C for 40 minutes, pipetting every 20 minutes to break up organoids. The resulting single cell suspension was fixed with ethanol dropwise (70% in filtered milli-Q H_2_O) whilst vortexing and left for 30 minutes at 4°C. The fixed cells were spun down (5mins, 1400rpm), ethanol removed and 100 uL of the PI solution was added (500 ml FACS buffer, 1 μl ribonuclease and 20 μl PI) followed by a 30-minute incubation on ice before FACS.

#### Primary Organoid Screen

Organoids were thawed approximately 2 weeks prior to treatment date, grown until macroscopic and passaged 2-3 times prior to treatment. A fresh batch of organoids was thawed for each separate week (4 screening weeks in total) to ensure that organoids were cultured for the same time prior to each treatment. A consistent batch of Matrigel was used for the entire experiment (LOT 8204010). A total of 6 highly confluent 40 μl wells of a 24-well plate were required to seed each 96 well plate. Seeding was performed using the above-described protocol, one day prior to treatment. Compounds for treatment were provided by Compounds Australia lyophilized at 2 concentrations 5 μl and 500 nM (96 well format, **Suppl. Table 8 - *Sheet 1*)**. Positive and negative controls were added on the day of screening to the outer wells in multiple replicate wells. The drug plates were then hydrated with organoid media **(Suppl. Table 9)** and the compounds added to plates using the then kinase compounds added by the Sciclone ALH3000 robot (Perkin Elmer). Organoids were imaged using the Cytation5 (BioTek) on D 0,1, 2 with bright field (BF) and on D3 with BF and the viability stain 7-Aminoactinomycin D (7-AAD, 4x magnification, 8 fields total). The CTG result was then obtained from the same wells, using the protocol described above.

The heat maps for the raw CTG values were checked for edge effects and obvious seeding/instrument issues. During the QC step we identified, 6 compounds in patient 1 and 5 compounds for patient 2 that had low seeding densities and were removed from further consideration. There were no seeding issues for patient 3. The raw CTG values were normalized to the mean of the DMSO control wells on a per-plate basis and the Fold Change (FC) calculated as Percent FC (PercFC) with DMSO being at 100%. Control quality control (QC) metrics were then calculated including the mean, standard deviation and from these the percentage coefficient of variation (%CV) of negative and positive controls and the Z’factor **(Suppl. Table 10)**.

The %CV for the three organoid DMSO controls were 13.4% (Patient 1, yP516_LT1), 27.6% (Patient 2, yP520_LT1) and 10.3% (Patient 3, yP295). Despite patient 2 slight %CV slightly outside the acceptable range (24%), all images were individually checked to confirm viability in BF and 7AA manually, including checking for low CTG due to seeding issues. Patient 2 organoids were known to have a slower growth rate than the other 2 patients which may have contributed to the variation. In all three patients, the positive controls were very variable, which was to be expected for toxic treatments and a 3D setting. The Z’ Factor was also calculated to assess the degree of overlap between each DMSO and positive control (FOLFOX 250nM) pair. Patients 1 and 3 showed minimal overlap between controls, with Patient 2 showing some overlap. The formula used to calculate Z’ Factor:

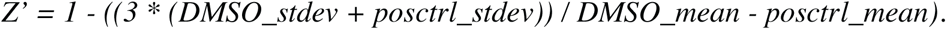

Further QC was ensured by comparing the CTG viability of the controls to demonstrate the variability and distribution of replicate control wells, displayed with a notched box plot. The values shown are cell counts normalised to the DMSO mean on a per-plate basis. The notch displays a 95% confidence interval around the median (based on the median +/− 1.58*IQR (inter-quartile range)/sqrt(n)). Additionally, analysis with *CellProfiler* confirmed the spheroid area and count were uniform across all controls, and that the percentage dead area was as expected in negative and positive controls **(Suppl. Figure 15**).

To identify hits from the screen, the CTG values were normalized to DMSO and the compounds were assigned into bins corresponding to their ability to induce cell death. Z-scores were calculated to determine statistically significant changes compared to DMSO controls. A strict cut-off for hits was applied to the compounds. Only compounds that had a PercFC less than or equal to 50% AND a significant Z’score less than or equal to −3 were considered HITS. Additionally, to find compounds that show the most overlap in their effect on the different patient cell types, the percentage CTG (PerFC) was assigned to bins:

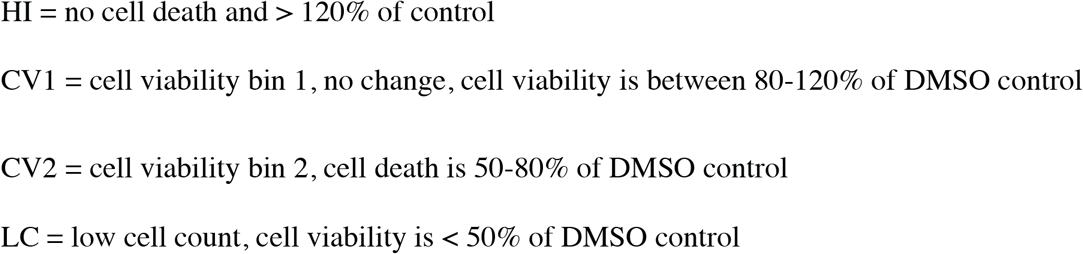

Finally, identification of hits was corroborated using multi-dimensional perturbation(mp) and Mahalanobis distance calculations from the imaging data whereby mp-values < 0.05 were considered as significant and Mahalanobis >= 2 as a change that is not achieved in any of the untreated or DMF controls. Compounds meeting these cut-offs were classified as active. Positive and negative controls were removed from the active compound dataset^31^.

#### Secondary Organoid Screen

The three PDTOs from the primary screen (yP516_LT1, yP520_LT1, yP295_LT) and an additional resistant organoid were used for the secondary screen. A total of six compounds were chosen from the primary screen including alisertib (AUKA inhibitor), barasertib (AUKB), AMG-900 (PanAUK inhibitor), milciclib (CDK2), PHA-767491 (CDK7/9 inhibitor) and dinaciclib (CDK1/5). A total of 5 concentrations of each drug, were combined with 6.25 uM, 12.5um and 25uM of FOLFOX, with two technical replicates and a control plate with kinase compounds only.

Similar to the primary screen a number of QC analysis were performed with the CTG value of each well normalised to the mean of the DMSO wells on a per-plate basis and the heat maps of the plates checked for edge effects and obvious seeding/instrument issues. The mean, standard deviation (stdev), and the % cv for the DMSO and FOLFOX controls, and Z’ Factor for each positive and negative pair were calculated (**Suppl. Table 10**). The kinase drug plates were prepared, hydrated and dosed onto organoids with the Sciclone ALH3000 robot (Perkin Elmer) as above. FOLFOX was then added subsequently within 1-hour post dosage of kinase drugs and organoids were then co-treated for 72 hours. Organoids were imaged on Day (D) 0, 1, 2 with bright field (BF) and on D3 with BF. The CTG result was then obtained using the protocol described above.

To assess whether two drugs act synergistically in inducing cell death, we applied the Bliss and Loewe reference model **(Figure 6b and Suppl. Figure 11b)^32,36^**. The visualization of the synergy scores is conducted as a two-dimensional and a three-dimensional interaction surface over the dose matrix using the synergyfinder R package. Negative scores indicate antagonism, scores around zero indicate additivity, and positive scores indicate synergism.

### EXTRACTION, LIBRARY PREPARATION and SEQUENCING of RNA and DNA

#### Extraction - Tumor Samples

Tumor samples collected directly from the patient’s surgical specimens were immediately transported to the laboratory for processing. Prior to preparing organoids a small sample of tumor and adjacent normal tissue (liver or colon depending on specimen) was separated and immediately frozen with dry ice. In patients where it was not possible to obtain normal tissue germline DNA was extracted from blood. DNA and RNA were then extracted from the same sample of tissue using the Qiagen AllPrep Mini Kit (Qiagen, #80204). In two samples RNA was extract using Trizol reagent (Invitrogen #155906026).

#### Extraction - Organoid samples

PDTOs were isolated from the Matrigel^R^, using Cultrex^R^ Organoid Harvesting Solution (#370010001) and placed on ice for 1 hour to remove any residual Matrigel which may contaminate the sample. Samples were then centrifuged and washed with PBS before pelleting and snap freezing. RNA and DNA was then extracted as above Qiagen AllPrep Mini Kit (Qiagen, #80204). Those PDTOs that underwent FOLFOX treatment (n=14) had RNA extracted using the Qiagen RNAeasy micro kit (Qiagen, # 74004).

#### Library Preparation and Sequencing – RNA-seq

There were two separate batches of RNA due to the high number of samples. A number of samples were duplicated across the two runs to allow examination of batch effects. Both libraries were prepared using the QuantSeq 3′ mRNA-Seq Library Prep Kit FWD from Illumina, following the standard protocol. RNA concentrations were quantified using RNA HS QuBit Reagents. Samples were combined in an equimolar pool and the 75bp long Single-End reads were generated using a NextSeq with a High Output 75 cycle kit.

##### Run MGC-CB-1345

150-300ng total RNA loaded per sample (most samples with 300ng). 16 PCR cycles, final library concentrations between 0.1-9.5ng/ *μ*L and final library product sizes (avg) 256-381bp.

##### Run MCG-CB-1427

100-1000ng total RNA was loaded per sample, the majority with 1000ng. 14 PCR cycles with the final library concentrations between 0.72 – 10.8 ng/ *μ*L and the final library product sizes (avg) 303–446 bp.

#### Library Preparation and Sequencing - Whole Exome Sequencing

DNA concentration was measured using the Qubit dsDNA HS Assay Kit (ThermoFisher). 150-300 ng of DNA was fragmented to approximately 200 bp using a focal acoustic device (Covaris S2, Sage Sciences). Libraries and hybridization capture were performed following the SureSelectXT HS recommended protocol with SureSelect Human All Exon V6+UTR baits (Agilent). Indexed libraries were sequenced on an Illumina NovaSeq with the S4 flow cell (Illumina) to generate on average 120 million paired-end 150 bp reads per sample. Extracted DNA was sheared using the Covaris M220 Focused-ultrasonicator with a target fragment length of either 350bp or 550bp through bead size selection. The Illumina TruSeq nano DNA library preparation kit was used for End repair and Adenylation of 3’ fragment end. The adaptor was ligated and the molecular barcode/index incorporated through amplification. The library was checked to ensure quality (Qubit, Bioanalyzer and KAPA Illumina library quantification kit using qPCR) prior to normalization and pooling before loading onto the Illumina NovaSeq 6000 for sequencing following Illumina’s Sequencing by Synthesis technology. Each image was converted to a BCL file by Illumina software Sequencing Analysis Viewer. Upon completion of sequencing, each BCL file was converted to a FASTQ file using Illumina default pipeline for further analysis including alignment and annotation.

## COMPUTATIONAL & STATISTICAL ANALYSES

We used R version 3.6.1 to perform data analysis and visualizations. The versions and citations for all statistical methods or packages used in this paper are provided in Suppl. Table 11. Details of applying different method are given in the corresponding sections. For general data wrangling and visualization, we used the base R functions as we all as several core packages from the tidyverse R package, such as tidyr (v 1.0.0), dplyr (v 1.0.0), and readr (v 1.3.1) for reading and manipulating the data, and ggplot2 (v 3.2.1) for data visualization. Heatmaps were generated using the custom functions generated based on the complexHeatmap R package (**Suppl. Table 11**).

## Analysis of the RNA- seq Data – PDTOs and Tumor Samples

### RNA-seq data analysis and DEG derivation in PDTO samples

The strand specific (forward) 3’ RNA-seq reads were aligned using HISAT2 against human genome GRCh37 release 75 version of Ensembl. We used featureCounts from the RSubread package to quantify the number of gene-level reads for each gene per individual samples. For the RNA-seq data from both PDTO and tumor samples, we examined the consistency between replicates using correlation and mean-average plots, and further examined the quality of the data using relative log expression (RLE) plots **(Suppl. Figure 16)** and principal component analysis (PCA) plots (**Suppl. Figure 17)**. We used the filterByExpr function from the edgeR package to filter lowly expressed genes and calculated count-per-million (CPM) values. Differential expression analysis was performed using the voom-limma pipeline.

The PDTOs were sequenced in two separate sequencing runs (as detailed above), and therefore we included “Run” as an extra term representing putative batch effects in our model.matrix. We also used the duplicateCorrelation function from limma to calculate correlation between replicates by considering replicated samples as a blocking variable. The consensus correlation between replicates as well as the blocking variable “sample” were used when running the voom function and fitting linear model using the lmFit function. After running eBayes, we considered genes with absolute log2FC > 1 and adjusted p-value < 0.05 as differentially expressed genes (DEGs).

### RNA-seq data analysis and DEGs derivation in tumor samples

For the tumor data, all samples were sequenced together and therefore we did not have known batch effects in the data; however, as we had one outlier sample, we used voomWithQualityWeights function, whose results were used in duplicateCorrelation function when calculating the correlation between samples from the same patient by considering patient variable as the blocking variable. We then ran voomWithQualityWeights again, this time by providing the consensus correlation between samples from the same patient as well as the blocking argument (i.e. patients), followed by fitting of linear models and empirical bayes provided in the limma package. We considered genes with absolute log2FC > 1 and adjusted p-value < 0.05 to define DEGs between PD vs CR, PR vs CR, and PD + PR vs CR. We used a more stringent threshold (absolute log2FC > 2 and adjusted p-value < 0.01) to define DEGs between PD vs PR and PD vs CR + PR due to large differences in the transcriptional profiles of these subsets.

### Consistency between tumor and PDTO samples using RNA-seq data

The count data from the PDTO vehicle treated samples were combined where we had replicated samples for the same PDTOs. There were 18 pairs of tumors and PDTOs after removal of the PC002_CoT samples due to its low quality. For each pair, we combined the count data, and filtered genes to retain only those with CPM > 1 in at least one of the samples in the pair. Then, we calculated the logRPKM values sung the rpkm function in the edegR package, followed by the Spearman’s correlation as well as the log-ratio of the expression values.

### Gene set testing and pathway analysis

Taking the DEGs from each of the comparisons, we used goana and kegga from the limma package to perform gene ontology (GO) and KEGG pathway enrichment analyses^54^. We also applied camera gene set testing with the MSigDB signatures retrieved from the msigdf R package^55^.

### Single-sample scoring

We used the singscore R package to score samples against established molecular signatures and quantify concordance between the transcriptome of each sample with the given signatures^21^. This approach is an unsupervized, rank-based method that does not need a list of DEGs or group labels, but instead calculates the normalized averaged mean-ranks of signatures genes within individual samples. This results in a signature score for each sample with a higher score associated with higher concordance of a sample to a given signature. Depending on gene set directionality, we used different settings of the simpleScore function as specified within the package documentation.

Signatures used with singscore were collected from different sources, with the majority of the signatures were taken from the Hallmark, C2 and C6 categories in MSigDB^20^. All the cell cycle, spindle, and E2F related signatures were taken from MSigDB by searching for associated key words. Other signatures were extracted from the literature: DNA damage repair genes from Wood et al^30^, the TP53-negative signature from Markert et al ^56^, pan-cancer stromal and immune signatures from Aran et al^57^, an NK signature from Cursons et al^58^, a T cell signature form Jerby-Arnon et al^59^, and an interferon signature from the imsig R package^60^.

## Analysis of the WES and WGS Data – PDTO and Tumor Samples

### Variant calling and copy number variation analysis in WGS data

For WGS data (*N_WGS_* = 36), trimming of sequence reads was carried out using Cutadapt, they were aligned to the reference GRCh37 assembly using BWA-MEM, coordinate-sorted with Samtools and duplicate-marked with Picard. The average depth of read in WGS data for the tumors was 58.9X, organoids 36.2X and normal 31.8X. Single nucleotide substitution variants were detected using a dual calling strategy using qSNP and the Genome Analysis Toolkit (GATK) HaplotypeCaller. The GATK HaplotypeCaller was also used to call short indels of ≤50 bp. Quality filtering removed reads with less than 35 reference matched contiguous bases as indicated in the CIGAR string, more than 3 mismatches in the sequencing MD field, and a mapping quality greater than 10. High confidence variants were selected based on passing further variant characteristics including minimum depth of 8 reads in the control data and 12 reads in the tumour data; at least 5 variant supporting reads present where the variant was not within the first or last 5 bases; at least 4 reads with unique start positions; identification of the variant in reads of both sequencing directions; and not adjacent (more than 4 base pairs) to a mononucleotide run of 7 or more bases in length. We used Variant Effect Predictor (VEP; with Ensembl release v 90) to annotate the variants, and further filtered them to only focus on protein coding genes and variants with medium and high impact. ascatNgs was used to estimate CNV in the WGS data. The circos plots for the WGS data were generated using the Circos software.

**WES** data was processed using Seqliner WES pipeline (seqliner.org). Sequence reads from FASTQ files (*N_WES_* = 53) were trimmed with Cutadapt and aligned to the human reference genome GRCh37 using BWA-MEM. Aligned reads were sorted and indexed with samtools and duplicates reads were marked with Picard UmiAwareMarkDuplicatesWithMateCigar. The average read depth for the tumors was 119.6X, for organoids was 97.7X, and for the normal samples was 116.5X.

GATK was used to perform local realignment around indels and base quality score recalibration. Somatic SNVs/INDELs were called using VarDict, GATK MuTect2 and MuTect. Variants were annotated using VEP (with Ensembl release 90). We kept somatic variants called by at least two variant callers, which were high-quality calls with bidirectional reads. We further filtered to only focus on protein coding genes and variants with medium and high impact. Copy number aberrations and LOH were detected using FACETs.

When analyzing mutation data, from 89 DNA sequencing samples (36 WGS and 53 WES), we removed 11 samples due to low quality/purity, which resulted in 78 samples (30 WGS and 48 WES). We then consolidated all replicated samples resulting in 73 samples from 35 patients. The WGS and WES data, that were annotated and filtered in the same way, were integrated to generate a combined genomic data set. In cases, with more than one type of consequence annotated for the same mutation, we prioritized mutation severity from the Ensemble database (https://asia.ensembl.org/info/genome/variation/prediction/predicted_data.html). We then only focused on the mutations on the canonical transcript for further visualizations, and for a small subset of genes with more than one canonical transcript, we annotated their variants as multi-hits (MHT). For CNV analyses, as FACETS does not give information for chromosome Y, we separately examined genes on the X chromosome for male samples to make sure that we do not annotate them as deletions. Circos plots and piano plots of the CNVs were generated using the rock R package. A summary of the annotations used for the genomic analysis for tumor and found in **Suppl. Table 12.**

## Data & code availability

Sample annotations for the expression and mutation analyses are in different sheets of **Suppl. Table 12**. Expression read counts for tumors and organoids (obtained using featureCount) will be available from GEO (https://www.ncbi.nlm.nih.gov/geo/query/acc.cgi?acc=GSE165649) from the time of publication. The codes reproducing the results of this study are available from Github (https://github.com/HollandeLab/FOLFOX_Resistant_mCRC).

## Notes

https://github.com/HollandeLab/FOLFOX_Resistant_mCRC

### Competing Interest Statement

The authors have declared no competing interest.

## REFERENCES

1. Network, N.C.C. NCCN Guidelines Version 4.2020 Colon Cancer. Vol. 2020 (2020).

2. Australia, C. Relative survival by stage at diagnosis (colorectal cancer). Vol. 2020 (2019).

3. Rawla, P., Sunkara, T. & Barsouk, A. Epidemiology of colorectal cancer: incidence, mortality, survival, and risk factors. Prz Gastroenterol 14, 89–103 (2019).

4. Adam, R. & Kitano, Y. Multidisciplinary approach of liver metastases from colorectal cancer. Annals of Gastroenterological Surgery 3, 50–56 (2019).

5. Van Cutsem, E., et al. ESMO consensus guidelines for the management of patients with metastatic colorectal cancer. Annals of oncology: official journal of the European Society for Medical Oncology 27, 1386–1422 (2016).

6. Nordlinger, B., et al. Perioperative FOLFOX4 chemotherapy and surgery versus surgery alone for resectable liver metastases from colorectal cancer (EORTC 40983): long-term results of a randomised, controlled, phase 3 trial. The Lancet. Oncology 14, 1208–1215 (2013).

7. Alberts, S.R., et al. Oxaliplatin, fluorouracil, and leucovorin for patients with unresectable liver-only metastases from colorectal cancer: a North Central Cancer Treatment Group phase II study. Journal of clinical oncology: official journal of the American Society of Clinical Oncology 23, 9243–9249 (2005).

8. Giacchetti, S., et al. Long-term survival of patients with unresectable colorectal cancer liver metastases following infusional chemotherapy with 5-fluorouracil, leucovorin, oxaliplatin and surgery. Annals of oncology: official journal of the European Society for Medical Oncology 10, 663–669 (1999).

9. Takahashi, T., et al. Multicenter phase II study of modified FOLFOX6 as neoadjuvant chemotherapy for patients with unresectable liver-only metastases from colorectal cancer in Japan: ROOF study. Int J Clin Oncol 18, 335–342 (2013).

10. André, T., et al. Oxaliplatin, fluorouracil, and leucovorin as adjuvant treatment for colon cancer. The New England journal of medicine 350, 2343–2351 (2004).

11. Grothey, A.M.D., et al. Duration of Adjuvant Chemotherapy for Stage III Colon Cancer. The New England journal of medicine 378, 1177–1188 (2018).

12. Sasaki, N. & Clevers, H. Studying cellular heterogeneity and drug sensitivity in colorectal cancer using organoid technology. Curr Opin Genet Dev 52, 117–122 (2018).

13. Ooft, S.W.F., Dijkstra K, Mclean C,. Patient-derived organoids can predict respone to chemotherapy in metastatic colorectal cancer patients Science Translational Medicine 11(2019).

14. van de Wetering, M., et al. Prospective derivation of a living organoid biobank of colorectal cancer patients. Cell 161, 933–945 (2015).

15. Vlachogiannis, G., et al. Patient-derived organoids model treatment response of metastatic gastrointestinal cancers. Science (New York, N.Y.) 359, 920–926 (2018).

16. Cristobal, A., et al. Personalized Proteome Profiles of Healthy and Tumor Human Colon Organoids Reveal Both Individual Diversity and Basic Features of Colorectal Cancer. Cell reports 18, 263–274 (2017).

17. Network, C.G.A. Comprehensive molecular characterization of human colon and rectal cancer. Nature 487, 330–337 (2012).

18. Yaeger, R., et al. Clinical Sequencing Defines the Genomic Landscape of Metastatic Colorectal Cancer. Cancer Cell 33, 125–136.e123 (2018).

19. Pitroda, S.P., et al. Integrated molecular subtyping defines a curable oligometastatic state in colorectal liver metastasis. Nature communications 9, 1793 (2018).

20. Liberzon, A., et al. The Molecular Signatures Database (MSigDB) hallmark gene set collection. Cell Syst 1, 417–425 (2015).

21. Foroutan, M., et al. Single sample scoring of molecular phenotypes. BMC Bioinformatics 19, 404 (2018).

22. Fischer, M., Quaas, M., Steiner, L. & Engeland, K. The p53-p21-DREAM-CDE/CHR pathway regulates G2/M cell cycle genes. Nucleic Acids Res 44, 164–174 (2016).

23. Quaas, M., Muller, G.A. & Engeland, K. p53 can repress transcription of cell cycle genes through a p21(WAF1/CIP1)-dependent switch from MMB to DREAM protein complex binding at CHR promoter elements. Cell cycle (Georgetown, Tex.) 11, 4661–4672 (2012).

24. Engeland, K. Cell cycle arrest through indirect transcriptional repression by p53: I have a DREAM. Cell Death & Differentiation 25, 114–132 (2018).

25. Beggs, A.D., et al. Loss of expression of the double strand break repair protein ATM is associated with worse prognosis in colorectal cancer and loss of Ku70 expression is associated with CIN. Oncotarget 3, 1348–1355 (2012).

26. Bakkenist, C.J., Lee, J.J. & Schmitz, J.C. ATM Is Required for the Repair of Oxaliplatin-Induced DNA Damage in Colorectal Cancer. Clinical colorectal cancer 17, 255–257 (2018).

27. Sundar, R., et al. Ataxia Telangiectasia Mutated Protein Loss and Benefit From Oxaliplatin-based Chemotherapy in Colorectal Cancer. Clinical colorectal cancer 17, 280–284 (2018).

28. Maeda, O., et al. Alterations in gene expression and DNA methylation profiles in gastric cancer cells obtained from ascitic fluids collected before and after chemotherapy. Molecular and clinical oncology 11, 91–98 (2019).

29. Willis, S., et al. High Expression of FGD3, a Putative Regulator of Cell Morphology and Motility, Is Prognostic of Favorable Outcome in Multiple Cancers. JCO Precision Oncology, 1–13 (2017).

30. Wood, R.D., Mitchell, M., Sgouros, J. & Lindahl, T. Human DNA Repair Genes. Science 291, 1284–1289 (2001).

31. Hutz, J.E., et al. The multidimensional perturbation value: a single metric to measure similarity and activity of treatments in high-throughput multidimensional screens. Journal of biomolecular screening 18, 367–377 (2013).

32. Bliss, C.I. THE TOXICITY OF POISONS APPLIED JOINTLY1. Annals of Applied Biology 26, 585–615 (1939).

33. Thompson, R., Gatenby, R. & Sidi, S. How Cells Handle DNA Breaks during Mitosis: Detection, Signaling, Repair, and Fate Choice. Cells 8(2019).

34. Zhang, Y. & Hunter, T. Roles of Chk1 in cell biology and cancer therapy. International journal of cancer 134, 1013–1023 (2014).

35. Xie, Y., et al. Mps1/TTK: a novel target and biomarker for cancer. J Drug Target 25, 112–118 (2017).

36. Loewe, S. The problem of synergism and antagonism of combined drugs. Arzneimittelforschung 3, 285–290 (1953).

37. Marin, J.J.G., et al. Cellular Mechanisms Accounting for the Refractoriness of Colorectal Carcinoma to Pharmacological Treatment. Cancers (Basel) 12(2020).

38. Huang, M.-Y., et al. Predictive value of ERCC1, ERCC2, and XRCC1 overexpression for stage III colorectal cancer patients receiving FOLFOX-4 adjuvant chemotherapy. Journal of surgical oncology 108, 457–464 (2013).

39. Huang, M.-Y., et al. ERCC overexpression associated with a poor response of cT4b colorectal cancer with FOLFOX-based neoadjuvant concurrent chemoradiation. Oncology letters 20, 212–212 (2020).

40. Kondo, J., et al. High-throughput screening in colorectal cancer tissue-originated spheroids. Cancer science 110, 345–355 (2019).

41. Francies, H.E., Barthorpe, A., McLaren-Douglas, A., Barendt, W.J. & Garnett, M.J. Drug Sensitivity Assays of Human Cancer Organoid Cultures. Methods in molecular biology (Clifton, N.J.) 1576, 339–351 (2019).

42. Walsh, A.J., Cook, R.S. & Skala, M.C. Functional Optical Imaging of Primary Human Tumor Organoids: Development of a Personalized Drug Screen. Journal of nuclear medicine: official publication, Society of Nuclear Medicine 58, 1367–1372 (2017).

43. Boehnke, K., et al. Assay Establishment and Validation of a High-Throughput Screening Platform for Three-Dimensional Patient-Derived Colon Cancer Organoid Cultures. Journal of biomolecular screening 21, 931–941 (2016).

44. Kupchik, H.Z., Collins, E.A., O’Brien, M.J. & McCaffrey, R.P. Chemotherapy screening assay using 3-dimensional cell culture. Cancer letters 51, 11–16 (1990).

45. Driehuis, E., et al. Pancreatic cancer organoids recapitulate disease and allow personalized drug screening. Proc Natl Acad Sci U S A (2019).

46. van de Wetering, M., et al. Prospective Derivation of a Living Organoid Biobank of Colorectal Cancer Patients. Cell 161, 933–945 (2015).

47. Kondo, J., et al. High-throughput screening in colorectal cancer tissue-originated spheroids. Cancer science 110, 345–355 (2019).

48. Phan, N., et al. A simple high-throughput approach identifies actionable drug sensitivities in patient-derived tumor organoids. Communications Biology 2, 78 (2019).

49. Guinney, J., et al. The consensus molecular subtypes of colorectal cancer. Nature Medicine 21, 1350 (2015).

50. Singh, M.P., Rai, S., Pandey, A., Singh, N.K. & Srivastava, S. Molecular subtypes of colorectal cancer: An emerging therapeutic opportunity for personalized medicine. Genes & Diseases (2019).

51. Zhang, X., et al. Mps1 kinase regulates tumor cell viability via its novel role in mitochondria. Cell Death & Disease 7, e2292–e2292 (2016).

52. Fagan-Solis, K.D., et al. A P53-Independent DNA Damage Response Suppresses Oncogenic Proliferation and Genome Instability. Cell reports 30, 1385–1399.e1387 (2020).

53. Kim, E.M. & Burke, D.J. DNA damage activates the SAC in an ATM/ATR-dependent manner, independently of the kinetochore. PLoS genetics 4, e1000015–e1000015 (2008).

54. Ritchie, M.E., et al. limma powers differential expression analyses for RNA-sequencing and microarray studies. Nucleic Acids Res 43, e47 (2015).

55. Wu, D. & Smyth, G.K. Camera: a competitive gene set test accounting for inter-gene correlation. Nucleic Acids Res 40, e133–e133 (2012).

56. Markert, E.K., Mizuno, H., Vazquez, A. & Levine, A.J. Molecular classification of prostate cancer using curated expression signatures. Proc Natl Acad Sci U S A 108, 21276–21281 (2011).

57. Aran, D., Sirota, M. & Butte, A.J. Systematic pan-cancer analysis of tumour purity. Nature communications 6, 8971 (2015).

58. Cursons, J., et al. A Gene Signature Predicting Natural Killer Cell Infiltration and Improved Survival in Melanoma Patients. Cancer Immunol Res 7, 1162–1174 (2019).

59. Jerby-Arnon, L., et al. A Cancer Cell Program Promotes T Cell Exclusion and Resistance to Checkpoint Blockade. Cell 175, 984–997.e924 (2018).

60. Nirmal, A.J., et al. Immune Cell Gene Signatures for Profiling the Microenvironment of Solid Tumors. Cancer Immunol Res 6, 1388–1400 (2018).

