## Supplemental Figures 1-17 and Supp figure legends for "Targeting of TP53-independent cell cycle checkpoints overcomes FOLFOX resistance in Metastatic Colorectal Cancer"

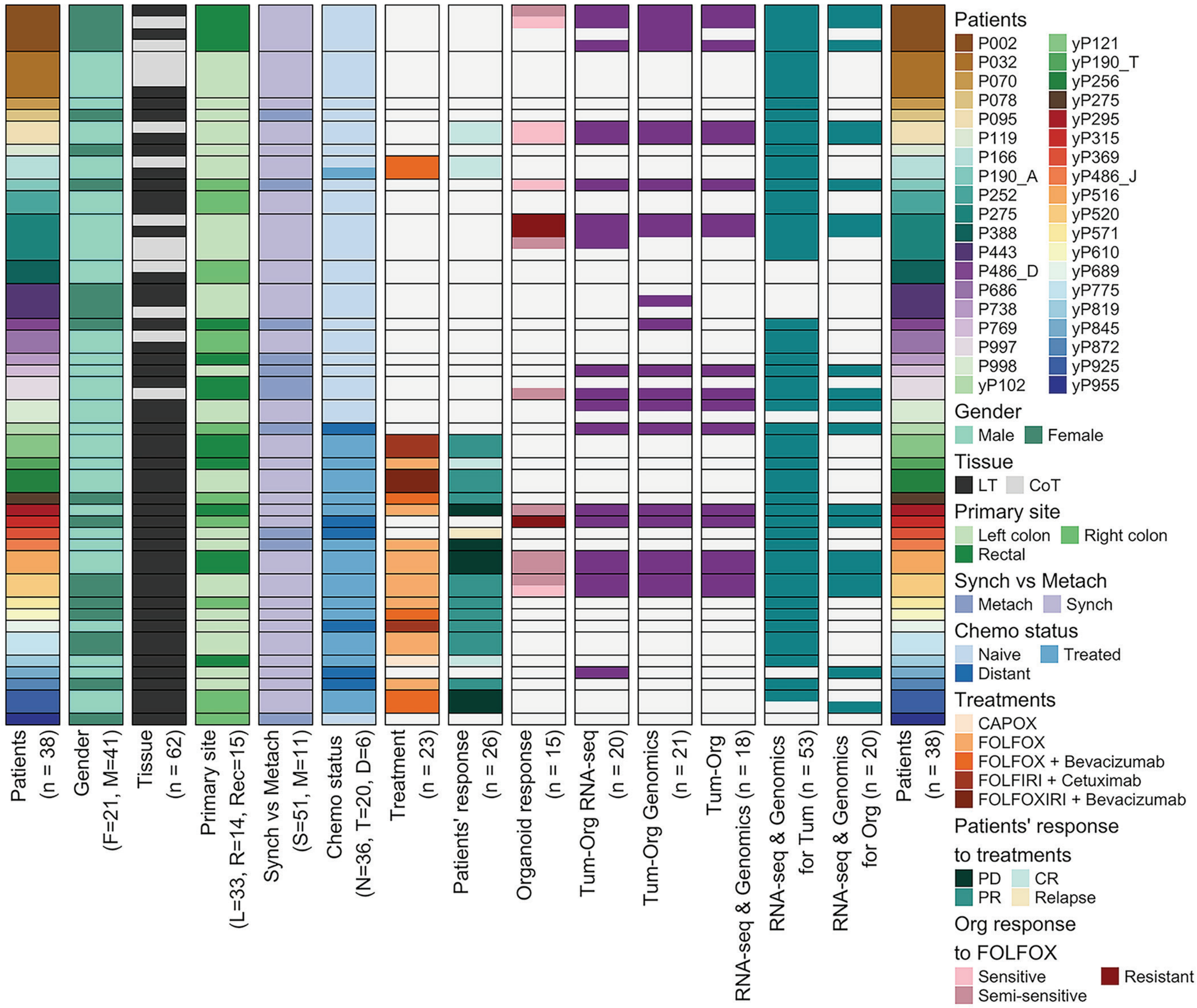

Patients (n = 38)
Gender (F=21, M=41)
Tissue (n = 62)
Primary site (L=33, R=14, Rec=15)
Synch vs Metach (S=51, M=11)
Chemo status (N=36, T=20, D=6)
Treatment (n = 23)
Patients' response (n = 26)
Organoid response (n = 15)
Tum-Org RNA-seq (n = 20)
Tum-Org Genomics (n = 21)
Tum-Org RNA-seq & Genomics (n = 18)
RNA-seq & Genomics for Tum (n = 53)
RNA-seq & Genomics for Org (n = 20)
Patients (n = 38)
Patients
P002 P032 P070 P078 P095 P119 P166 P190_A P252 P275 P388 P443 P486_D P686 P738 P769 P997 P998 yP102
yP121 yP190_T yP256 yP275 yP295 yP315 yP369 yP486_J yP516 yP520 yP571 yP610 yP689 yP775 yP819 yP845 yP872 yP925 yP955
Gender
Male Female
Tissue
LT CoT
Primary site
Left colon Right colon Rectal
Synch vs Metach
Metach Synch
Chemo status
Naive Treated Distant
Treatments
CAPOX
FOLFOX
FOLFOX + Bevacizumab
FOLFIRI + Cetuximab
FOLFOXIRI + Bevacizumab
Patients' response to treatments
PD CR PR Relapse
Org response to FOLFOX
Sensitive Resistant Semi-sensitive

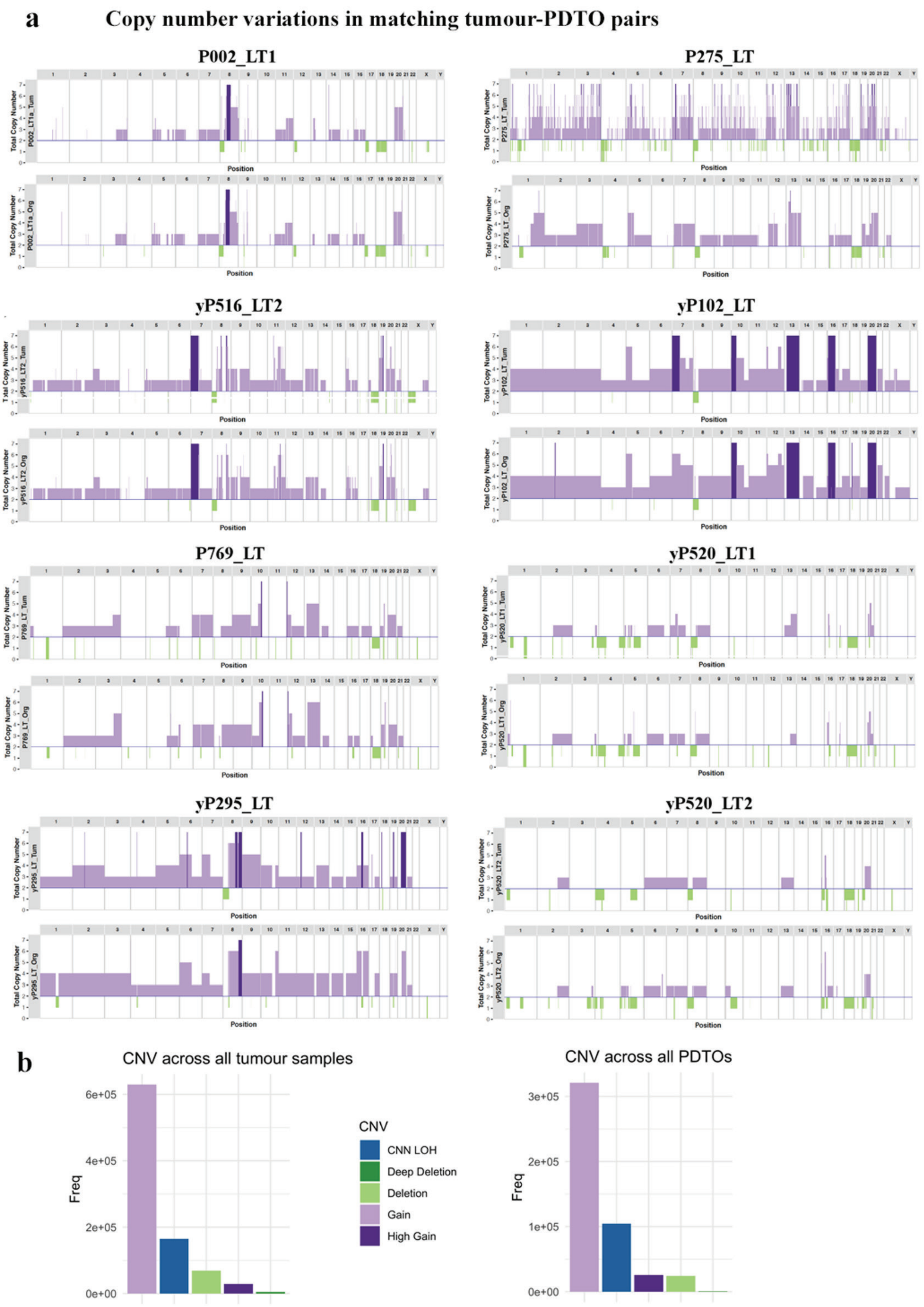
a
Copy number variations in matching tumour-PDTO pairs
P002_LT1
P275_LT
yP516_LT2
yP102_LT
P769_LT
yP520_LT1
yP295_LT
yP520_LT2
Total Copy Number
Position
b
CNV across all tumour samples
CNV across all PDTOs
Freq
6e+05
4e+05
2e+05
0e+00
3e+05
2e+05
1e+05
0e+00
CNV
CNN LOH
Deep Deletion
Deletion
Gain
High Gain

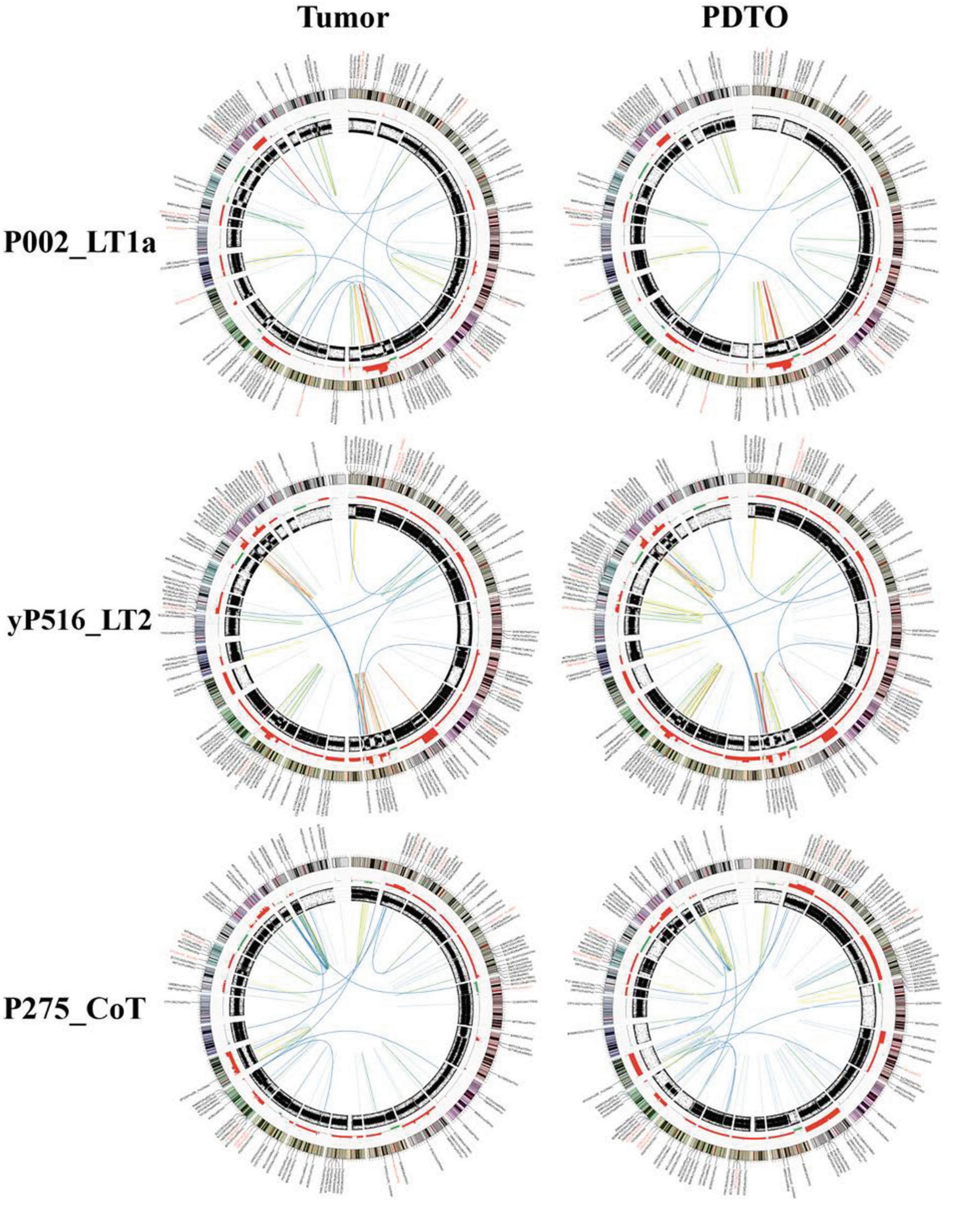
Tumor
PDTO
P002_LT1a
yP516_LT2
P275_CoT

a

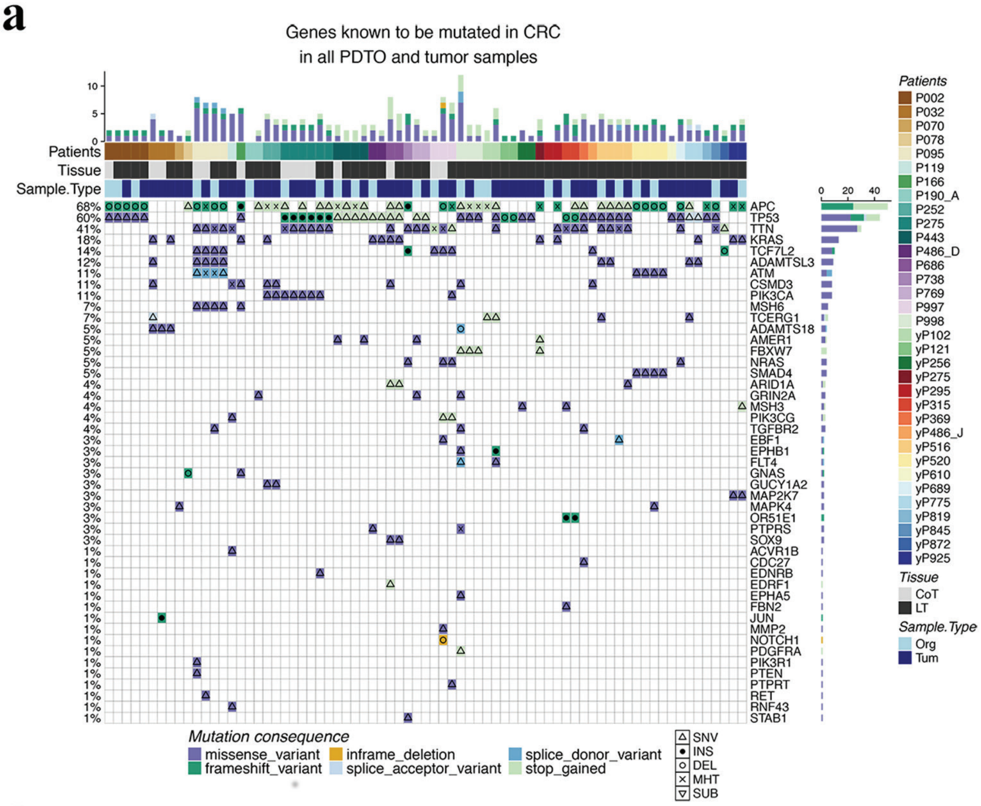
Genes known to be mutated in CRC
in all PDTO and tumor samples
Patients
Tissue
Sample.Type
68% APC
60% TP53
41% TTN
18% KRAS
14% TCF7L2
12% ADAMTSL3
11% ATM
11% CSMD3
11% PIK3CA
7% MSH6
7% TCERG1
5% ADAMTS18
5% AMER1
5% FBXW7
5% NRAS
5% SMAD4
4% ARID1A
4% GRIN2A
4% MSH3
4% PIK3CG
4% TGFBR2
3% EBF1
3% EPHB1
3% FLT4
3% GNAS
3% GUCY1A2
3% MAP2K7
3% MAPK4
3% OR51E1
3% PTPRS
3% SOX9
1% ACVR1B
1% CDC27
1% EDNRB
1% EDRF1
1% EPHA5
1% FBN2
1% JUN
1% MMP2
1% NOTCH1
1% PDGFRA
1% PIK3R1
1% PTEN
1% PTPRT
1% RET
1% RNF43
1% STAB1
Patients
P002 P032 P070 P078 P095 P119 P166 P190_A P252 P275 P443 P486_D P686 P738 P769 P997 P998 yP102 yP121 yP256 yP275 yP295 yP315 yP369 yP486_J yP516 yP520 yP610 yP689 yP775 yP819 yP845 yP872 yP925
Tissue
CoT
LT
Sample.Type
Org
Tum
Mutation consequence
missense_variant
frameshift_variant
inframe_deletion
splice_acceptor_variant
splice_donor_variant
stop_gained
SNV
INS
DEL
MHT
SUB

b

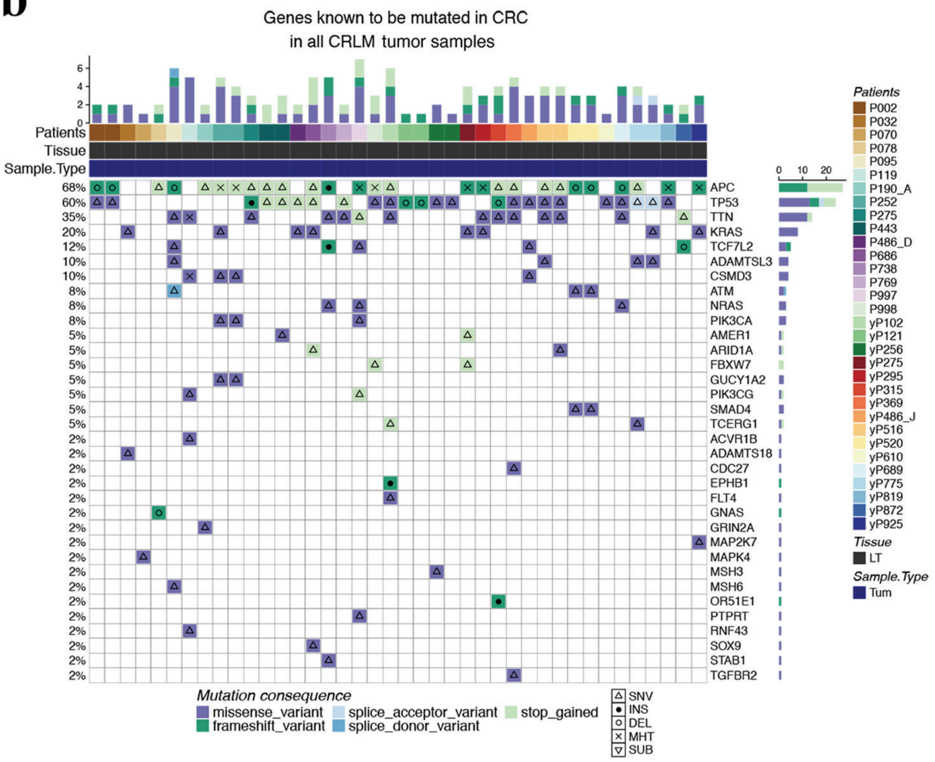
Genes known to be mutated in CRC
in all CRLM tumor samples
Patients
Tissue
Sample.Type
68% APC
60% TP53
35% TTN
20% KRAS
12% TCF7L2
10% ADAMTSL3
10% CSMD3
8% ATM
8% NRAS
8% PIK3CA
5% AMER1
5% ARID1A
5% FBXW7
5% GUCY1A2
5% PIK3CG
5% SMAD4
5% TCERG1
2% ACVR1B
2% ADAMTS18
2% CDC27
2% EPHB1
2% FLT4
2% GNAS
2% GRIN2A
2% MAP2K7
2% MAPK4
2% MSH3
2% MSH6
2% OR51E1
2% PTPRT
2% RNF43
2% SOX9
2% STAB1
2% TGFBR2
Patients
P002 P032 P070 P078 P095 P119 P190_A P252 P275 P443 P486_D P686 P738 P769 P997 P998 yP102 yP121 yP256 yP275 yP295 yP315 yP369 yP486_J yP516 yP520 yP610 yP689 yP775 yP819 yP872 yP925
Tissue
LT
Sample.Type
Tum
Mutation consequence
missense_variant
frameshift_variant
splice_acceptor_variant
splice_donor_variant
stop_gained
SNV
INS
DEL
MHT
SUB

a

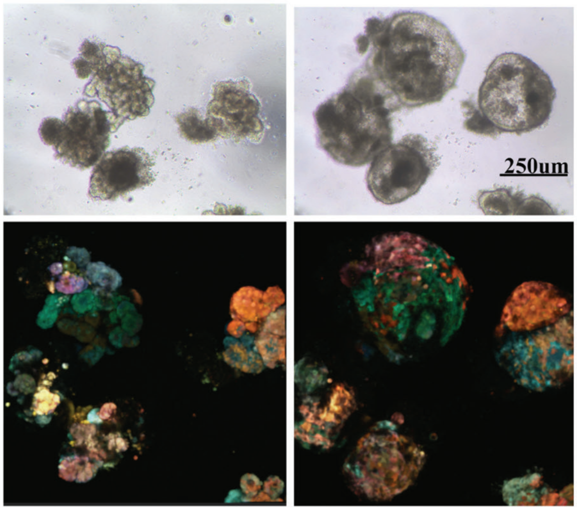
250um

b

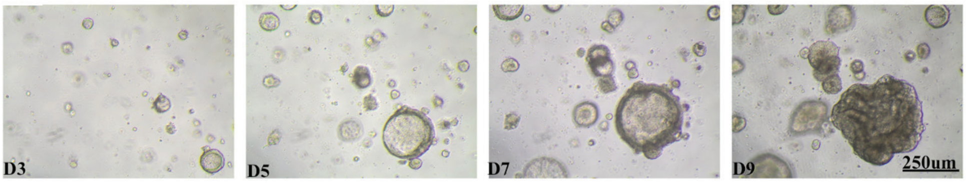
D3
D5
D7
D9
250um

c

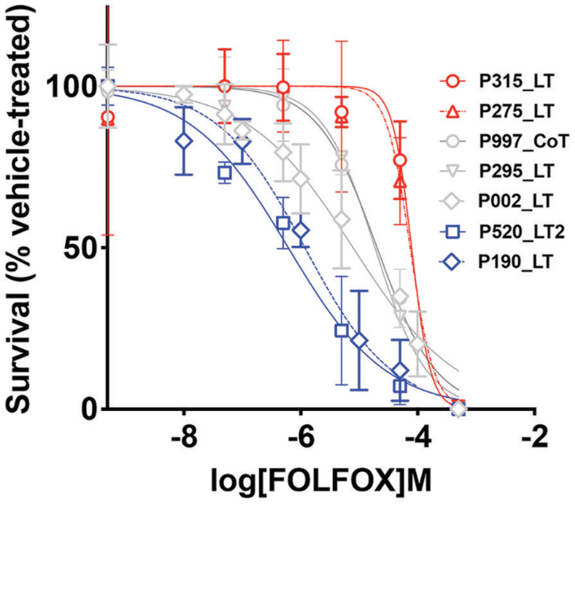
Survival (% vehicle-treated)
100
50
0
-8
-6
-4
-2
log[FOLFOX]M
P315_LT
P275_LT
P997_CoT
P295_LT
P002_LT
P520_LT2
P190_LT

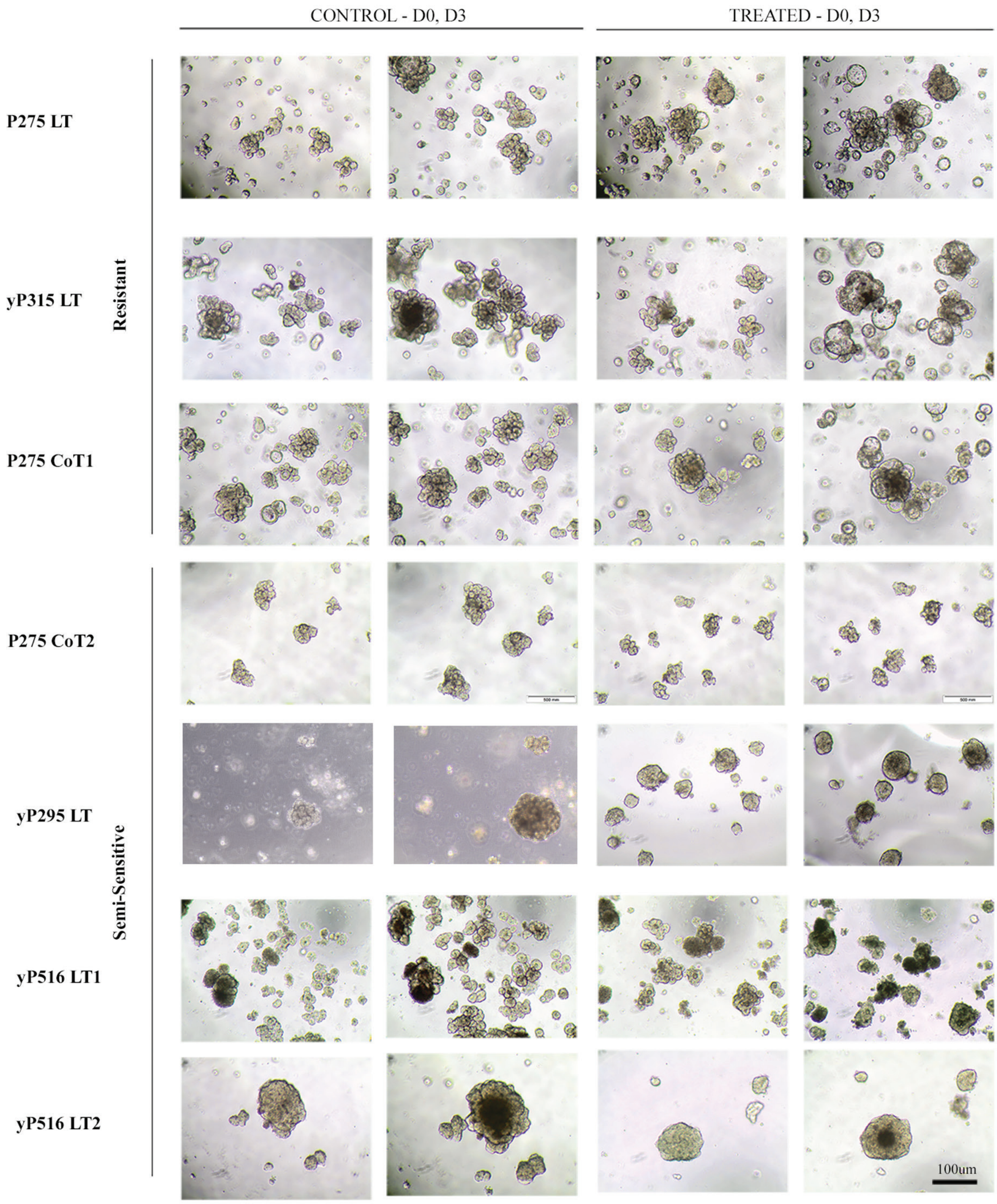

CONTROL - D0, D3
TREATED - D0, D3
P275 LT
yP315 LT
P275 CoT1
Resistant
P275 CoT2
yP295 LT
yP516 LT1
yP516 LT2
Semi-Sensitive
100um

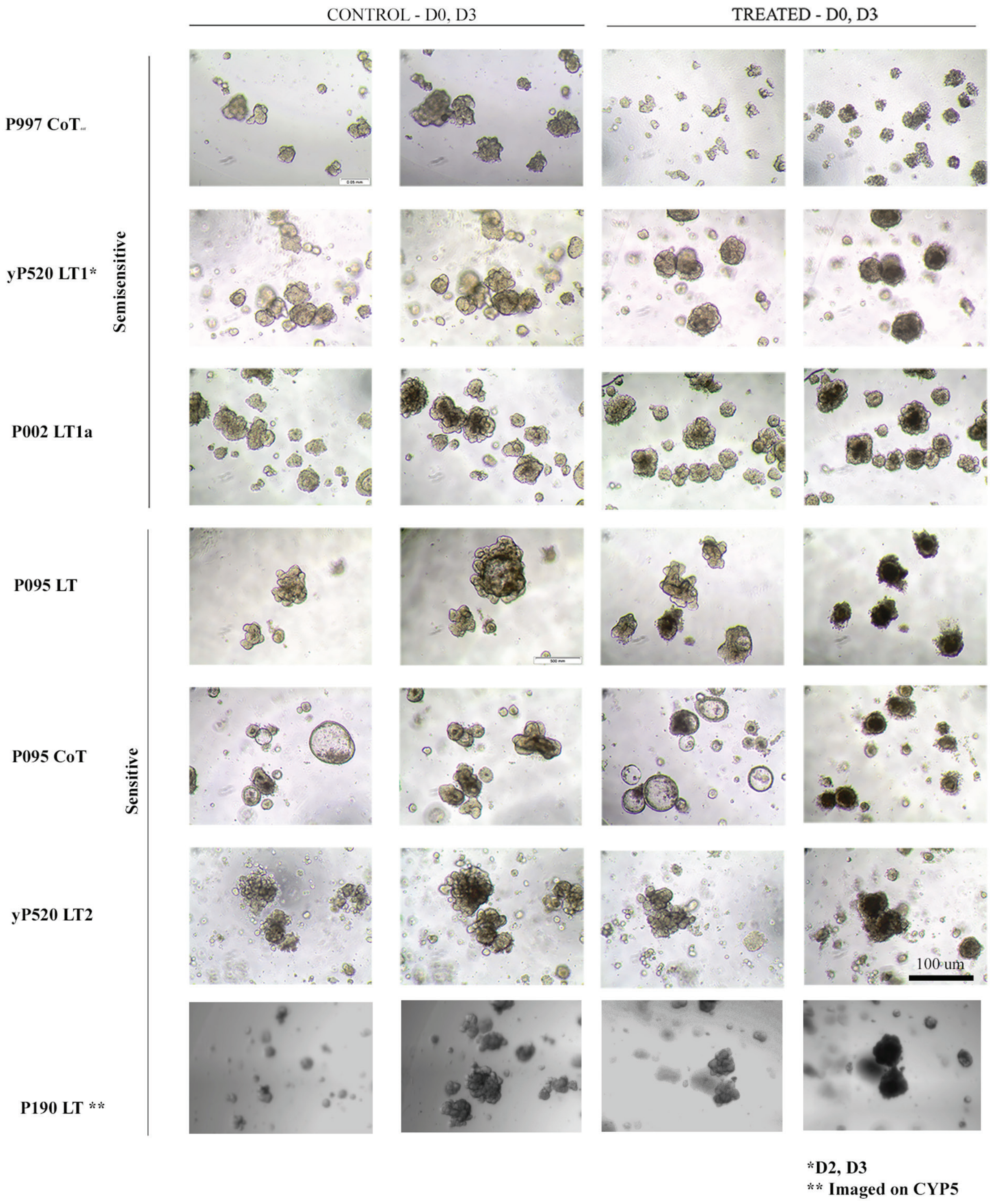
CONTROL - D0, D3
TREATED - D0, D3
Semisensitive
Sensitive
P997 CoT
yP520 LT1*
P002 LT1a
P095 LT
P095 CoT
yP520 LT2
P190 LT **
100 um
*D2, D3
** Imaged on CYP5

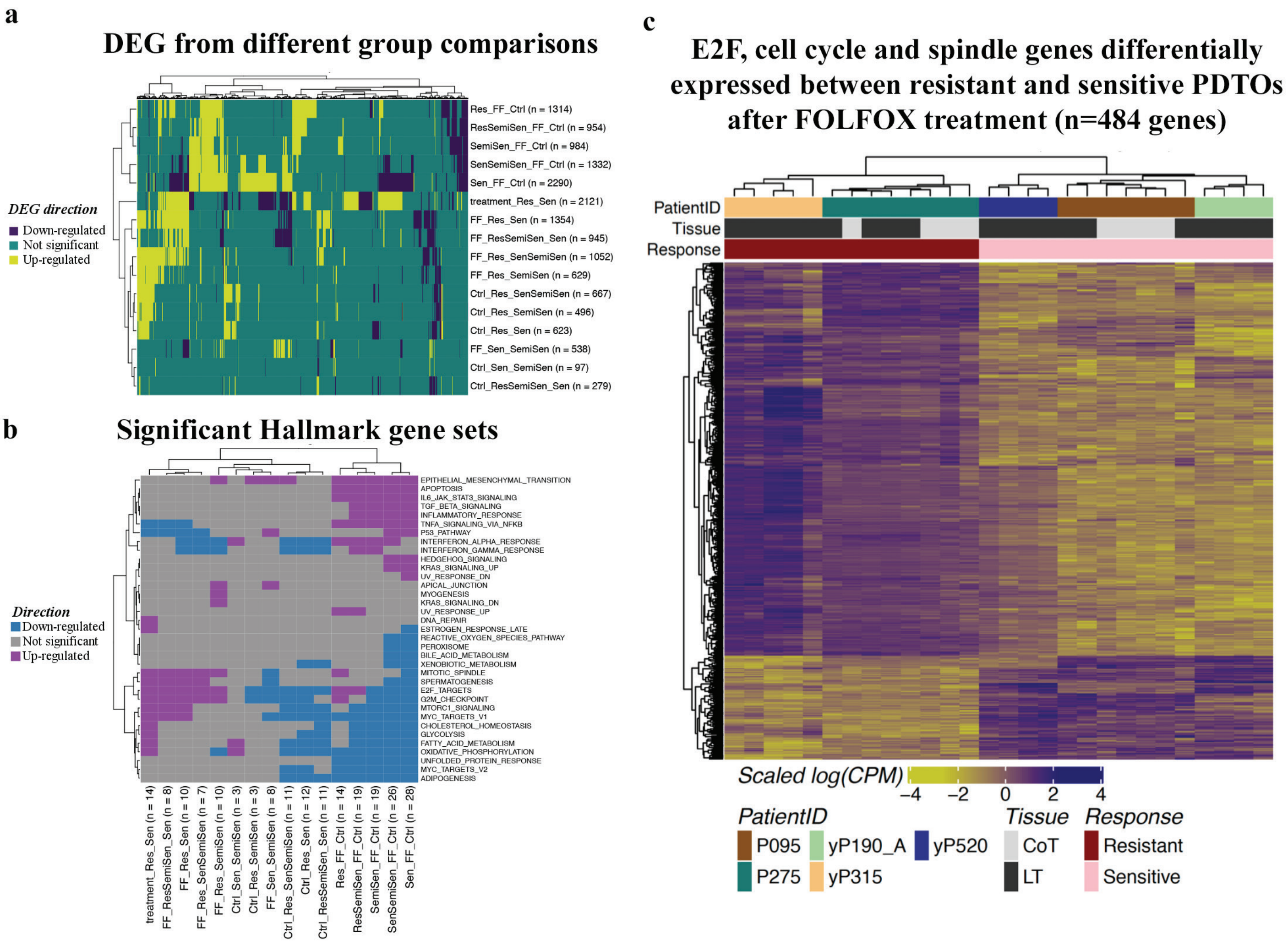
a
DEG from different group comparisons
DEG direction
Down-regulated
Not significant
Up-regulated
Res_FF_Ctrl (n = 1314)
ResSemiSen_FF_Ctrl (n = 954)
SemiSen_FF_Ctrl (n = 984)
SenSemiSen_FF_Ctrl (n = 1332)
Sen_FF_Ctrl (n = 2290)
treatment_Res_Sen (n = 2121)
FF_Res_Sen (n = 1354)
FF_ResSemiSen_Sen (n = 945)
FF_Res_SenSemiSen (n = 1052)
FF_Res_SemiSen (n = 629)
Ctrl_Res_SenSemiSen (n = 667)
Ctrl_Res_SemiSen (n = 496)
Ctrl_Res_Sen (n = 623)
FF_Sen_SemiSen (n = 538)
Ctrl_Sen_SemiSen (n = 97)
Ctrl_ResSemiSen_Sen (n = 279)
b
Significant Hallmark gene sets
Direction
Down-regulated
Not significant
Up-regulated
EPITHELIAL_MESENCHYMAL_TRANSITION
APOPTOSIS
IL6_JAK_STAT3_SIGNALING
TGF_BETA_SIGNALING
INFLAMMATORY_RESPONSE
TNFA_SIGNALING_VIA_NFKB
P53_PATHWAY
INTERFERON_ALPHA_RESPONSE
INTERFERON_GAMMA_RESPONSE
HEDGEHOG_SIGNALING
KRAS_SIGNALING_UP
UV_RESPONSE_DN
APICAL_JUNCTION
MYOGENESIS
KRAS_SIGNALING_DN
UV_RESPONSE_UP
DNA_REPAIR
ESTROGEN_RESPONSE_LATE
REACTIVE_OXYGEN_SPECIES_PATHWAY
PEROXISOME
BILE_ACID_METABOLISM
XENOBIOTIC_METABOLISM
MITOTIC_SPINDLE
SPERMATOGENESIS
E2F_TARGETS
G2M_CHECKPOINT
MTORC1_SIGNALING
MYC_TARGETS_V1
CHOLESTEROL_HOMEOSTASIS
GLYCOLYSIS
FATTY_ACID_METABOLISM
OXIDATIVE_PHOSPHORYLATION
UNFOLDED_PROTEIN_RESPONSE
MYC_TARGETS_V2
ADIPOGENESIS
treatment_Res_Sen (n = 14)
FF_ResSemiSen_Sen (n = 8)
FF_Res_Sen (n = 10)
FF_Res_SenSemiSen (n = 7)
FF_Res_SemiSen (n = 10)
Ctrl_Sen_SemiSen (n = 3)
Ctrl_Res_SemiSen (n = 3)
FF_Sen_SemiSen (n = 8)
Ctrl_Res_SenSemiSen (n = 11)
Ctrl_Res_Sen (n = 12)
Ctrl_ResSemiSen_Sen (n = 11)
Res_FF_Ctrl (n = 14)
ResSemiSen_FF_Ctrl (n = 19)
SemiSen_FF_Ctrl (n = 19)
SenSemiSen_FF_Ctrl (n = 26)
Sen_FF_Ctrl (n = 28)
c
E2F, cell cycle and spindle genes differentially expressed between resistant and sensitive PDTOs after FOLFOX treatment (n=484 genes)
PatientID
Tissue
Response
Scaled log(CPM)
−4
−2
0
2
4
PatientID
P095
yP190_A
yP520
P275
yP315
Tissue
CoT
LT
Response
Resistant
Sensitive

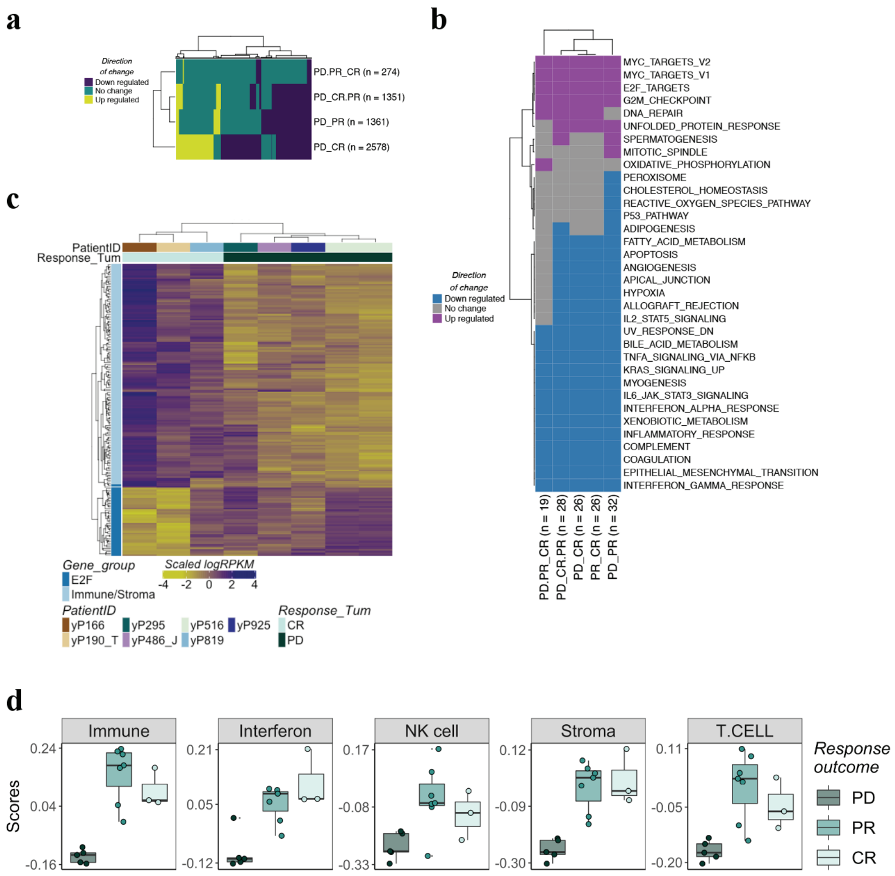

a
Direction of change
Down regulated
No change
Up regulated
PD.PR_CR (n = 274)
PD_CR.PR (n = 1351)
PD_PR (n = 1361)
PD_CR (n = 2578)
b
MYC_TARGETS_V2
MYC_TARGETS_V1
E2F_TARGETS
G2M_CHECKPOINT
DNA_REPAIR
UNFOLDED_PROTEIN_RESPONSE
SPERMATOGENESIS
MITOTIC_SPINDLE
OXIDATIVE_PHOSPHORYLATION
PEROXISOME
CHOLESTEROL_HOMEOSTASIS
REACTIVE_OXYGEN_SPECIES_PATHWAY
P53_PATHWAY
ADIPOGENESIS
FATTY_ACID_METABOLISM
APOPTOSIS
ANGIOGENESIS
APICAL_JUNCTION
HYPOXIA
ALLOGRAFT_REJECTION
IL2_STAT5_SIGNALING
UV_RESPONSE_DN
BILE_ACID_METABOLISM
TNFA_SIGNALING_VIA_NFKB
KRAS_SIGNALING_UP
MYOGENESIS
IL6_JAK_STAT3_SIGNALING
INTERFERON_ALPHA_RESPONSE
XENOBIOTIC_METABOLISM
INFLAMMATORY_RESPONSE
COMPLEMENT
COAGULATION
EPITHELIAL_MESENCHYMAL_TRANSITION
INTERFERON_GAMMA_RESPONSE
Direction of change
Down regulated
No change
Up regulated
PD.PR_CR (n = 19)
PD_CR.PR (n = 28)
PD_CR (n = 26)
PR_CR (n = 26)
PD_PR (n = 32)
c
PatientID
Response_Tum
Gene_group
E2F
Immune/Stroma
Scaled logRPKM
-4 -2 0 2 4
PatientID
yP166
yP190_T
yP295
yP486_J
yP516
yP819
yP925
Response_Tum
CR
PD
d
Immune
Interferon
NK cell
Stroma
T.CELL
Scores
0.24
0.04
-0.16
0.21
0.05
-0.12
0.17
-0.08
-0.33
0.12
-0.09
-0.30
0.11
-0.05
-0.20
Response outcome
PD
PR
CR

a

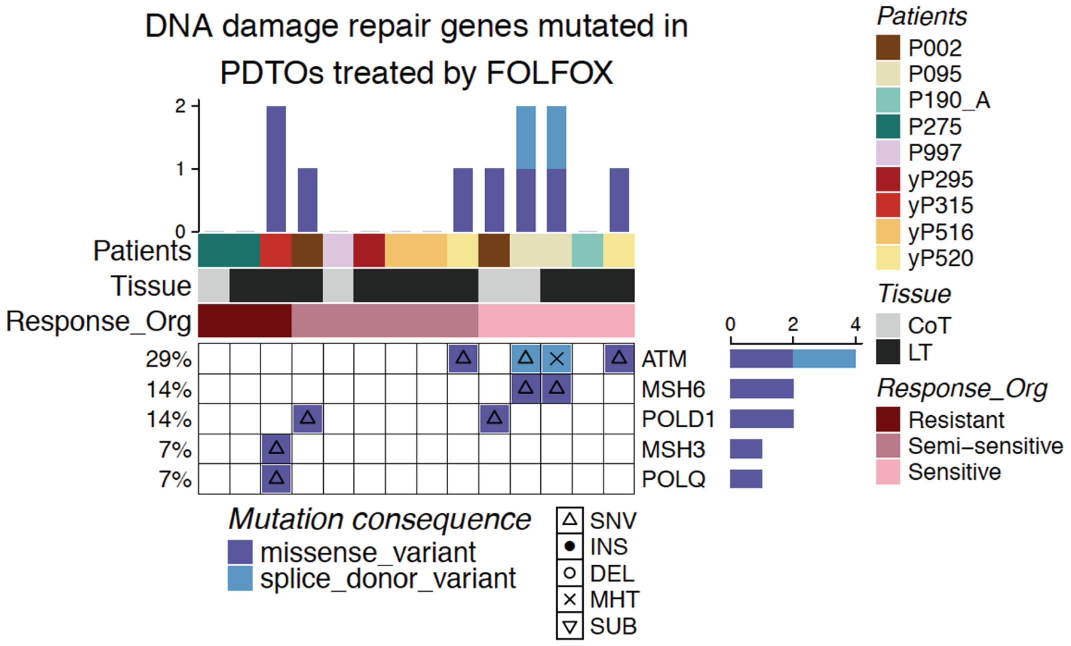
DNA damage repair genes mutated in
PDTOs treated by FOLFOX
Patients
Tissue
Response_Org
29%
14%
14%
7%
7%
ATM
MSH6
POLD1
MSH3
POLQ
Mutation consequence
missense_variant
splice_donor_variant
SNV
INS
DEL
MHT
SUB
Patients
P002
P095
P190_A
P275
P997
yP295
yP315
yP516
yP520
Tissue
CoT
LT
Response_Org
Resistant
Semi–sensitive
Sensitive

b

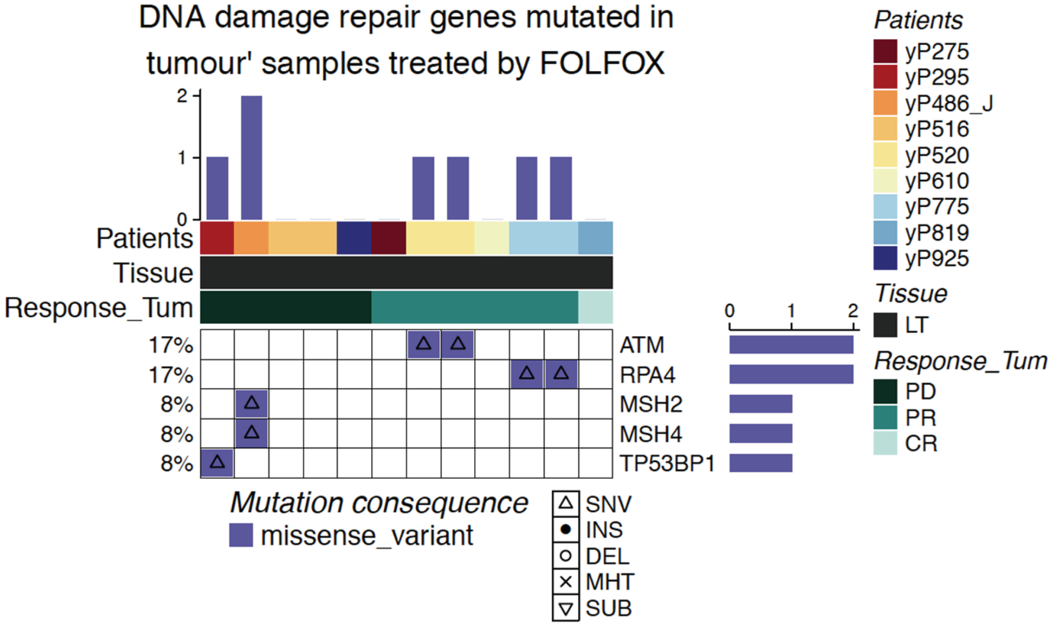
DNA damage repair genes mutated in
tumour' samples treated by FOLFOX
Patients
Tissue
Response_Tum
17%
17%
8%
8%
8%
ATM
RPA4
MSH2
MSH4
TP53BP1
Mutation consequence
missense_variant
SNV
INS
DEL
MHT
SUB
Patients
yP275
yP295
yP486_J
yP516
yP520
yP610
yP775
yP819
yP925
Tissue
LT
Response_Tum
PD
PR
CR

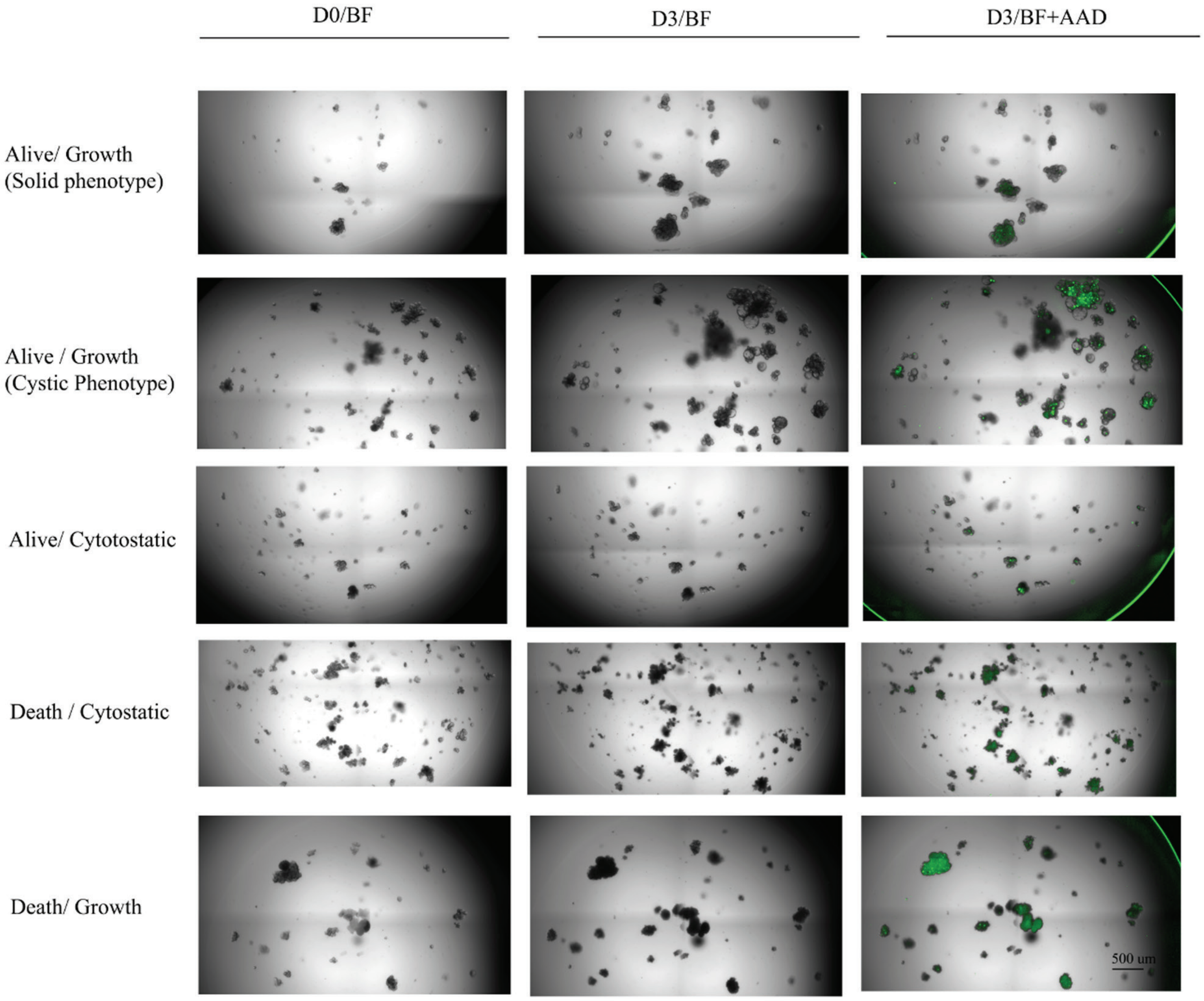

D0/BF
D3/BF
D3/BF+AAD
Alive/ Growth
(Solid phenotype)
Alive / Growth
(Cystic Phenotype)
Alive/ Cytotostatic
Death / Cytostatic
Death/ Growth
500 um

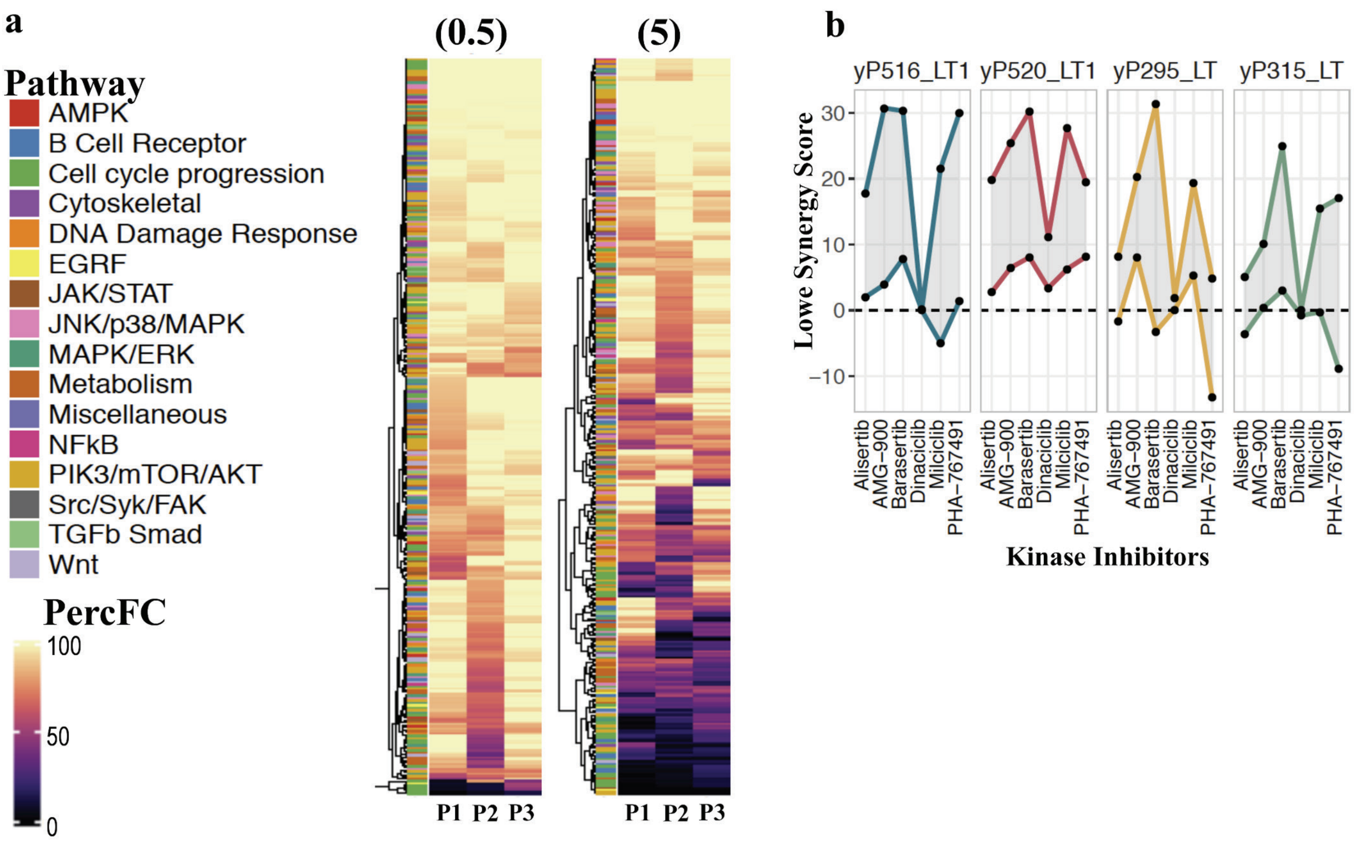
a
Pathway
AMPK
B Cell Receptor
Cell cycle progression
Cytoskeletal
DNA Damage Response
EGRF
JAK/STAT
JNK/p38/MAPK
MAPK/ERK
Metabolism
Miscellaneous
NFkB
PIK3/mTOR/AKT
Src/Syk/FAK
TGFb Smad
Wnt
PercFC
100
50
0
(0.5)
(5)
P1 P2 P3
P1 P2 P3
b
yP516_LT1
yP520_LT1
yP295_LT
yP315_LT
Lowe Synergy Score
30
20
10
0
−10
Alisertib
AMG−900
Barasertib
Dinaciclib
Milciclib
PHA−767491
Kinase Inhibitors

CHK1 Inhibitors and Controls - P1 (yP516_LT1)

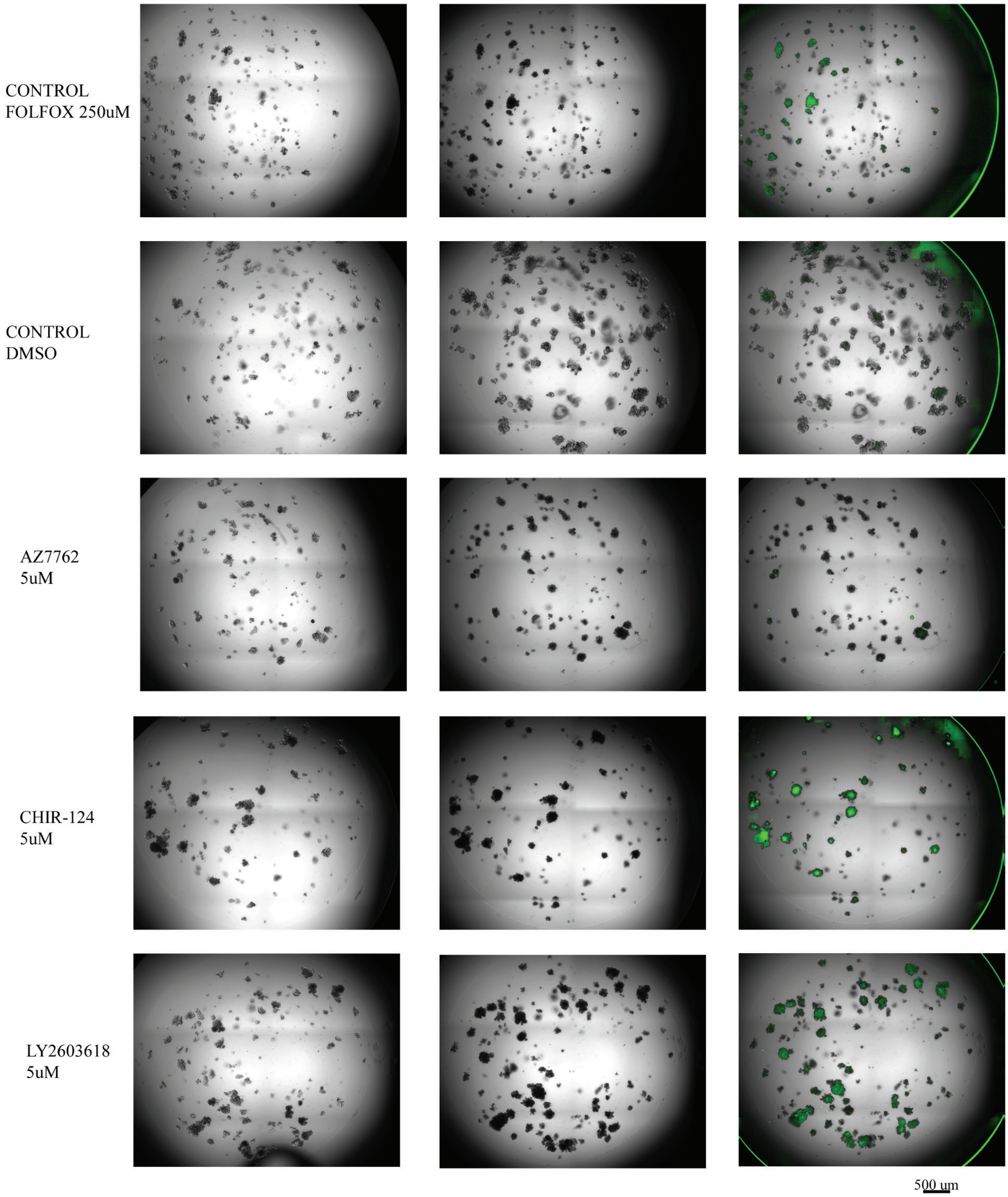

CHK1 Inhibitors and Controls - P2 (yP520_LT1)

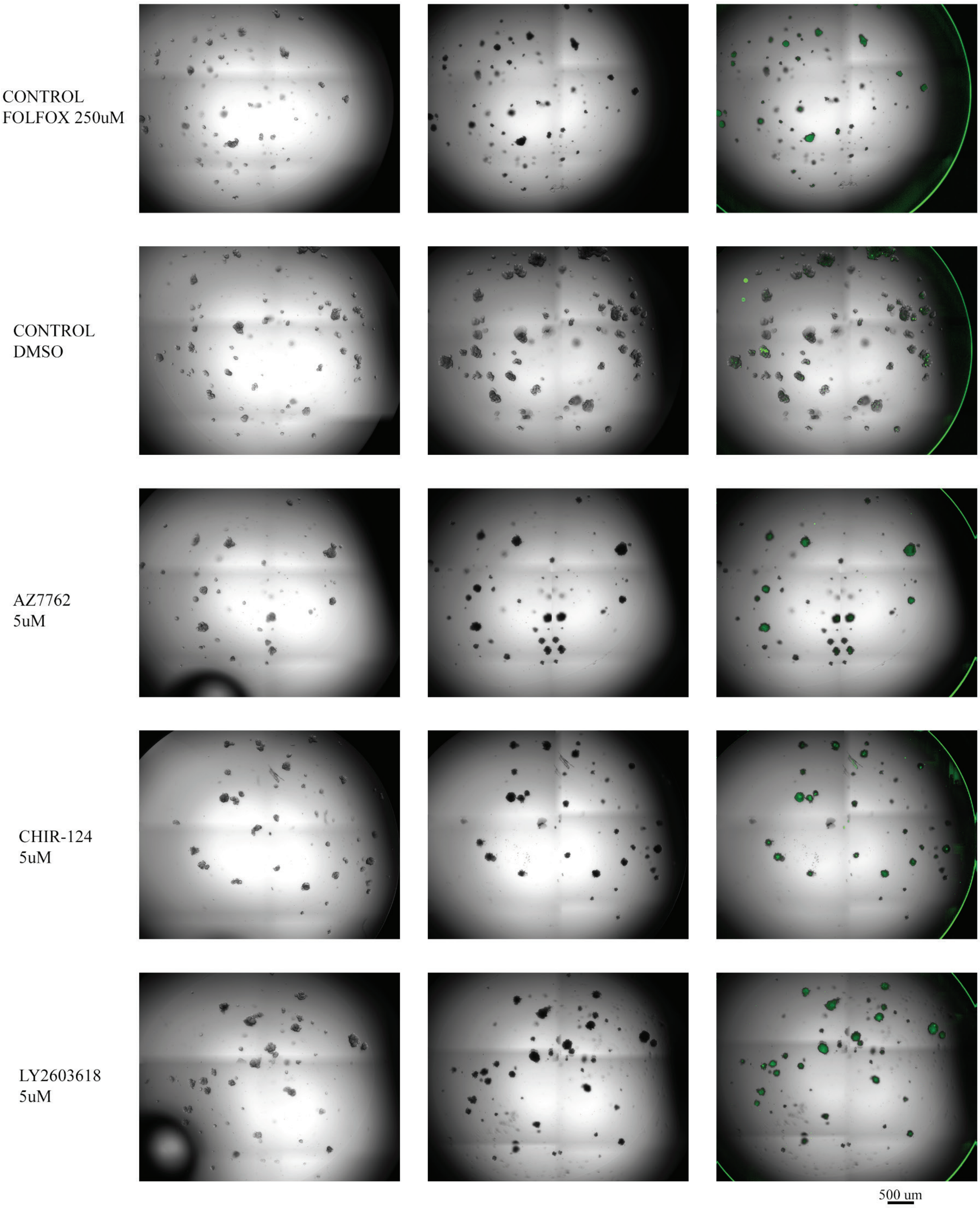

### CHK1 Inhibitors and Controls - P3 (yP295_LT)

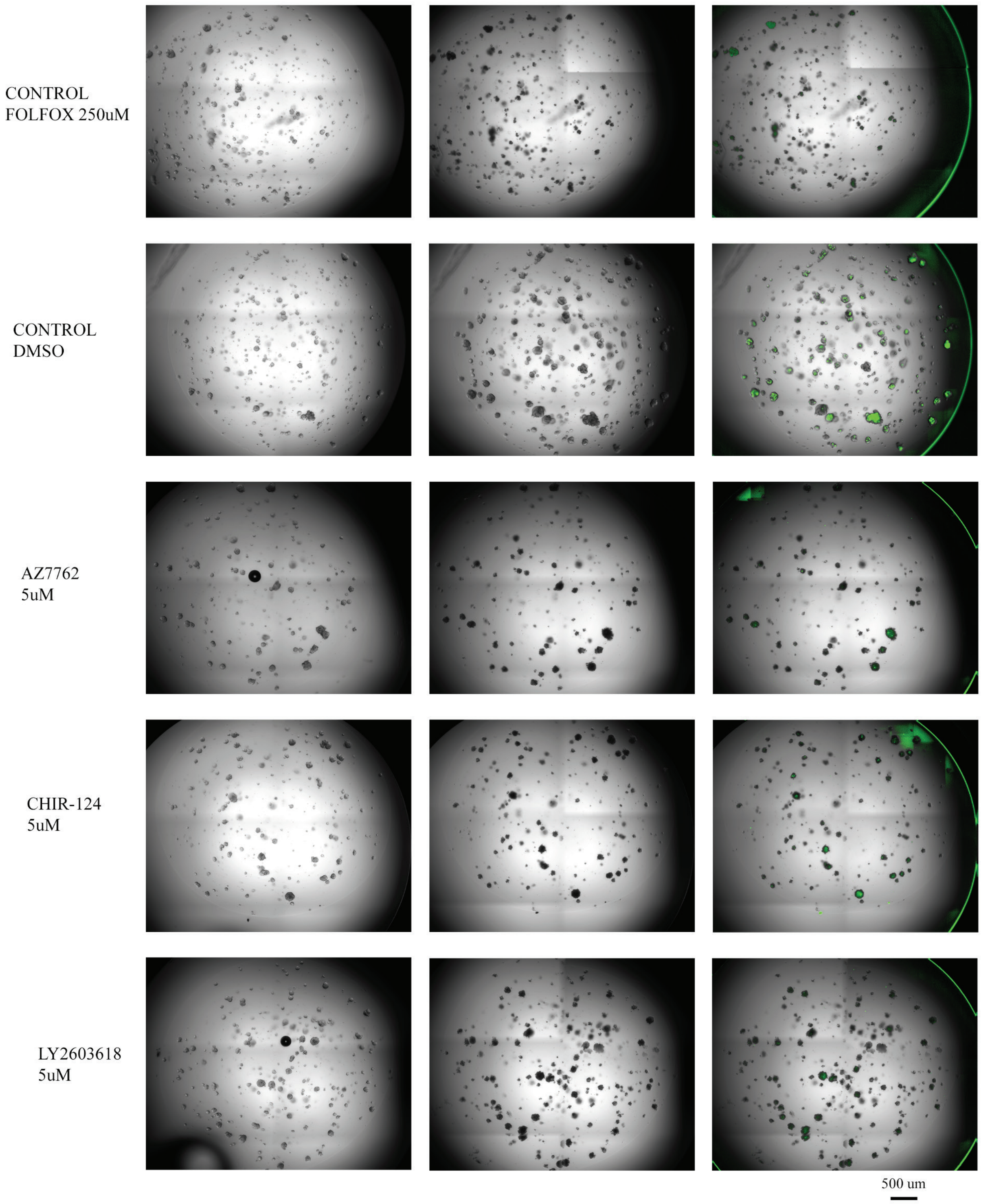

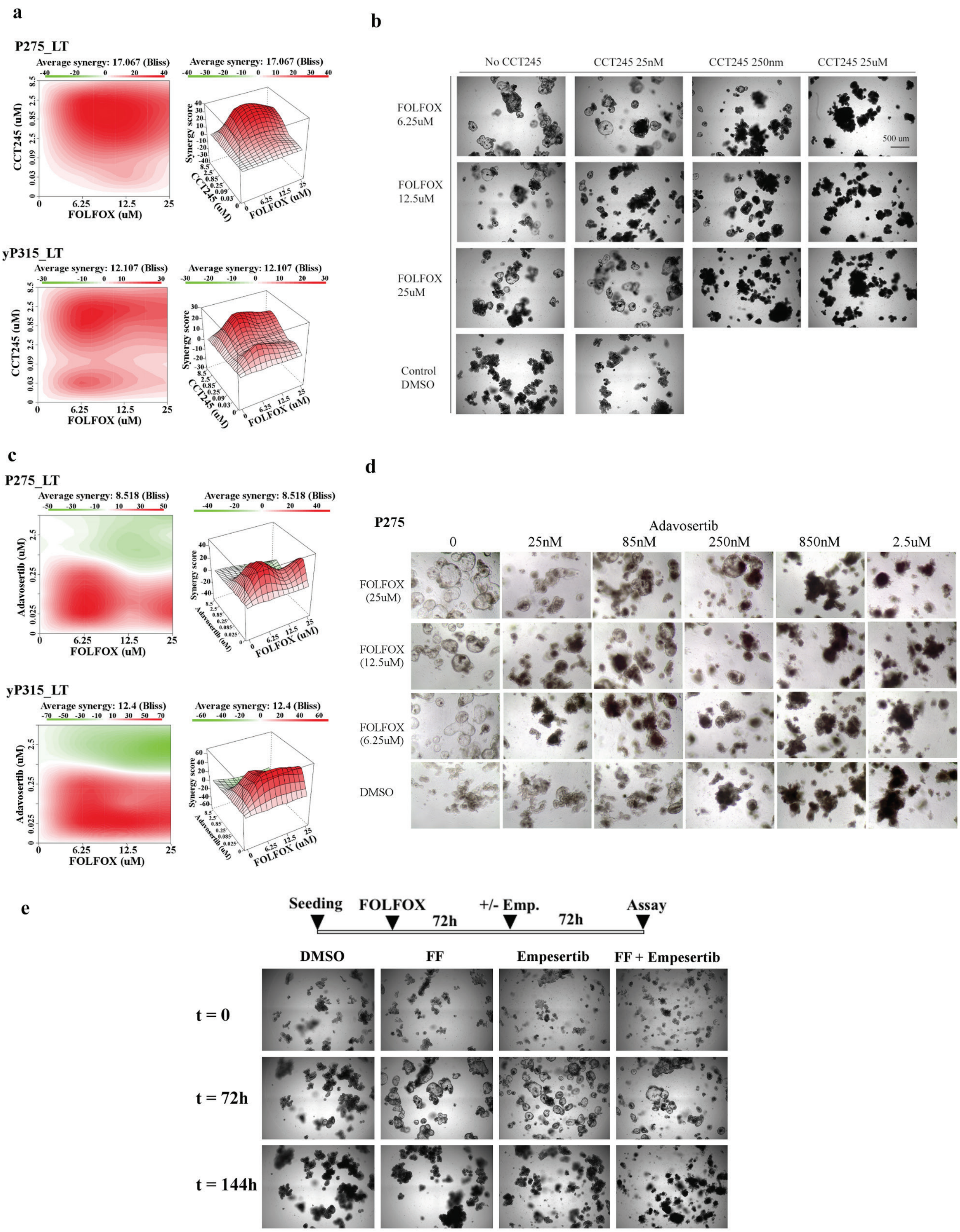

a
P275_LT
Average synergy: 17.067 (Bliss)
CCT245 (uM)
FOLFOX (uM)
Synergy score
yP315_LT
Average synergy: 12.107 (Bliss)
b
No CCT245
CCT245 25nM
CCT245 250nm
CCT245 25uM
FOLFOX 6.25uM
FOLFOX 12.5uM
FOLFOX 25uM
Control DMSO
500 um
c
P275_LT
Average synergy: 8.518 (Bliss)
Adavosertib (uM)
yP315_LT
Average synergy: 12.4 (Bliss)
d
P275
Adavosertib
0
25nM
85nM
250nM
850nM
2.5uM
FOLFOX (25uM)
FOLFOX (12.5uM)
FOLFOX (6.25uM)
DMSO
e
Seeding
FOLFOX
72h
+/- Emp.
72h
Assay
DMSO
FF
Empesertib
FF + Empesertib
t = 0
t = 72h
t = 144h

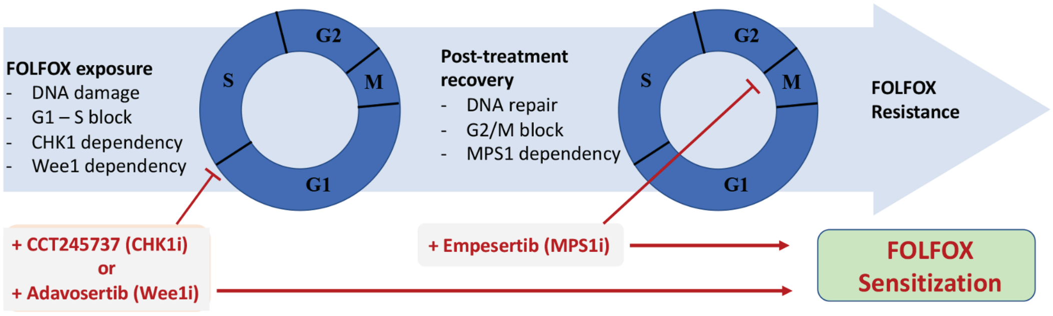

FOLFOX exposure
- DNA damage
- G1 – S block
- CHK1 dependency
- Wee1 dependency
S
G2
M
G1
Post-treatment recovery
- DNA repair
- G2/M block
- MPS1 dependency
S
G2
M
G1
FOLFOX Resistance
+ CCT245737 (CHK1i)
or
+ Adavosertib (Wee1i)
+ Empesertib (MPS1i)
FOLFOX Sensitization

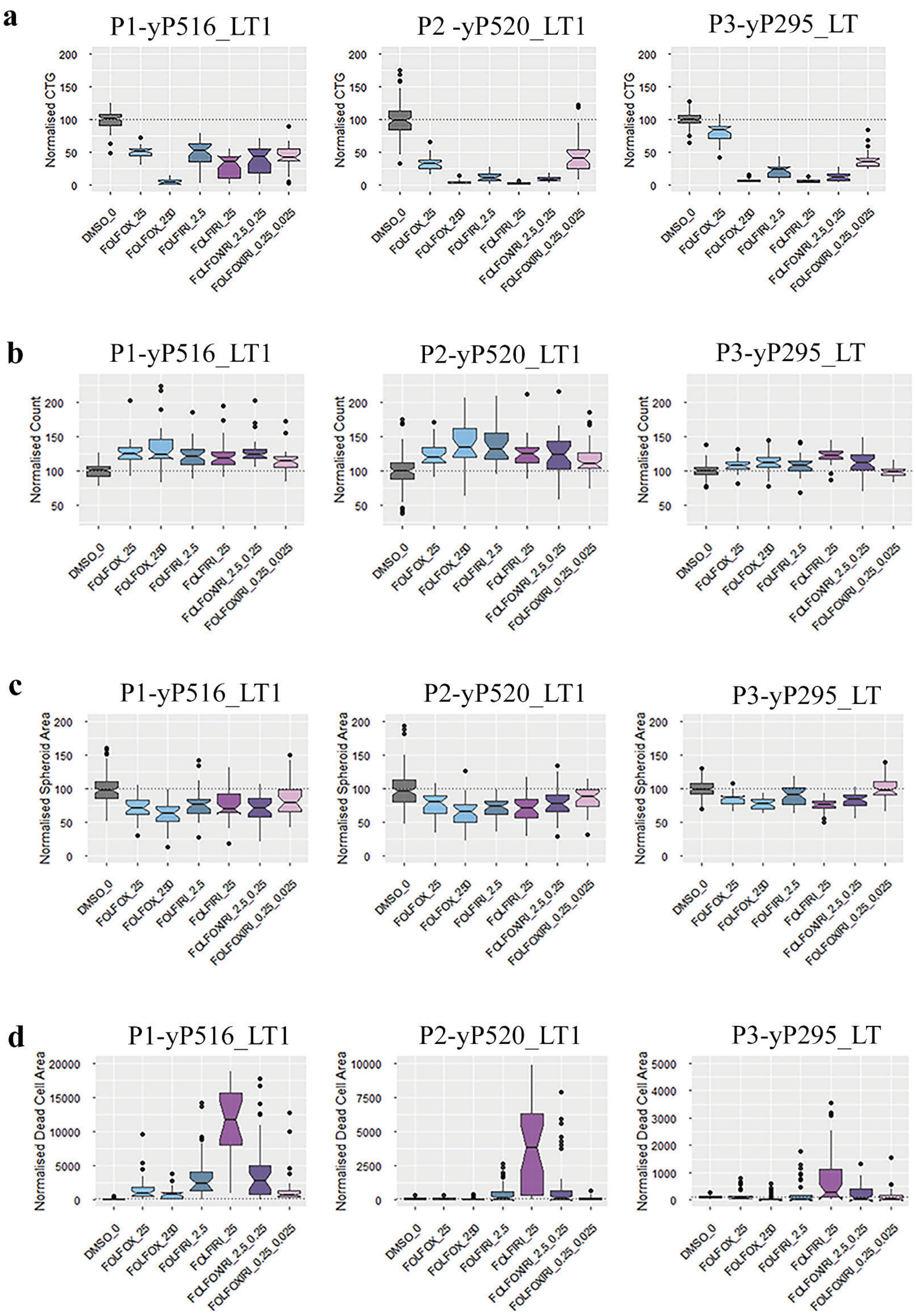

a
P1-yP516_LT1
P2 -yP520_LT1
P3-yP295_LT
Normalised CTG
b
P1-yP516_LT1
P2-yP520_LT1
P3-yP295_LT
Normalised Count
c
P1-yP516_LT1
P2-yP520_LT1
P3-yP295_LT
Normalised Spheroid Area
d
P1-yP516_LT1
P2-yP520_LT1
P3-yP295_LT
Normalised Dead Cell Area
DMSO_0
FOLFOX_25
FOLFOX_250
FOLFIRI_2.5
FOLFIRI_25
FOLFOXIRI_2.5_0.25
FOLFOXIRI_0.25_0.025

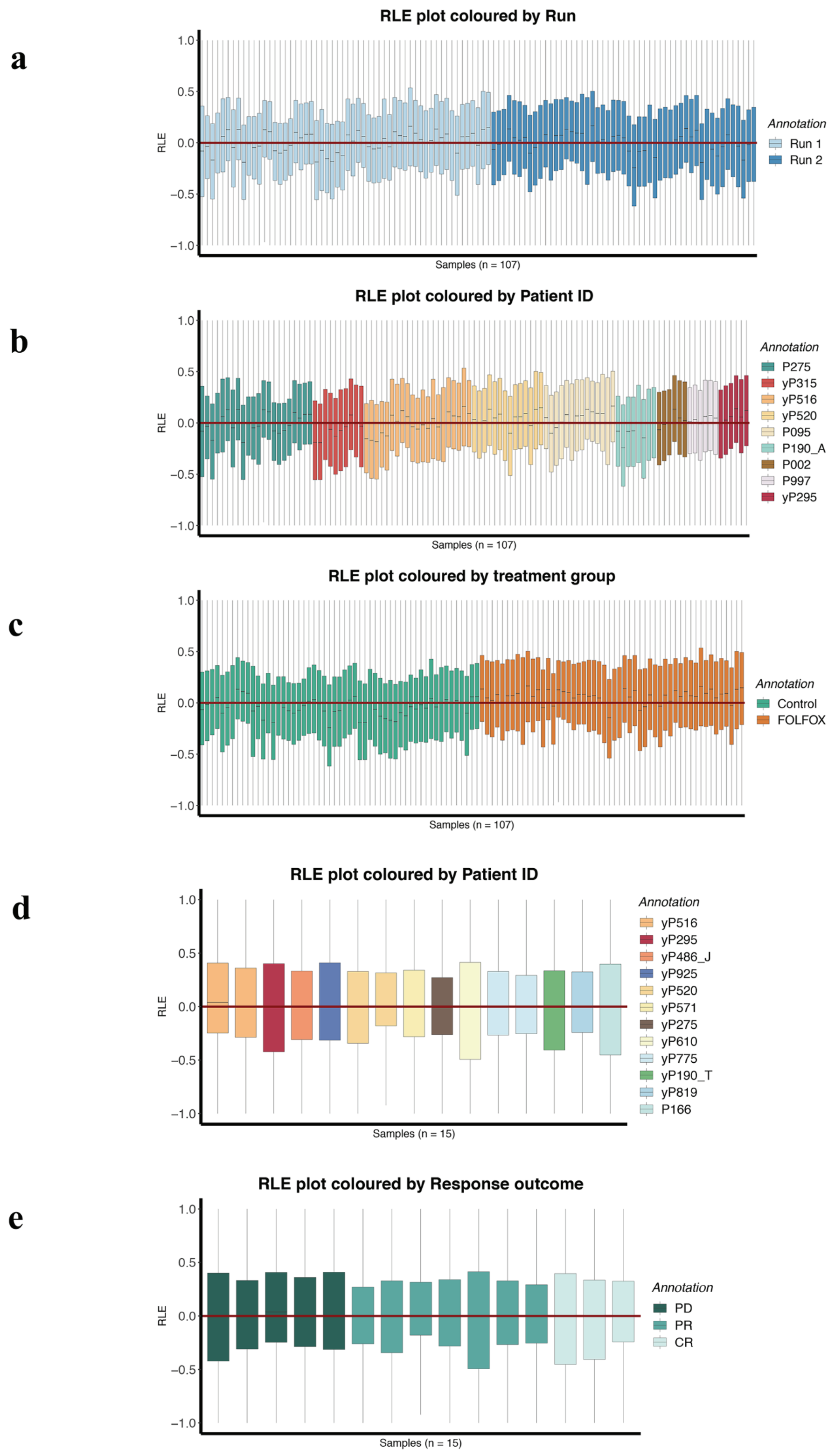
a
RLE plot coloured by Run
RLE
Samples (n = 107)
Annotation
Run 1
Run 2
b
RLE plot coloured by Patient ID
RLE
Samples (n = 107)
Annotation
P275
yP315
yP516
yP520
P095
P190_A
P002
P997
yP295
c
RLE plot coloured by treatment group
RLE
Samples (n = 107)
Annotation
Control
FOLFOX
d
RLE plot coloured by Patient ID
RLE
Samples (n = 15)
Annotation
yP516
yP295
yP486_J
yP925
yP520
yP571
yP275
yP610
yP775
yP190_T
yP819
P166
e
RLE plot coloured by Response outcome
RLE
Samples (n = 15)
Annotation
PD
PR
CR

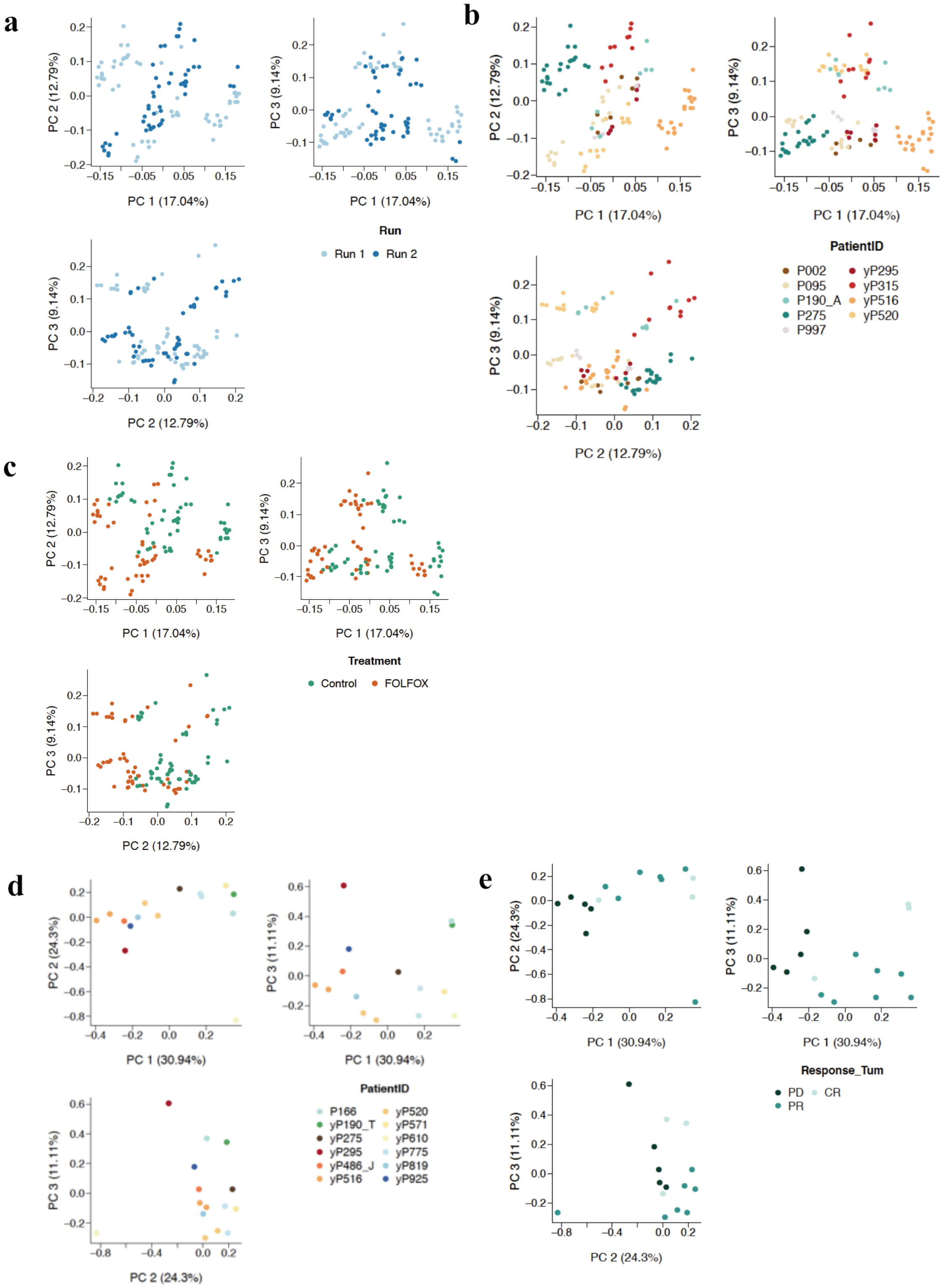

a
PC 1 (17.04%)
PC 2 (12.79%)
PC 3 (9.14%)
Run
Run 1
Run 2
b
PatientID
P002
P095
P190_A
P275
P997
yP295
yP315
yP516
yP520
c
Treatment
Control
FOLFOX
d
PC 1 (30.94%)
PC 2 (24.3%)
PC 3 (11.11%)
PatientID
P166
yP190_T
yP275
yP295
yP486_J
yP516
yP520
yP571
yP610
yP775
yP819
yP925
e
Response_Tum
PD
PR
CR

**Supplementary Figures.**

**Supplementary Figure 1. Summary of Available Data.** Graphic overview of samples collected and analyzed: A total of 38 patients were recruited, with a total of 62 tumor samples collected (49 CRLM, 13 colorectal primary tumors). PDTOs were grown from 20 of these samples. For mutation analysis using the WGS/WES data, we had a total of 89 samples (including both tumor and organoids), and after removal of low-quality samples (based on the genomics data, e.g. low estimated purity), this decreased to 78 samples, including a small number of samples from which we obtained both WGS and WES data. Mutational information from these replicated samples was then consolidated (see methods), resulting in 73 samples used for mutation analysis (Suppl Table 7). For RNA-seq data analysis, we removed the RNA-seq data of the primary colon tumor samples from patient PC002 (PC002_CoT) due to its low quality, resulted in 18 matched tumor-PDTO samples. We also removed these samples when performing DE analyses in the PDTOs treated with FOLFOX.

**Supplementary Figure 2. Copy Number Variation concordance between PDTOs and parent Tumors. a)** Piano plots demonstrating concordance of copy number variation patterns between parent tumor (*upper plot in each pair*) and matching PDTOs (*lower plot in each pair*) **b)** Bar chart highlighting the concordance in the proportions of copy number variation categories across all tumor samples (*left panel)* and matching PDTOs (*right panel*).

**Supplementary Figure 3.** Circos plots for three tumor/PDTO pairs exemplifying the concordance of chromosomal structural rearrangements between PDTO and parent tumors.

**Supplementary Figure 4. Mutational Landscape of Samples - Commonly Reported Mutations in Colorectal Cancer. a)** Oncoprint showing genes commonly known to undergo alterations in microsatellite stable colorectal cancer across our whole cohort of samples (tumors and PDTOs). **b)** Oncoprint showing genes commonly known to undergo alterations in microsatellite stable colorectal cancer across all collected colorectal liver metastases (n = 40). Mutation consequences and mutation type (Single Nucleotide Variants - SNV, Insertions – INS, Deletions – DEL, Multi-hits – MHT and Substitutions – SUB) are indicated.

**Supplementary Figure 5. Organoids Maturation and Determination of FOLFOX sensitivity. a)** Bright field and fluorescence imaging showing cellular heterogeneity of PDTOs following lentiviral transduction with 6 fluorescent LeGO (Lentiviral Gene Ontology) vectors (as per Mohme M et al., Mol Ther. 2017 Mar 1;25(3):621-633.)(x4 magnification). **b)** Bright field imaging showing progressive maturation of PDTOs from immature cystic organoid to complex, branching PDTO at day 9, typical of mature PDTOs used for the cytotoxicity assays. **C)** Representative dose response curves depicting the response of 7 PDTOs to FOLFOX, highlighting differences between very sensitive (P520_LT2 and P190_LT) and resistant (P315_LT and P275_LT, red curves) PDTOs, with a range of semi-sensitive ones in between. Stemming from this data, 25uM was selected as treatment dose for molecular studies as it discriminates efficiently the responses from resistant and sensitive organoids.

**Supplementary Figure 6a and b - Morphological differences between resistant and sensitive PDTOs after exposure to FOLFOX.** Bright field imaging showing the impact of a 72-hour treatment with 25uM FOLFOX on PDTO morphology depending on whether organoids are resistant, semi-sensitive or sensitive as indicated, compared with their morphology when treated with vehicle.

**Supplementary Figure 7- PDTO RNAseq Analysis. a)** Heatmap summarizing the number and directionality of Differentially Expressed genes across all tested differential expression analyses performed on our PDTO RNAseq data. Number of DEGs for each analysis is shown in brackets next to the analysis name. For each comparative analysis, upregulated genes are indicated in yellow, down-regulated genes in dark blue. Other genes are not DEGs for that analysis.
*Comparisons between vehicle-treated (control) organoids:* Ctrl_Res_Sen = DEGs between vehicle treated Resistant and Sensitive organoids; Ctrl_Res_SemiSen = DEGs between vehicle-treated Resistant and Semi-sensitive organoids; Ctrl_Sen_SemiSen = DEGs between vehicle-treated Sensitive and Semi-sensitive organoids; Ctrl_Res_SenSemiSen = DEGs between vehicle-treated Resistant and (Sensitive + Semi-sensitive) organoids; Ctrl_ResSemiSen_Sen = DEGs between vehicle-treated (Resistant + Semi-sensitive) and Sensitive organoids.
*Comparisons between FOLFOX-treated organoids:* FF_Res_Sen = DEGs between FOLFOX-treated Resistant and Sensitive organoids; FF_Res_SemiSen = DEGs between FOLFOX-treated Resistant and Semi-sensitive organoids; FF_Sen_SemiSen = DEGs between FOLFOX-treated Sensitive and Semi-sensitive organoids; FF_Res_SenSemiSen = DEGs between FOLFOX-treated Resistant and (Sensitive + Semi-sensitive) organoids; FF_ResSemiSen_Sen = DEGs between FOLFOX-treated (Resistant + Semi-sensitive) and Sensitive organoids.
*Comparisons between treatment conditions for each organoid subgroup:* Res_FF_Ctrl = DEGs between FOLFOX-treated and vehicle-treated Resistant organoids**;** Sen_FF_Ctrl = DEGs between FOLFOX-treated and vehicle-treated Sensitive organoids; SemiSen_FF_Ctrl = DEGs between FOLFOX-treated and vehicle-treated Semi-sensitive organoids**;** ResSemiSen_FF_Ctrl = DEGs between FOLFOX-treated and vehicle-treated (Resistant + Semi-sensitive) organoids**;** SenSemiSen_FF_Ctrl = DEGs between FOLFOX-treated and vehicle-treated (Sensitive + Semi-sensitive) organoids**;** treatment_Res_Sen = DEGs between FOLFOX-treated and vehicle-treated organoids in Resistant vs Sensitive organoids [i.e. (FF_Res - Ctrl_Res) vs (FF_Sen - Ctrl_Sen)].
**b)** Heatmap showing significant Hallmark gene-sets identified for each of the comparisons above, as well as whether these genesets are up or downregulated in each comparative analysis **c)** Heatmap showing DEGs between the most FOLFOX-resistant and sensitive PDTOs, highlighting the up-regulation of E2F targets and of genes involved in the G2M checkpoints and mitotic spindle.

**Supplementary Figure 8. RNAseq Analysis of Tumors and Immune Profile. a**) Heatmap summarizing the number and directionality of Differentially Expressed genes across all tested differential expression analyses performed on our FOLFOX-treated tumor sample RNAseq data. Number of DEGs for each analysis is shown in brackets next to the analysis name. For each comparative analysis, upregulated genes are indicated in yellow, down-regulated genes in dark blue. Other genes are not DEGs for that analysis. PD.PR_CR, DEGs between tumors from patients with Progressive Disease or Partial Response and those with Complete Response under neoadjuvant FOLFOX; PD_CR.PR, DEGs between tumors from patients with Progressive Disease and those with Complete or Partial Response under neoadjuvant FOLFOX; PD_PR, DEGs between tumors from patients with progressive disease and those with Partial Response under neoadjuvant FOLFOX; PD_CR, DEGs between tumors from patients with Progressive Disease and those with Complete Response under neoadjuvant FOLFOX. **b)** Heatmap showing significant Hallmark gene-sets (obtained using camera gene set testing of whole transcriptomic data) that differentiate tumor samples treated with neo-adjuvant FOLFOX for each of the sensitivity group comparisons **c)** Heatmap showing E2F, immune and stromal genes that are differentially expressed between tumors with progressive disease (PD) and those that show a complete response (CR). **d)** Boxplots showing immune related signature scores in patient samples treated with neoadjuvant FOLFOX, stratified by response outcome. Tumors that showed a partial or complete response in patients following treatment with FOLFOX had a higher expression of immune related signature scores compared with those tumors that progressed.

**Supplementary Figure 9. Oncoprint illustrating mutation detected in genes associated with DNA Damage Repair** in PDTOs treated with FOLFOX in vitro (**a**) and in samples from patients treated with neoadjuvant FOLFOX (**b**). Mutation types and consequences are indicated as described in the legend for Suppl Figure 4.

**Supplementary Figure 10. Categories of PDTOs Responses detected via imaging in the large kinase inhibitor screen.** Representative Bright Field (BF) and 7AAD images taken prior to (D0) and after (D3) drug treatment. Phenotypic categories derived from analysis of these images: AliveG = alive with growth, AliveC = alive but with no size increase between Day 0 and Day 3, DeathG = where the size of organoids has increased before their death, and Death C = death with reduced size. Magnification bar = 500 um.

**Supplementary Figure 11. Summary of Primary and Secondary PDTO Kinase Screen. a)** Heatmap summarizing results from the primary kinase inhibitor screening assay performed on PDTOs generated from 3 independent patient tumors that progressed under neoadjuvant FOLFOX treatment (yP295_LT1, yP520_LT1, yP516_LT1), showing the direct comparison of results among the 3 patients after treatment with 0.5uM (*left panel*) or 5uM (*right panel*) of each drug**. b)** Graph summarizing synergy scores calculated using the Lowe Reference Model across 4 PDTO lines. Starting from the left of each graph, the lower line represents the average synergy score across the dose-response grid while the upper line indicates the maximum score that can be reached for any given dose-combination across the grid, highlighting the presence of concentration windows for which strong synergy is detected between FOLFOX and inhibitors of cell cycle progression.

**Supplementary Figure 12a, b, c. Bright Field and 7-AAD images from Organoid Screen Showing Response of 3 PDTOs to Chk1 Inhibitors.** Representative images taken prior to (D0, bright field, left column) and after treatment (D3, bright field only, middle column and bright field + 7-AAD, right column) with 250uM FOLFOX (positive control), DMSO (negative control) or 5uM AZ7762, CHIR-124 or LY2603618 as indicated. Images were taken from 3 independent PDTOs, yP516_LT1 (**a**), yP520_LT1 (**b**) and yP295_LT (**c**). Magnification bar = 500 um.

**Supplementary Figure 13. Synergy Analysis and Bright Field images PDTOs treated with FOLFOX combined to CHK1 or WEE1 inhibitors.**
**a**) Example 3D plots and Average Synergy scores in FOLFOX-resistant PDTOs from two independent patients (P275_LT, top panels and yP516_LT, bottom panels), calculated using the BLISS model from the quantification of CTG data after treatment with increasing concentrations of FOLFOX and/or of the CHK1 inhibitor CCT245747, illustrating the synergy between these compounds. **b**) Representative bright field images illustrating the strongly reduced cell viability of FOLFOX-resistant PDTOs following a 72-hour treatment with increasing concentrations of FOLFOX and of the CHK1 inhibitor CCT-245747.

**c**) Example 3D plots and Average Synergy scores in FOLFOX-resistant P275_LT (top panels) and yP516_LT (bottom panels) PDTOs, calculated using the BLISS model from the quantification of CTG data after treatment with increasing concentrations of FOLFOX and/or of the WEE1 inhibitor adavosertib, illustrating the synergy between these compounds. **d**) Representative bright field images illustrating the strongly reduced cell viability of FOLFOX-resistant PDTOs following a 72-hour treatment with increasing concentrations of FOLFOX and of the WEE1 inhibitor adavosertib. **e**) Representative bright field images illustrating the strongly reduced cell viability of FOLFOX-resistant PDTOs following a 72-hour treatment with 10uM FOLFOX, followed by treatment with the MPS1 inhibitor empesertib (5uM) for a further 72h. Magnification bar (panel b) for all images = 500 um.

**Supplementary Figure 14. Schematic diagram representing the proposed impact of targeting CHK1, WEE1, and MPS1-controlled checkpoints in sensitizing resistant tumors to FOLFOX.** Upon FOLFOX exposure, resistant samples arrest their cell cycle during the transition to S-phase and undergo significant DNA damage. Our results unravel their dependance on CHK1 and WEE1 during that phase, with inhibitors targeting these kinases synergizing strongly with FOLFOX to resensitize and exert a strong cytotoxic effect on PDTOs. In FOLFOX-only treated samples, PDTOs move to a G2/M block following FOLFOX withdrawal, while the majority of tumor cells repair their DNA. This step is strongly dependent on the activity of the Spindle Assembly Checkpoint master regulator MPS1, and MPS1 inhibition during that FOLFOX withdrawal phase has a very powerful cytotoxic effect on otherwise FOLFOX-resistant PDTOs.

**Supplementary Figure 15: Plot summarizing the comparison of PDTO viability after treatment with chemotherapy controls in the primary kinase inhibitor screening assay.** Data for CTG and imaging readout is summarized for PDTOs exposed to DMSO or to increasing concentrations of FOLFOX, FOLFIRI or FOLFOXIRI across all assay plates. The values shown are cell counts normalized to the DMSO mean on a per-plate basis. The notch represents the 95% confidence interval around the median (based on the median +/- 1.58*(inter-quartile range)/sqrt(n)).

**Supplementary Figure 16. Relative-log expression (RLE) plots of the RNA-seq data** of the 107 organoid samples treated with FOLFOX or vehicle, colored according to sequencing run (a), patient ID (b), and treatment groups (c); RLE plots of the tumor samples from patients treated by FOLFOX colored according to patient ID and response group are presented in (d) and (e), respectively.

**Supplementary Figure 17. PCA plots of the RNA-seq data** of the 107 organoid samples treated with FOLFOX or vehicle, colored according to sequencing run (a), patient ID (b), and treatment groups (c); PCA plots of the tumor samples from patients treated with FOLFOX colored according to patient ID and response group are presented in (d) and (e), respectively.
