## Supplemental Table 9 for "Targeting of TP53-independent cell cycle checkpoints overcomes FOLFOX resistance in Metastatic Colorectal Cancer"

| **Compound** | **Company/Manufacturer** | **Final Concentration** |
| --- | --- | --- |
| Advanced Dulbecco’s Modified Eagle Medium(DMEM)/F12 | Thermo Fisher Scientific, Waltham, MA, USA (Cat#12634010) | NA |
| Penicillin/Streptomycin | Life Technologies, Carlsbad, CA, USA (Cat#151540-122) | 250 IU/ml |
| A8301(ALK inhibitor) | Merck, Darmstadt, Germany  (Cat#SML0788) | 500 nM |
| B27 Supplement | Life Technologies, Carlsbad, CA,  USA (Cat#17504044) | 40 ul/ml |
| Epidermal growth factor (human, recombinant) | Miltenyi Biotec, Bergisch Gladbach, Germany (Cat#130-097-749) | 50 ng/ml |
| Leu[15] Gastrin I (human) | Merck, Darmstadt, Germany  (Cat#G9145) | 20 ng/mL |
| N-acetyl cysteine | Merck, Darmstadt, Germany  (Cat#A9165) | 1 mM |
| StemMACS^TM^ SB202190 (p38/MAPK inhibitor) | Miltenyi Biotec, Bergisch Gladbach, Germany (Cat#130-106-275) | 10 nM |
| YP-27632 (ROCK inhibitor) | Abcam, Cambridge, UK  (Cat # 120129) | 10 uM |
| Glutamax | Life Technologies, Carlsbad, CA,  USA (Cat#35050-061) | 5 mM |
| SB431542^a^ (ALK inhibitor) | Tocris, Abingdon, UK  (Cat #S4317) | 10 nM |

**Supplementary Table 9 – Organoid Media**

**^a^** SB431542 used for initial establishment of organoids only, not during passaging
