## Supplemental Table 11 for "Targeting of TP53-independent cell cycle checkpoints overcomes FOLFOX resistance in Metastatic Colorectal Cancer"

| **Method/package** | **version** | **ref** |
| --- | --- | --- |
| HISAT2 | 2.0.4 | <https://www.nature.com/articles/s41587-019-0201-4> |
| Rsubread | 3.8 | <https://academic.oup.com/nar/article/47/8/e47/5345150> |
| edgeR | 3.28.0 | <https://academic.oup.com/bioinformatics/article/26/1/139/182458> |
| Limma | 3.42.0 | <https://academic.oup.com/nar/article/43/7/e47/2414268> |
| voom | - | <https://genomebiology.biomedcentral.com/articles/10.1186/gb-2014-15-2-r29> |
| MSigdf | 7.0 | Github: toledoem/msigd |
| singscore | 1.6.0 | <https://bmcbioinformatics.biomedcentral.com/articles/10.1186/s12859-018-2435-4> |
| Cutadapt | 1.9 (WGS) 1.9.1 (WES) | <http://journal.embnet.org/index.php/embnetjournal/article/view/200> |
| BWA-MEM | 0.7.12 (WGS)  0.7.17 (WES) | <https://arxiv.org/abs/1303.3997> |
| Seqliner (WES) | 0.8.2 | seqliner.org |
| samtools | 1.1 (WGS)   1.8 (WES) | <https://pubmed.ncbi.nlm.nih.gov/19505943/> |
| Picard | 1.129 (WGS)  2.17.3 (WES) | <https://broadinstitute.github.io/picard/> |
| qSNP | 2.0 | <https://journals.plos.org/plosone/article?id=10.1371/journal.pone.0074380> |
| GATK HaplotypeCaller | 3.3-0 (WGS)  3.8 (WES) | <https://pubmed.ncbi.nlm.nih.gov/20644199/>  <https://www.ncbi.nlm.nih.gov/pmc/articles/PMC3083463/>  <https://pubmed.ncbi.nlm.nih.gov/25431634/> |
| ascatNgs | 4.0.1 | <https://pubmed.ncbi.nlm.nih.gov/27930809/> |
| Circos | 0.68-1 | <https://www.ncbi.nlm.nih.gov/pmc/articles/PMC2752132/> |
| VarDict | 1.4.6 | <https://pubmed.ncbi.nlm.nih.gov/27060149/> |
| MuTect | 1.1.7 | <https://pubmed.ncbi.nlm.nih.gov/23396013/> |
| Mutect2 | 3.8 | <https://pubmed.ncbi.nlm.nih.gov/20644199/>  <https://www.ncbi.nlm.nih.gov/pmc/articles/PMC3083463/>  <https://pubmed.ncbi.nlm.nih.gov/25431634/> |
| FACETS | 0.5.6 | <https://www.ncbi.nlm.nih.gov/pmc/articles/PMC5027494/> |
| rock | 0.0.17 | Github: pdiakumis/rock |
| tidyverse | 1.3.0 | <https://tidyverse.tidyverse.org/articles/paper.html> |
| complexHeatmap | 2.2.0 | <https://pubmed.ncbi.nlm.nih.gov/27207943/> |
| synergyfinder | 2.0.11 | <https://link.springer.com/protocol/10.1007%2F978-1-4939-7493-1_17> |
